# Interpreting population and family-based genome-wide association studies in the presence of confounding

**DOI:** 10.1101/2023.02.26.530052

**Authors:** Carl Veller, Graham Coop

**Affiliations:** Department of Evolution and Ecology, and Center for Population Biology, University of California, Davis, CA 95616

## Abstract

A central aim of genome-wide association studies (GWASs) is to estimate direct genetic effects: the causal effects on an individual’s phenotype of the alleles that they carry. However, estimates of direct effects can be subject to genetic and environmental confounding, and can also absorb the ‘indirect’ genetic effects of relatives’ genotypes. Recently, an important development in controlling for these confounds has been the use of within-family GWASs, which, because of the randomness of Mendelian segregation within pedigrees, are often interpreted as producing unbiased estimates of direct effects. Here, we present a general theoretical analysis of the influence of confounding in standard population-based and within-family GWASs. We show that, contrary to common interpretation, family-based estimates of direct effects can be biased by genetic confounding. In humans, such biases will often be small per-locus, but can be compounded when effect size estimates are used in polygenic scores. We illustrate the influence of genetic confounding on population- and family-based estimates of direct effects using models of assortative mating, population stratification, and stabilizing selection on GWAS traits. We further show how family-based estimates of indirect genetic effects, based on comparisons of parentally transmitted and untransmitted alleles, can suffer substantial genetic confounding. In addition to known biases that can arise in family-based GWASs when interactions between family members are ignored, we show that biases can also arise from gene-by-environment (G×E) interactions when parental genotypes are not distributed identically across interacting environmental and genetic backgrounds. We conclude that, while family-based studies have placed GWAS estimation on a more rigorous footing, they carry subtle issues of interpretation that arise from confounding and interactions.

## 1 Introduction

Genome-wide association studies (GWASs) have identified thousands of genetic variants that are associated with a wide variety of traits in humans. In the standard ‘population-based’ approach, the GWAS is conducted on a set of ‘unrelated’ individuals. The associations that are detected can arise when a variant causally affects the trait or when it is in tight physical linkage with causal variants nearby.

Central to the aims of GWASs is the estimation of variants’ effect sizes on traits of interest. These effect size estimates are important for identifying and prioritizing variants and implicated genes for functional followup, and may be used to form statistical predictors of trait values or to understand the causal or mechanistic role of genetic variation in traits. Understanding sources of error and bias in GWAS effect size estimates is therefore crucial.

The interpretation of GWAS effect size estimates is complicated by four broad factors (Vilhjálmsson and Nordborg 2013; Young et al. 2019). First, the causal pathways from an allele to phenotypic variation need not reside in the individuals who enrolled in the GWAS, but can also reflect causal effects on the individual’s environment of the genotypes of their siblings, parents, other ancestors, and neighbors (indirect genetic effects, or dynastic effects; Wolf et al. 1998). Second, a phenotypic association can result from correlations between the allele and environmental causes of trait variation (environmental confounding; Lander and Schork 1994). Third, a phenotypic association can be generated at a locus if it is genetically correlated with causal loci outside of its immediate genomic region (genetic confounding; Vilhjálmsson and Nordborg 2013). Fourth, an allele’s effect on a trait might depend on the environment and the allele’s genetic background (gene-by-environment and gene-by-gene interactions, or G×E and G×G; Freeman 1973; Marchini et al. 2005; Gauderman et al. 2017).

Since our primary interest here will be genetic confounding, we briefly describe some potential sources of the long-range allelic associations that drive it: population structure, assortative mating, and selection on the GWAS trait.

Population structure leads to genetic correlations across the genome when allele frequencies differ across populations or geographic regions: sampled individuals from particular populations are likely to carry, across their genomes, alleles that are common in those populations, which induces correlations among these alleles, potentially across large genomic distances. Such genetic correlations persist even after the populations mix, as alleles that were more common in a particular source population retain their association until uncoupled by recombination.

Assortative mating brings alleles with the same directional effect on a trait (or on multiple traits, in the case of cross-trait assortative mating) together in mates, and therefore bundles these alleles in offspring and subsequent generations. This bundling manifests as positive genetic correlations among alleles with the same directional effect (Wright 1921; Crow and Felsenstein 1968), which can confound effect size estimates in a GWAS on the trait.

Finally, natural selection on a GWAS trait can result in genetic correlations by favoring certain combinations of trait-increasing and trait-decreasing alleles. A form of selection that is expected to be common for many traits of interest is stabilizing selection, which penalizes deviations from an optimal trait value. By favoring compensating combinations of trait-increasing and trait-decreasing alleles, stabilizing selection generates negative correlations among alleles with the same directional effect (Bulmer 1971, 1974), and therefore can confound effect size estimates in a GWAS performed on the trait under selection or on a genetically correlated trait.

The potential for dynastic, environmental, and genetic confounds to bias GWAS effect size estimates has long been recognized (Lander and Schork 1994; Ewens and Spielman 1995), and so a major focus of the literature has been to develop methods to control for these confounds (Pritchard and Rosenberg 1999; Price et al. 2010). Standard approaches include using estimates of genetic relatedness as covariates in GWAS regressions (Price et al. 2006; Yang et al. 2014) or downstream analyses such as LD-Score regression (Bulik-Sullivan et al. 2015a,b; Bulik-Sullivan 2015). Such methods aim to control for both environmental and genetic confounding, but do so imperfectly (e.g., Berg et al. 2019; Sohail et al. 2019). Further, it is often unclear what features of genetic stratification are being addressed (Vilhjálmsson and Nordborg 2013; Young et al. 2019): assortative mating in particular may not be well accounted for by these methods (Border et al. 2022b). Moreover, in reality, there is no bright line separating dynastic, environmental, and genetic confounding.

One promising way forward is to estimate allelic effects within families, either by comparing the separate associations of parentally transmitted and untransmitted alleles with trait values in the offspring (Spielman et al. 1993; Allison 1997; Eaves et al. 2014; Weiner et al. 2017; Kong et al. 2018), or by associating differences in siblings’ trait values with differences in the alleles they inherited from their parents (Abecasis et al. 2000; Visscher et al. 2006; Lee et al. 2018). The idea is that, by controlling for parental genotypes, within-family association studies control for both environmental stratification and indirect/dynastic effects, while Mendelian segregation randomizes alleles across genetic backgrounds. In principle, this allows the ‘direct genetic effect’ of an allele—the causal effect of an allele carried by an individual on their trait value—to be estimated. Recognizing that a variant detected in a GWAS will usually not itself be causal for the trait variation but instead will only be correlated with true causal variants, the direct effect of a genotyped variant is usually interpreted as reflecting the direct causal effects of nearby loci that are genetically correlated with the focal locus (Young et al. 2019)—but not the effects of more distant loci that might also be genetically correlated with the focal genotyped locus (e.g., because of population structure or assortative mating).

Consistent with both the presence of substantial confounds in some population-based GWASs and the mitigation of these confounds in within-family GWASs, family-based estimates of direct effect sizes and aggregate quantities based on these estimates (e.g., SNP-based heritabilities) are substantially smaller than population GWAS estimates for a number of traits, most notably social and behavioural traits (Lee et al. 2018; Selzam et al. 2019; Mostafavi et al. 2020; Howe et al. 2022; Young et al. 2022). Likewise, estimates of genetic correlations between traits are sometimes substantially reduced when calculated using direct effect estimates from within-family GWASs (e.g. Howe et al. 2022). While some of these findings could reflect the contribution of indirect genetic effects to population GWASs, it is also likely that, at least for some traits, standard controls for population stratification in population GWASs have been insufficient (Berg et al. 2019; Sohail et al. 2019; Young et al. 2022; Okbay et al. 2022; Nivard et al. 2022; Border et al. 2022a).

Our aim in this paper is to study a general model of confounding in GWASs, to generate clear intuition for its influence on estimates of effect sizes in both population- and family-based designs. A number of the issues that we analyze have previously been raised, particularly in the context of population-based GWASs (e.g., Rosenberg and Nordborg 2006; Platt et al. 2010; Vilhjálmsson and Nordborg 2013; Young et al. 2019); here, we analyze them in a common framework that allows for comparison of multiple sources of confounding in both population and family-based GWASs. There is a large literature on GWASs in non-human organisms (e.g., Atwell et al. 2010; Hayes and Goddard 2010; Peiffer et al. 2014; Josephs et al. 2017). However, although the results and intuition that we derive here apply equally well to human and non-human GWASs, we shall interpret them primarily from the perspective of human GWASs, in which the inability to experimentally randomize environments, together with the small effects that investigators hope to detect, makes confounding a particular concern.

Our first focus is on confounding—and genetic confounding in particular—in the absence of G×E and G×G interactions. To better understand the differences between population and within-family GWASs, we first study a general model of genetic confounding in the absence of G×E and G×G interactions. We derive expressions for estimators of direct effects in both population and within-family GWASs, as functions of the true direct and indirect effects at a locus and the genetic confounds induced by other loci. In doing so, we find that family-based estimates of direct effects are in fact susceptible to genetic confounding, contrary to standard interpretation. Reassuringly, in many of the models we consider, the resulting biases are likely to be small in humans. We also address a related case: family-based GWAS designs that consider transmitted and untransmitted parental alleles and in which the indirect (or ‘dynastic’) effect of an allele is estimated from its association with the offspring’s phenotype when carried by the parent but not transmitted to the offspring. We show that this estimator of indirect effects can be substantially biased by genetic and environmental confounds, in a similar way to population estimates of direct effects. Next, we consider various sources of genetic confounding—assortative mating, population structure, and stabilizing selection on GWAS traits— and how they influence estimates of direct effects in both population and within-family GWASs.

We then turn to sibling indirect effects, which are known to bias estimates of direct effects in sibling-based GWASs (Young et al. 2019, 2022). We characterize this bias in a simple model, and contrast it to the bias caused by sibling indirect effects in a population GWAS.

Finally, we consider G×E and G×G interactions, showing how their presence can bias population and family-based estimates of direct genetic effects in contrasting ways, complicating the interpretation of family-based estimates.

## 2 Effect size estimates in association study designs

Our primary focus will be on how genetic confounding can bias the estimation of direct genetic effects. These genetic confounds are due to associations between a genotyped variant at a GWAS locus and causal variants at other loci. As we will see, two kinds of association must be distinguished: cis-linkage disequilibrium (cis-LD) and trans-linkage disequilibrium (trans-LD). Genetic variants *A* and *B* are in positive cis-LD if, when an individual inherits *A* from a given parent, the individual is disproportionately likely to inherit *B* from that parent (Fig. 1A). *A* and *B* are in positive trans-LD if, when an individual inherits *A* from one parent, the individual is disproportionately likely to inherit *B* from the other parent (Fig. 1B). These covariances have also been called gametic and non-gametic LD, respectively (e.g. Weir 2008). To quantify the degrees of cis-LD and trans-LD, we denote by *D_ij_* and 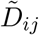 the allelic covariances between focal variants at loci *i* and *j* in cis and in trans, and we denote by *r_ij_* and 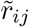 the analogous allelic correlation coefficients. For some of our results, it will be important to distinguish the LD present in the sample on which the association study is performed and the LD present among the parents of the sample.

**Figure 1:**
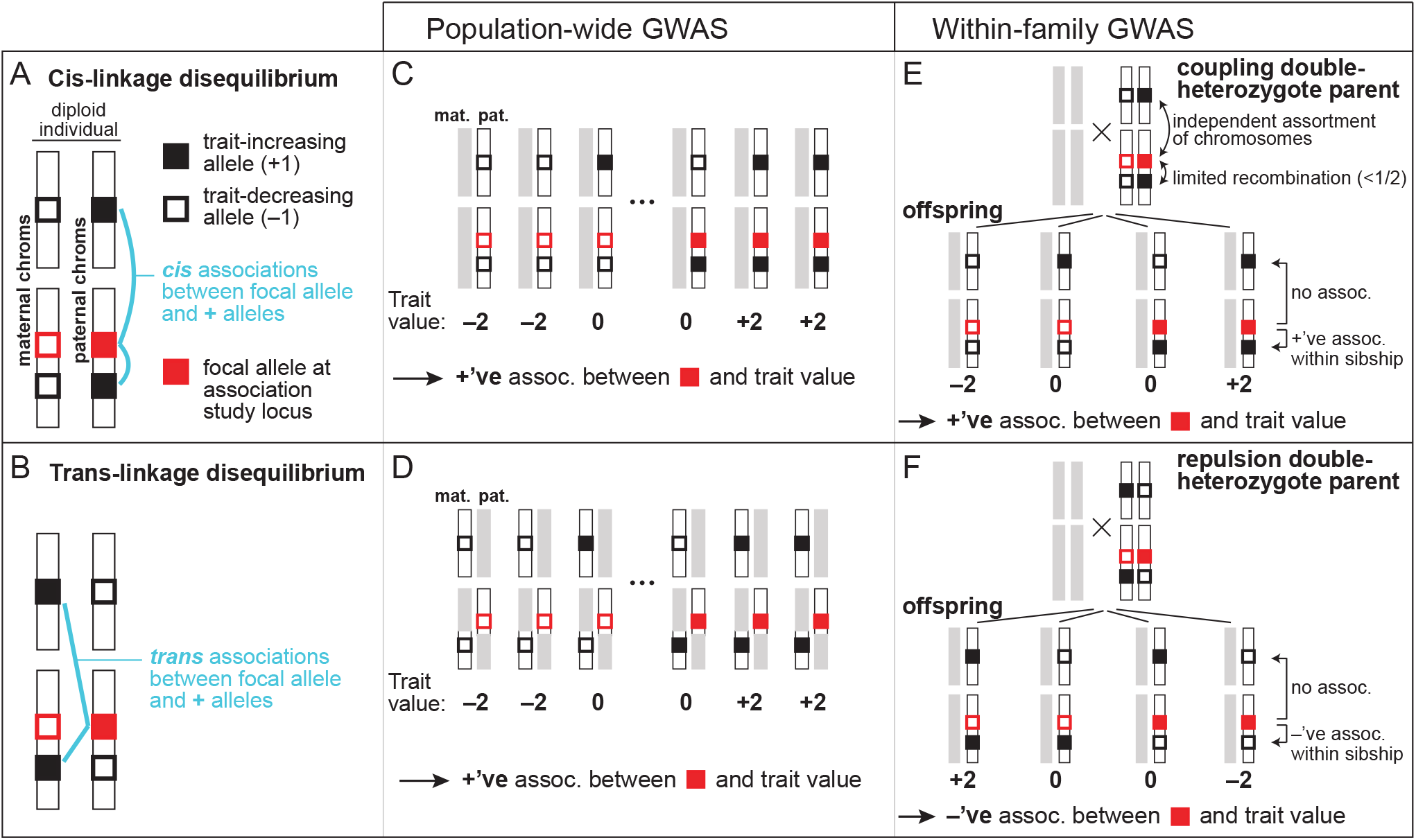
The influence of cis- and trans-LD on effect size estimates in population-based and within-family association studies. (A) The focal allele at an association study locus (solid red square) is in positive cis-LD with trait-increasing alleles at other loci (solid black squares) if it is disproportionately likely to be found alongside them on an individual’s maternally or paternally inherited genome. (B) The focal allele at the study locus is in positive trans-LD with trait-increasing alleles at other loci if it is disproportionately likely to be found across from them on the maternally and paternally inherited genomes. (C,D) In a population association study, both positive cis- and trans-LD between the focal allele at the study locus and trait-increasing alleles anywhere else in the genome—either on the same chromosome as the study locus or on different chromosomes—generate a spuriously high effect size estimate at the study locus. (E,F) In a sibling association study, a trait-increasing allele causes a spuriously increased effect size estimate at the study locus if the parent is a coupling double heterozygote for the focal and trait-increasing alleles, having inherited them from the same parent (E), but a spuriously decreased estimate if the parent is a repulsion double heterozygote, having inherited them from different parents (F). These biases arise only if the trait-affecting locus is on the same chromosome as the focal study locus. The net bias depends on the relative frequencies of coupling and repulsion double heterozygotes in the parents, which depends on the difference in the degrees of cis- and trans-LD.

Consider a trait Y influenced by genetic variants at a set of polymorphic loci L, each of which segregates for two alleles. For ease of interpretation, and without loss of generality, we designate the ‘focal’ allele at locus *l* ∈ *L* to be the allele that directly increases the trait value, and we denote by *p_l_* the frequency of this allele. Allelic effects are assumed to be additive within and across loci, such that the trait value of an individual can be written

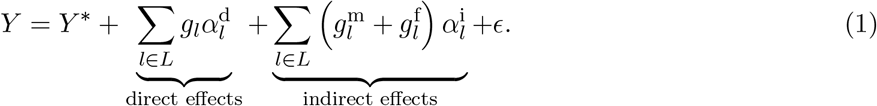

Here, *g_l_*, 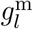, and 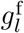 are the numbers (0, 1, or 2) of focal alleles carried at locus *l* by the individual, their mother, and their father respectively, 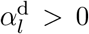 is the direct effect of the focal allele at *l*, and 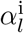 is its indirect effect via the maternal and paternal genotypes. (For simplicity, we assume that indirect effects via the maternal and paternal genotypes are equal; this assumption is relaxed in Appendix A1.) *ϵ* is the environmental noise, with 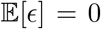, and *Y*\* is the expected trait value of the offspring of parents who carry only trait-decreasing alleles.

### 2.1 Population-based association studies

The variants at a genotyped locus will usually not themselves have causal effects on the trait, but will instead be in cis-LD with—and thus ‘tag’—causal variants at nearby loci. Thus, we typically think of the association at a focal genotyped locus as reflecting the direct contributions of a relatively small number of tightly linked loci, *L*_local_, found within tens or perhaps hundreds of kb from the focal locus (Pritchard and Przeworski 2001). Under the additive model, therefore, the standard interpretation is that a population association study performed at a focal genotyped locus λ provides an estimate of the quantity

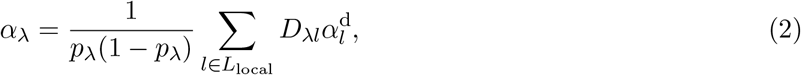

where *p*_λ_ is the frequency of the focal allele at λ, and *D*_λ*l*_ is the degree of cis-LD between the focal allele at λ and a causal allele at a nearby locus *l* ∈ *L*_local_. It is reasonable to think of this quantity as the ‘direct effect’ tagged by the focal variant at the genotyped locus λ: in the absence of confounding, it can be interpreted as the average phenotypic effect of randomly choosing a non-focal allele in the population and swapping it for a focal allele, where in this hypothetical swap, the causal alleles near the locus are included. For concreteness, we assume some fixed *L*_local_ in our analyses, but in practice researchers seldom have a pre-defined number of ‘local’ SNPs in mind.

Effect size estimation in a population GWAS is complicated by the presence of environmental and genetic stratification. Under the model in Eq. (1), if we perform a standard population association study at locus λ, the estimated effect of the focal allele on the trait Y is

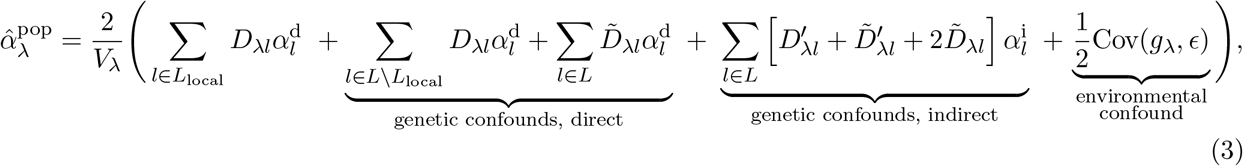

where, of the cis- and trans-LD terms, *D*_λ*l*_ and 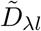 are defined in the GWAS sample while 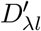 and 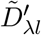 are defined in their parents (Appendix A1.2). *V*_λ_ is the genotypic variance at λ, equal to 2*p*_λ_(1 – *p*_λ_)(1 + *F*_λ_) where *F*_λ_ is Wright’s coefficient of inbreeding at λ.

The environmental confound is Cov(*g*_λ_, *ϵ*)/*V*_λ_; all non-local cis- and trans-LD terms in the study sample (*D*_λ*l*_ and 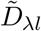, *l* ∉ *L*_local_) are direct genetic confounds (Fig. 1C,D); and all cis- and trans-LD terms among parents of sampled individuals (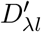 and 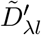), together with all trans-LD terms in the study sample 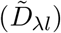, are indirect genetic confounds.

The direct genetic confounds arise because an allele carried by an offspring at λ is correlated with the alleles that they carry at other loci *l* ∈ *L* (via *D*_λ*l*_ and 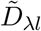) that directly affect the trait value. The indirect genetic confounds arise because an allele carried by the offspring at λ—say, the maternal allele—is correlated with alleles carried by the offspring’s mother at other loci (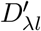 and 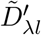) and alleles carried by their father (as reflected by the trans-LD in the offspring, 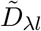). These alleles in the parents can indirectly affect the offspring’s trait value.

Thus, as is now well appreciated, population-based GWASs potentially suffer from many types of confounds (Vilhjalmsson and Nordborg 2013; Young et al. 2019). In practice, they can be reduced by including principal components—which capture genome-wide relatedness among GWAS participants—as regressors in a GWAS, or by using relatedness matrices in mixed models (Price et al. 2006; Yang et al. 2014). However, it is often unclear exactly what these methods control for in a given application (Vilhjálmsson and Nordborg 2013; Young et al. 2019), and they have been shown to be inadequate in important cases (e.g., Berg et al. 2019; Sohail et al. 2019). When principal components (or other controls) fail to account fully for stratification, then Eq. (3) can be interpreted as a decomposition of the remaining, uncontrolled-for confounding in the GWAS.^1^

### 2.2 Within-family association studies

The two within-family association study designs that we consider are parent-offspring GWASs and sibling GWASs. Other designs have been proposed to control for genetic and environmental confounding in the estimation of aggregate quantities such as heritability (e.g., Young et al. 2018a), but our primary focus is on the estimation of single-marker effect sizes. We do later turn to the interpretation of polygenic score regressions within families.

#### Estimates of direct genetic effects

Parent-offspring studies can be used to estimate trait associations separately for parentally transmitted and untransmitted variants at a locus λ, 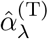 and 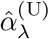, by regressing the trait value *Y* on the transmitted and untransmitted genotypes, 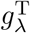 and 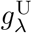 (Kong et al. 2018). The aim is often to estimate the direct effect of a variant, 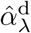, as the difference between these two estimates:

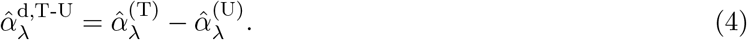

A second aim is to treat 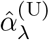 as an estimate of the indirect, or family, effect of the variant. We return to this second aim later.

In Appendix A1.4, we show that, in the absence of interactions between parental and offspring genotypes, the estimate of the direct effect of a variant at locus λ in a parent-offspring study is

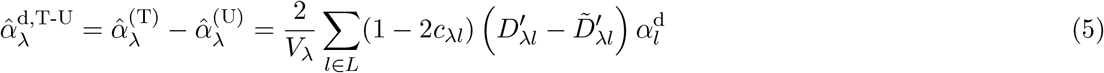

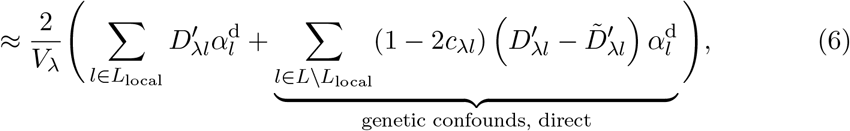

where *c*_λ*l*_ is the sex-averaged recombination rate between λ and *l*. The cis- and trans-LD terms 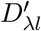 and 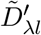 are measured in the parents.

Similarly, an estimate of the direct effect can be obtained from pairs of siblings by regressing the differences in their phenotypes on the differences in their genotypes at the focal locus λ. In the presence of genetic confounds, this procedure yields a similar estimate to Eq. (6):

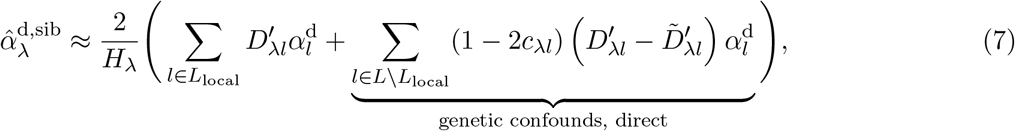

where *H*_λ_ is the fraction of parents who are heterozygous at locus λ (Appendix A1.3). An assumption in sibling GWASs is that an offspring’s phenotype is not influenced by the genotypes of their siblings—i.e., that there are no sibling indirect genetic effects. We consider violations of this assumption later.

In Eqs. (6) and (7), there is no environmental confound, because family-based GWASs successfully randomize the environments of family members with respect to within-family genetic transmission.

The derivations above further show that, while population association studies are biased by sums of trans- and cis-LD between the focal locus and all causal loci (Eq. 3), within-family association studies are instead biased by *differences* between trans- and cis-LD, and moreover, that the biases in within-family studies are driven only by LD between the focal locus and causal loci on the same chromosome (*c*_λ*l*_ < 1/2). To provide an intuition for this result, we focus our discussion on a sibling association study performed at λ; the intuition is identical for the analogous parent-offspring study.

Because the difference between two siblings in their maternally inherited genotypes is independent of the difference in their paternally inherited genotypes, we may consider maternal and paternal transmissions separately in studying how a locus *l* ∈ *L* can confound effect size estimation at λ in a sibling association study. We will phrase our discussion in terms of maternal transmission.

For effect size estimation at λ to be genetically confounded by maternal transmission at a distant locus *l*, the mother must be heterozygous at both loci. For if she were homozygous at *l*, then maternal transmission at *l* could not contribute to any trait differences between her offspring, while if she were homozygous at λ, maternal transmission would not result in genetic variation among her offspring at λ with which trait variation could be associated. Therefore, we restrict our focus to mothers who are heterozygous at both λ and *l*, or ‘double heterozygotes’. Two kinds exist (Fig. 1E,F): coupling double heterozygotes who carry the focal alleles at λ and *l* on the same haploid genome (‘in cis’), and repulsion double heterozygotes who carry them on separate haploid genomes (‘in trans’).

We first consider the case where the recombination rate between λ and *l* is small (*c*_λ*l*_ ≪ 1/2). In this case, if the mother is a coupling double heterozygote, then her offspring will tend to inherit either both or neither of the focal alleles at λ and *l* (Fig. 1E). Therefore, if one sibling inherits the focal allele at λ and another does not, the first sibling will tend to inherit the focal (trait-increasing, as we have defined it) allele at *l* and the second sibling will not, so that the effect of locus *l* positively confounds the association between λ and the trait (Fig. 1E). If the mother is instead a repulsion double heterozygote, then her offspring will tend to inherit either the focal allele at λ or the focal allele at *l*, but not both (Fig. 1F). In this case, if one sibling inherits the focal allele at λ and another does not, the second sibling will tend to inherit the focal (trait-increasing) allele at *l* and the first sibling will not, so that the effect of locus *l* negatively confounds the association between λ and the trait (Fig. 1F). When λ and *l* are linked, therefore, the way in which *l* genetically confounds the effect size estimate at λ depends, positively or negatively, on whether the fraction of coupling double heterozygotes among parents is greater or smaller, respectively, than the fraction of repulsion double heterozygotes.

In contrast, if λ and *l* are unlinked (*c*_λ*l*_ = 1/2), then transmissions from coupling and repulsion double heterozygote parents are equal, and so *l* cannot confound estimates at λ (Fig. 1E,F). Put differently, meiosis in double heterozygotes fully randomizes joint allelic transmissions at λ and *l*, with offspring equally likely to inherit any possible combination of alleles at the two loci.

Therefore, only linked loci *l* can confound a family-based association study at λ, and they do so in proportion to (i) how small the recombination rate between λ and *l* is, and (ii) the difference between the fractions of parents who are coupling and repulsion double heterozygotes at λ and *l*. Accordingly, if we write these fractions of parents as 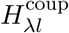 and 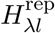, then 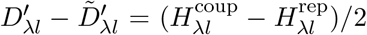, and so Eq. (7) (and Eq. 6) can be rewritten in terms of the relative frequencies of the two kinds of double-heterozygotes:

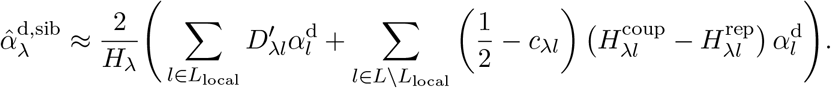

In a species with many chromosomes, such as humans, for a given locus, there will be many more unlinked loci than linked loci. Therefore, the set of loci that can confound a family-based association study at a given locus will be much smaller than the set of loci that can confound a population association study at the locus. It will often be the case, therefore, that biases in the estimation of direct genetic effects will be smaller in family-based studies than in population studies, a point that we explore below when we consider sources of genetic confounding.

#### Estimates of indirect genetic effects

We now return to the regression of the trait on the untransmitted genotype in parent-offspring GWASs, 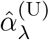, which has sometimes been treated as an estimate of the indirect effect 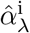. Assuming equal indirect effects via maternal and paternal genotypes (an assumption that we relax in Appendix A1.4),

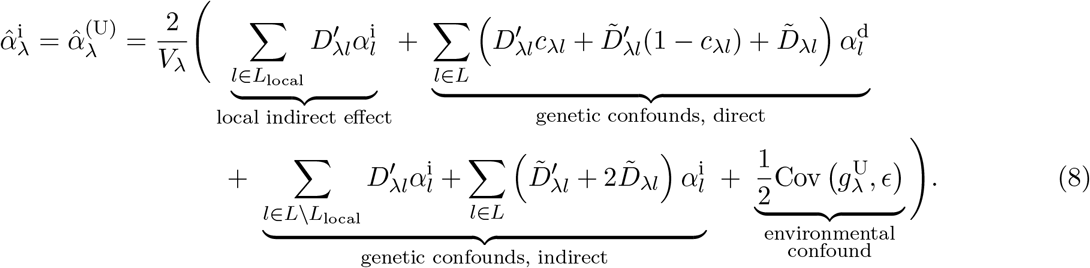

The direct genetic confound reflects associations of the untransmitted alleles at the focal locus with alleles that are transmitted to the offspring at causal loci *l* ∈ *L* and which directly affect the offspring’s trait value (via 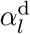). These associations are due to covariances among alleles in each parental genome (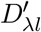 and 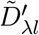) and across the parental genomes (reflected as trans-LD in the offspring, 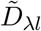). The indirect genetic confound reflects associations of the untransmitted alleles to alleles at other loci in the parents, which can indirectly affect the offspring trait value (via 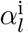). Finally, unlike in family-based estimates of direct genetic effects (Eqs. 6 and 7), family-based estimates of indirect effects suffer from environmental confounding, in the same way that population GWASs do (Eq. 3).

Therefore, estimating the indirect effect by regressing the trait value on the untransmitted genotype is highly susceptible to environmental confounding as well as both direct and indirect genetic confounding, in a similar way to estimating the direct effect via a population-based association study (Shen and Feldman 2020). Adjustments for assortative mating in particular have been included in some PGS-based analyses of indirect effects (e.g., Kong et al. 2018; Young et al. 2022). However, it is not clear how robust these adjustments are in the presence of multiple forms of confounding.

### 2.3 Polygenic scores and their phenotypic associations

A current drawback to family-based GWASs is that sample sizes are often small, limiting power to estimate direct genetic effects. Because of this limitation, instead of estimating per-locus effect sizes in family designs, investigators often measure the within-family phenotypic association of a combined linear predictor, a polygenic score (PGS), constructed using effect size estimates across many loci from a population GWAS. In the sibling-based version of this study design, the difference in siblings’ population-based PGSs is regressed on their difference in phenotypes (e.g., Lee et al. 2018; Selzam et al. 2019). In parent-offspring designs, the population-based PGSs constructed separately for transmitted and untransmitted alleles are used as linear predictors of the offspring’s phenotype, and the difference in their slopes in this regression is estimated (e.g., Kong et al. 2018; Okbay et al. 2022).

When such PGS regressions are used within families for the same phenotype as the population GWAS, a non-zero slope of the PGS is usually interpreted as reflecting the fact that the PGS—despite having been calculated from a population GWAS and therefore subject to many potential confounds—nevertheless does capture the direct genetic effects of alleles. When the PGS for one phenotype is regressed within families on the value of another phenotype, non-zero slopes are often interpreted as evidence that direct genetic effects on the two phenotypes are causally related, for example through pleiotropic effects of the alleles involved.

Suppose that we have performed a population GWAS for trait 1, generating effect size estimates 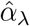 at a set of genotyped loci λ ∈ Λ. To construct a PGS for trait 1, these effect size estimates are used as weights in a linear sum across an individual’s genotype:

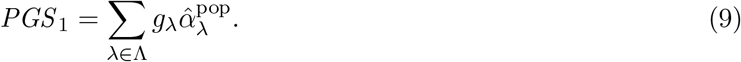

In a sibling-based study (the results and intuition below will be the same for a parent-offspring study), the difference between siblings’ trait-1 PGSs, Δ*PGS*_1_, is regressed on the difference in their values for trait 2, Δ*Y*_2_ (note that trait 2 could be the same as trait 1). If *L* is the set of loci that causally underlie variation in trait 2, and *β_l_* are the true effects of variants at these loci on trait 2, then the numerator of the slope in this regression can be written as

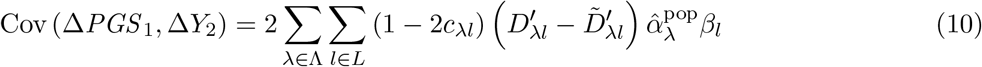

(see Appendix A2). Note that, while the population-based effect size estimates 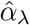 depend on cis- and trans-LD, as detailed by Eq. (3), the patterns of LD may differ from those in the family study (the 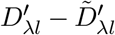 term in Eq. 10) if the population- and family-based studies differ in relevant aspects of sample composition.

The intuition for Eq. (10) is similar to that for the single-locus effect size estimate in a sibling GWAS (Eq. 7). The numerator of the difference in slopes of transmitted and untransmitted PGSs in a parent-offspring design takes a similar form to Eq. (10).

In the absence of confounding and under some simplifying assumptions, the sibling PGS covariance measures the contribution of each locus included in the PGS to the additive genic covariance between traits 1 and 2 that is tagged by the genotyped variants included in the PGS (see Eq. A.23 in Appendix A2). Under these assumptions, the sibling PGS slope therefore does provide a measure of the underlying pleiotropy between the traits.

Interpretation of the sibling PGS slopes is more complicated in the presence of genetic confounding (see Eq. A.22 in Appendix A2), which is absorbed into the effect size estimates 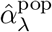 (Eq. 3) so that the PGS applies a potentially strange set of weights to the genotyped loci it includes. (A related problem occurs when indirect genetic effects absorbed by the population-based PGS change the interpretation of within-family PGS slopes—see Trejo and Domingue (2018); Fletcher et al. (2021).) A non-zero sibling PGS slope still establishes that the trait-1 PGS loci are in systematic signed intra-chromosomal LD with loci that causally affect trait 2. However, it no longer necessarily implies that traits 1 and 2 are causally related via pleiotropy, for two reasons. To understand these reasons, suppose that the causal loci for traits 1 and 2 are distinct, i.e., that there is in fact no pleiotropy. First, a SNP included in the trait-1 PGS could tag local variants that causally affect trait 1 but which are also, via sources of confounding such as cross-trait assortative mating, in systematic long-range LD with variants on the same chromosome that causally affect trait 2. Such SNPs will be predictive of sibling differences in trait 2, even though they locally tag only trait-1 causal variants. Second, LD between variants on the same or distinct chromosomes that are causal for trait 1 and trait 2 will cause some SNPs that locally tag trait-2 causal variants to be significantly associated with trait 1 in a population GWAS, and therefore to be included in the trait-1 PGS. These SNPs, since they tag trait-2 causal variants, will be predictive of sibling differences in trait 2.

In summary, in the presence of confounding, non-zero sibling PGS slopes cannot be viewed as de facto evidence for causal relationships between traits.

## 3 Sources of genetic confounding in association studies

As we have seen, genetic confounding of association studies depends, in ways that vary across study designs, on levels of non-local cis- and trans-LD between the study locus and loci that influence the study trait. Below, we consider various processes that give rise to non-local cis- and trans-LD, and their likely impact on the different association study designs. We focus our attention on the potential for these sources of LD to confound measurement of several key metrics. First, the average deviation of the estimated effect size from its true value, 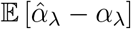. This measure indicates if effect sizes are systematically overestimated or underestimated because of genetic confounding. Second, the average squared effect size estimate, weighted by heterozygosity: 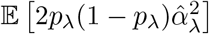. This quantity is related to important measures such as the genetic variance and SNP-based heritability (Bulik-Sullivan et al. 2015b). It is also directly related to the variance of effect size estimates, and therefore captures the additional noise that genetic confounding creates in effect size estimation at a given locus. Finally, if GWASs have been performed on more than one trait, the covariance across loci of the effect size estimates for two traits may be of interest. This covariance is determined by the average heterozygosity-weighted product 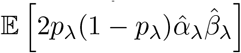, where 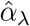 and 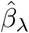 are the effect size estimates at locus λ for traits 1 and 2.

In what follows, for simplicity, we ignore indirect effects and assume that there is no environmental confounding (i.e., no correlation between genotypes and the environmental effects *ϵ*). For each of the sources of genetic confounding that we consider, we calculate the three measures listed above both analytically and in whole-genome simulations carried out in SLiM 4.0 (Haller and Messer 2019). In our simulations, we use two recombination maps: (i) for illustrative purposes, a simple hypothetical map where the genome lies along a single chromosome of length 1 Morgan, and (ii) the human linkage map generated by Kong et al. (2010). A more detailed description of the simulations can be found in the Methods, and code is available at github.com/cveller/confoundedGWAS.

### 3.1 Assortative mating

Assortative mating is the tendency for mating pairs to be correlated for particular traits—either the same trait (same-trait assortative mating) or distinct traits (cross-trait assortative mating). For example, humans are known to exhibit same-trait assortative mating for height and cross-trait assortative mating for educational attainment and height (amongst many other examples, reviewed in Horwitz and Keller 2022; Border et al. 2022a). Assortative mating generates both cis- and trans-LD: It generates positive trans-LD among trait-increasing alleles because genetic correlations between mates translate to genetic correlations between maternally and paternally inherited genomes, and it generates positive cis-LD among trait-increasing alleles because, over generations, recombination converts trans-LD into cis-LD (Crow and Felsenstein 1968). (In some cases, assortative mating can generate cis-LD by mechanisms additional to recombination—see Veller et al. 2020.)

#### Constant-strength assortative mating

If the strength of assortative mating (measured by the phenotypic correlation among mates *ρ*) is constant over time, and there are no other sources of genetic confounding such as population structure, then, for a given pair of loci *l*,*l′* ∈ *L*, the positive cis-LD *D_ll′_*, will initially be smaller than the positive trans-LD 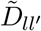, but will gradually grow towards an equilibrium value equal to the trans-LD 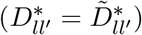; in this equilibrium, assortative mating generates new cis-LD at the same rate as old cis-LD is destroyed by recombination (Crow and Felsenstein 1968, Appendix A3.1).

Therefore, in a population GWAS, effect size estimates will initially be biased upwards because of positive trans-LD, and the magnitude of the bias will grow over time as positive cis-LD too is generated from this trans-LD (Eq. 3; Fig. 2). In contrast, in a family-based GWAS, effect size estimates will initially be biased downwards because the positive trans-LD exceeds the positive cis-LD (Eqs. 6 and 7; Fig. 2). However, as the cis-LD grows over time towards the value of the trans-LD, the magnitude of the downward bias will shrink, and, in equilibrium, the family-based GWAS will not be confounded by assortative mating (Fig. 2).

**Figure 2:**
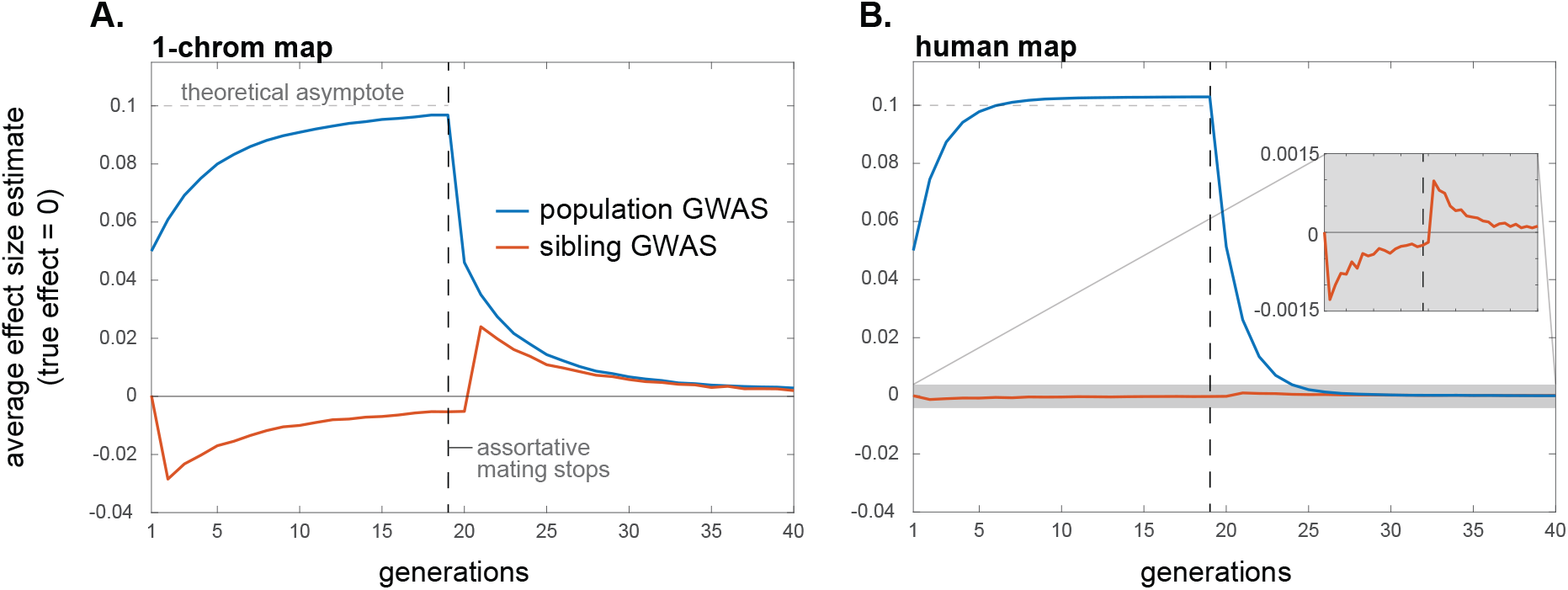
Assortative mating systematically biases effect size estimation in population and within-family GWASs, although the bias in within-family GWASs is expected usually to be small. Here, cross-trait assortative mating between traits 1 and 2 occurs for the first 19 generations, after which mating is random. Assortative mating is sex-asymmetric, with strength *ρ* = 0.2. Distinct sets of loci underlie variation in trait 1 and 2, with effect sizes at causal loci normalized to 1. Plotted are average estimated effects of the focal alleles at loci causal for trait 1 in population and within-family GWASs on trait 2, for a hypothetical genome with one chromosome of length 1 Morgan (A) and for humans (B). Since the traits have distinct genetic bases, the true effects on trait 2 of the alleles at trait-1 loci are zero. The horizontal lines at 0.1 are a theoretical ‘first-order’ approximation of the asymptotic bias in a population GWAS (Appendix A3.1). Profiles are averages across 10,000 replicate simulation trials. Simulation details can be found in the Methods.

Under certain simplifying assumptions, we can calculate the average bias that assortative mating induces in a population GWAS in equilibrium, in the absence of other sources of genetic confounding such as population structure (Appendix A3.1). In the case of same-trait assortative mating, effect size estimates are inflated by an average factor of approximately *h*^2^*ρ*/(1 – *h*^2^*ρ*), where *ρ* is the phenotypic correlation among mates and *h*^2^ is the trait heritability (for similar calculations, see Yengo et al. 2018; Border et al. 2022b). In the case of cross-trait assortative mating, if assortative mating is directional/asymmetric with respect to sex—i.e., the correlation *ρ* is between female trait 1 and male trait 2—then assortative mating generates spurious associations between trait 1 and alleles that affect trait 2 (and vice versa). If the loci underlying the two traits are distinct, then, in equilibrium, the spurious effect size estimate at non-causal loci is approximately *h*^2^*ρ*/2 times the effect at causal loci, assuming the traits to have the same heritabilities and genetic architectures (horizontal dahsed line in Fig. 2). If cross-trait assortative mating is bi-directional/symmetric with respect to sex, then, in equilibrium, the average spurious effect size estimate at non-causal loci is approximately *h*^2^*ρ* times the effect at causal loci. Upward biases in effect size estimates at causal loci are also expected under cross-trait assortative mating, but these are second-order relative to the biases at non-causal loci (Fig. S1).

The systematic over- and under-estimation of effect sizes that assortative mating induces in population and family-based GWASs, respectively, will also affect our second measure of interest, the heterozygosity-weighted average squared effect size estimate 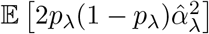 (and therefore also downstream quantities such as SNP heritabilities). In a population GWAS, the presence of trans-LD and the gradual creation of cis-LD under assortative mating will increase the biases in effect size estimates over time (Fig. 2), which will concomitantly increase the average value of 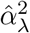 (Fig. 3; also see Border et al. 2022b). Moreover, cross-trait assortative mating will generate signals of genetic correlations among traits even in the absence of any pleiotropic effects of underlying variants (Border et al. 2022a). In a family-based GWAS, the temporary attenuation of effect size estimates owing to a transient excess of trans-LD over cis-LD under assortative mating will lead to a similar attenuation in the average squared effect size estimate (Fig. 3), although, like the bias in effect size estimates themselves, this attenuation is expected to be small in humans (Fig. 3B).

**Figure 3:**
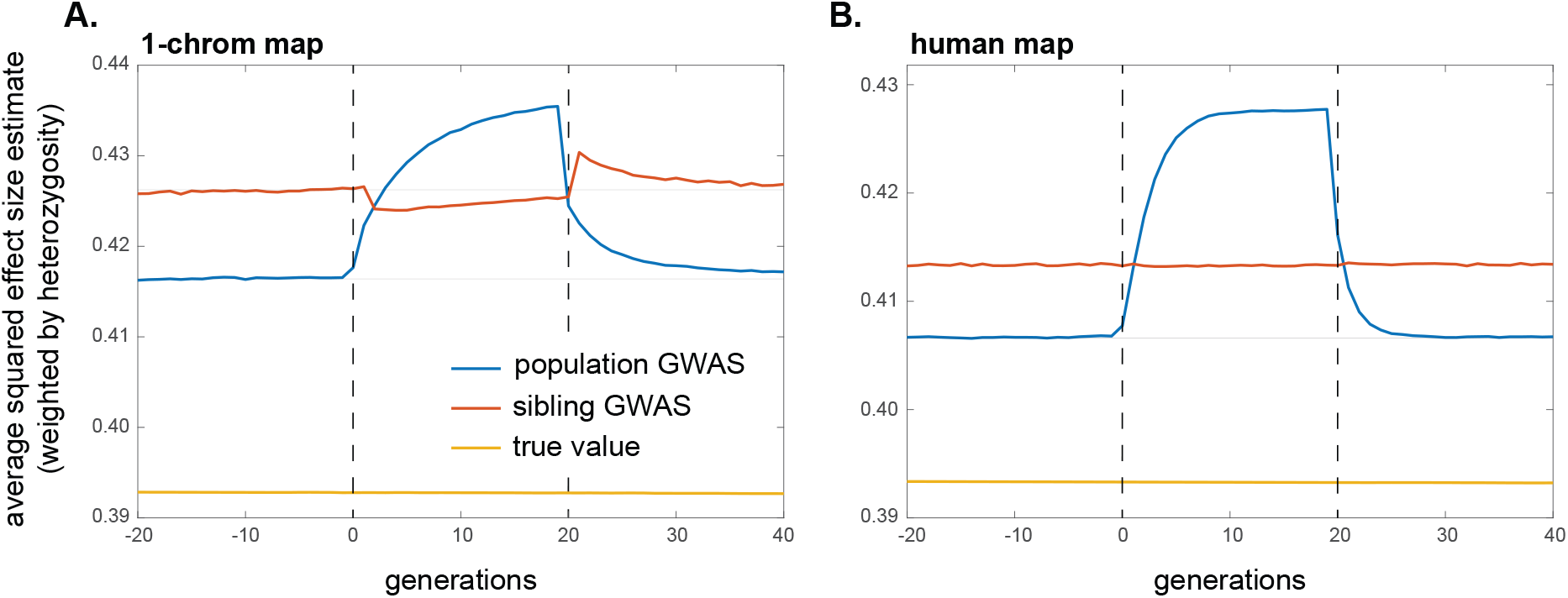
The impact of assortative mating on the average squared effect size estimate in population and within-family GWASs. Same-trait assortative mating of strength *ρ* = 0.2 occurs in generations 0–19; mating is random before and after this period. Under random mating, the average squared effect size estimates exceed the true average squared effect size (yellow line) because random drift generates chance local LD with causal alleles that inflates the variance of effect size estimation (e.g. Bulik-Sullivan et al. 2015b). The magnitude of this variance inflation depends on the GWAS design and sample size, and the effect of assortative mating and its cessation should be judged in reference to it. To guide the eye in this judgment, the faint horizontal lines show the average squared effect size estimate in the last 20 generations of the initial burn-in period of random mating. Profiles are averages across 10,000 replicate simulation trials. Simulation details can be found in the Methods.

As shown by Border et al. (2022a,b), the effects of assortative mating on estimates of heritability and genetic correlations described above are not well controlled for by LD Score regression (Bulik-Sullivan et al. 2015a,b). The LD score of a variant proxies the amount of local causal variation the SNP tags, but because assortative mating generates long-range signed LD among causal variants, it causes local causal variants to be in long-range signed LD with other causal variants throughout the genome. Therefore, the slope of the LD score regression absorbs the effects of assortative mating, causing its estimates of heritability and of the degree of pleiotropy to be inflated.

#### Historical assortative mating

If, at some point in time, assortative mating for traits ceases and mating becomes random with respect to those traits, the positive trans-LD that was present under assortative mating will immediately disappear, leaving only the positive cis-LD that had built up; this cis-LD will then be gradually eroded by recombination. If equilibrium had been attained under assortative mating, the cis-LD would have grown to match the per-generation trans-LD. Therefore, in the first generation after assortative mating ceases, the upward bias in population GWAS effect size estimates would halve as the trans-LD disappears (Eq. 3); the bias would then shrink gradually to zero as the cis-LD erodes (Fig. 2). A similar pattern will be observed for the heterozygosity-weighted average value of 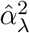 in the population GWAS, which eventually returns to its equilibrium level under random mating (Fig. 3).

In contrast, with the disappearance of the positive trans-LD but the persistence of positive cis-LD, the bias in family-based effect size estimates will suddenly become positive once assortative mating ceases (having temporarily been negative under assortative mating before equilibrium was attained); this bias too will then gradually shrink to zero as recombination erodes the remaining cis-LD (Fig. 2). Concomitantly, the average squared effect size estimate in the family GWAS will suddenly increase when assortative mating ceases, after which it too will gradually return to its equilibrium value under random mating (Fig. 3).

#### Assortative mating between traits with different genetic architectures

An important practical question is how genetic confounding affects the GWAS loci we prioritize for functional follow-up and for use in the construction of polygenic scores. SNPs are usually prioritized on the basis of their GWAS p-value, which is proportional to the estimated variance explained by a SNP, 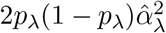 (where *p*_λ_ is the minor allele frequency). The results above assume, in the case of cross-trait assortative mating, that the traits involved have similar genetic architectures (distribution of *p_l_* and *α_l_* at causal loci, and the total number of causal loci). In that case, if there is no pleiotropy between the traits, then while SNPs that tag trait-1 causal loci are predictive of the value of trait 2 owing to LD between trait-1 and trait-2 causal loci, we nonetheless expect the SNPs that tag trait-2 causal loci to be better predictors of trait 2, such that GWAS investigators would primarily pick out SNPs tagging trait-2 causal loci for prioritization and use in polygenic scores.

However, analysis of human GWASs suggests that quantitative traits can have widely different genetic architectures, with, in particular, substantial differences in the effective numbers of causal loci involved and in the distribution of minor allele frequencies (Simons et al. 2022, and references therein). If two traits with distinct genetic bases show cross-trait assortative mating, but trait 1 has a denser genetic architecture (fewer causal loci) than trait 2, then the genetic signal of assortative mating—systematic LD between trait-1 and trait-2 causal loci—will be more heavily loaded per-locus onto trait-1 loci than onto trait-2 loci. In a GWAS on trait 2, this will inflate the magnitude of spurious effect size estimates at SNPs that tag trait-1 loci relative to effect size estimates at SNPs that tag causal trait-2 loci. In Appendix A3.1, we quantify this effect, showing that, in a population GWAS for trait 2, the average magnitude of spurious effect size estimates at trait-1 loci is proportional to 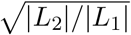, where |*L*_1_| and |*L*_2_| are the numbers of loci underlying variation in traits 1 and 2 respectively. Thus, when trait 1 has a denser genetic architecture than trait 2 (|*L*_2_|/|*L*_1_| is large), the magnitudes of effect size estimates at non-causal trait-1 loci could substantially overlap with those at causal trait-2 loci (as illustrated in Fig. 4), potentially causing part of the apparent, mappable genetic architecture of the trait-2 GWAS to actually tag trait-1 loci.

**Figure 4:**
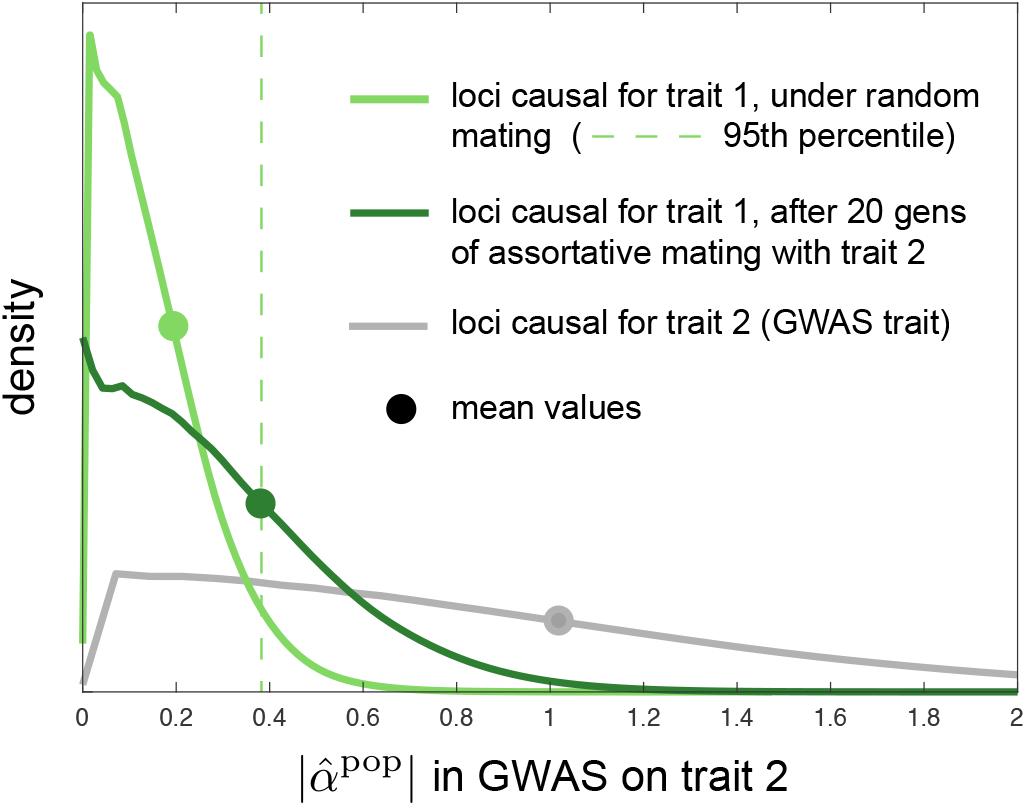
Cross trait assortative mating for traits with different genetic architectures can generate large spurious effect size estimates in population GWASs. Here, the traits have equal heritability, but the number of loci contributing variation to trait 1 is ten-fold smaller than that for trait 2. Shown, for a population GWAS on trait 2, are estimated distributions of the magnitude of effect size estimates at loci causal for trait 2 (grey) and at loci causal for trait 1 (greens), under random mating (light green) and after 20 generations of cross-trait assortative mating (sex-asymmetric, of strength *ρ* = 0.2) for traits 1 and 2 (dark green). Although the true effects of trait-1 loci on trait 2 are zero in these simulations (no pleiotropy), there is sampling noise in effect size estimation at trait-1 loci under random mating (light green line), so that the mean magnitude of effect size estimates is shifted away from zero (light green dot; dashed line displays 95th percentile under random mating). Under assortative mating, the magnitudes of the spurious effect size estimates at trait-1 loci shift significantly rightward (dark green line), coming to overlap substantially with the distribution of effect size estimate magnitudes at causal trait-2 loci (grey line; the distribution for trait-2 loci does not appreciably differ under random and assortative mating). Densities are estimated from pooled effect size estimates from 1,000 replicate simulations. Simulation details in Methods.

### 3.2 Population structure

When a population GWAS draws samples from individuals of dissimilar ancestries, differences in the distribution of causal genotypes, and potentially of environmental exposures, can confound the association study (Lander and Schork 1994; Vilhjálmsson and Nordborg 2013). Correcting for confounds due to population structure has therefore been an important pursuit in the GWAS literature (Spielman et al. 1993; Pritchard et al. 2000; Price et al. 2010).

For concreteness, consider a simple model where two populations diverged recently, with no subsequent gene flow between them. Genetic drift—and possibly selection—in the two populations will have led to allele frequency differences between them at individual loci. If allele frequencies have diverged at both a genotyped study locus and at loci that causally influence the study trait, these frequency differences will manifest as linkage disequilibria between the study locus and the causal loci in a sample taken across both populations, even if the loci are not in LD within either population. Specifically, if the frequencies of the focal allele at a given locus *k* are 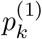 and 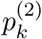 in populations 1 and 2, then the cis-LD between the focal alleles at the association study locus λ and a causal locus *l* is

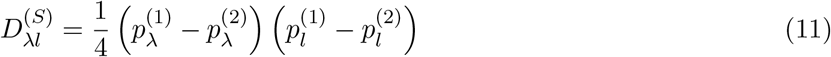

in a sample that weights the two populations equally, with the superscript (*S*) denoting that this LD is due to stratification. The trans-LD takes exactly the same form: 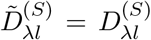. From Eq. (3), locus *l* therefore confounds estimation of the direct effect at λ in a population GWAS, by an amount proportional to

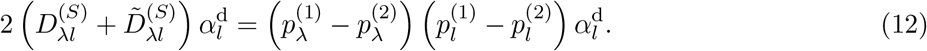

These genetic confounds are in addition to environmental confounding that would arise if the environments of the two populations alter their average trait values by different amounts.

In contrast, estimates of direct effects obtained from within-family association studies are not genetically confounded, because cis- and trans-LD are equal (Eqs. 6 and 7). Another way of seeing this is to consider that, by controlling for family, within-family GWASs control for the population, and in the scenario considered, by construction, there are no within-population LDs to confound effect size estimation.

#### Allele frequency divergence due to drift

How do the confounds introduced by population structure affect the first of our measures of interest, the average deviation of effect size estimates from their true values? The answer depends on the source of allele frequency differences between the two populations. If the differences are due to neutral genetic drift, they will be independent of each other (assuming causal loci are sufficiently widely spaced) and independent of the direction and size of effects at individual loci. Therefore, the LD induced by these allele frequency differences will, on average, not bias effect size estimates in a population GWAS:

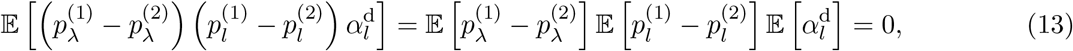

since 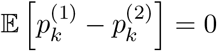 at any locus *k*.

However, the LD induced by population structure will inflate the average squared effect size estimate, and by extension the variance of effect size estimates (Fig. 5). In Appendix A3.2, we quantify this effect for the same simple case of two separate populations. We find that the average squared effect size estimate in a population GWAS is an increasing function of the divergence between the two populations (as measured by *F_ST_*), the number of loci contributing variation to the study trait, and the true average squared effect size per locus (see also Rosenberg and Nordborg 2006; Lee and Lee 2023a).

**Figure 5:**
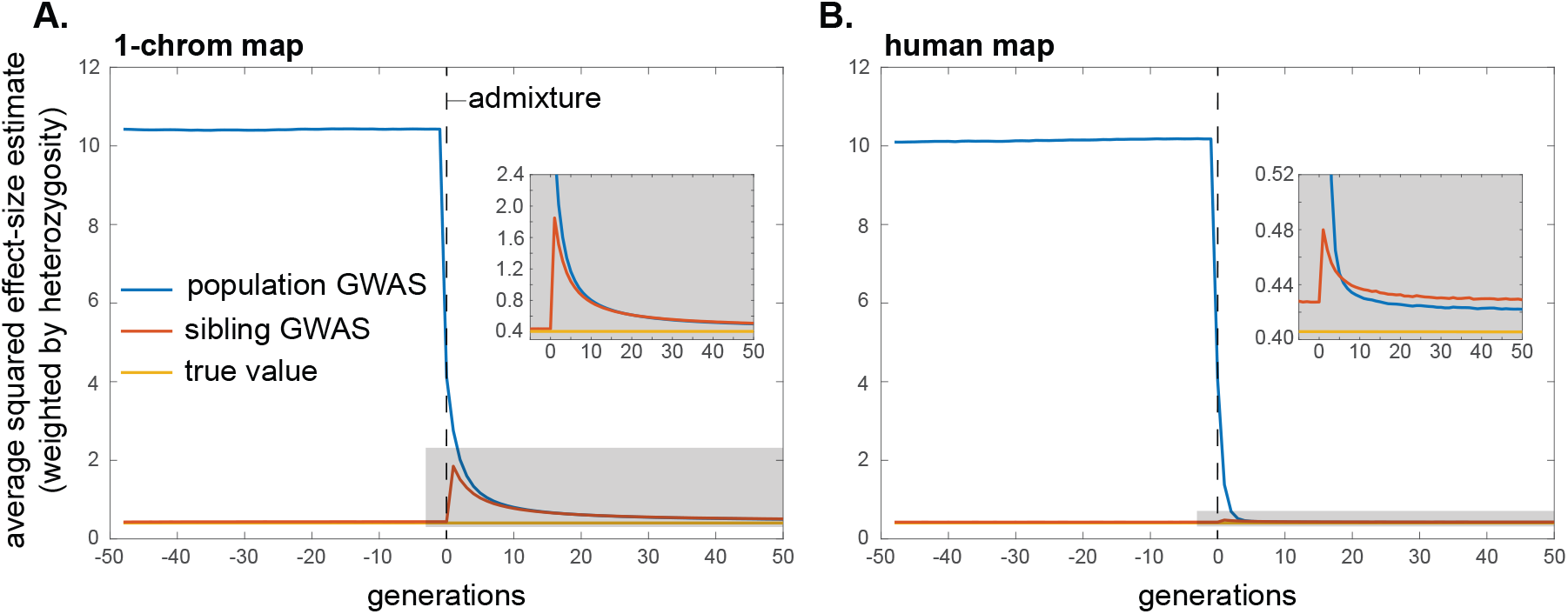
The impact of population structure and admixture on the average squared effect size estimate in population and within-family GWASs. Here, two populations are isolated until generation 0, at which point they mix in equal proportions. Initial allele frequencies are chosen independently for the two populations, such that allele frequency differences between the populations resemble those that would accumulate over time via random drift. As in Fig. 3, the equilibrium value of the mean squared effect size estimate under random mating is greater than the true mean squared effect size, in both the population and within-family GWAS, owing to linkage disequilibria among causal alleles that arise due to drift. This explains why, in the insets, the blue (population) and red (within-family) profiles do not shrink all the way down to the yellow (true) line after admixture, when mating is random. Note too the difference in scale of the y-axes in the insets: the return to equilibrium is much more rapid under the human genetic map than for a hypothetical genome of one chromosome of length 1 Morgan, since, with more recombination, the ancestry-based linkage disequilibria are broken down more rapidly. Profiles are averages across 10,000 replicate simulation trials. Simulation details can be found in the Methods.

In contrast, because effect size estimates from within-family GWASs are not confounded in this model of isolated populations, the average squared effect size estimate will not differ substantially from its expectation in an unstructured population (Fig. 5).

While we have focused on a simple model of two isolated populations, the result that within-family association studies are not confounded holds for other kinds of population structure as well. Specifically, we may be concerned that a population GWAS suffers from genetic confounding along some given axis of population stratification. However, the family-based estimates will be unbiased by confounding along such an axis if the maternal and paternal genotypes at each locus are exchangeable with respect to each other along this axis (Appendix A3.2). This requirement will be met in expectation under many models of local genetic drift in discrete populations or along geographic gradients. However, as we will shortly argue, migration and admixture introduce further complications.

#### Allele frequency divergence due to selection or phenotype-biased migration

Selection and phenotype-biased migration can also generate allele frequency differences among populations (for a review of phenotype-biased migration, see Edelaar and Bolnick 2012). Unlike genetic drift, both of these forces can lead to systematic directional associations between effect sizes and changes in allele frequencies between populations. For example, if selection has favored alleles that increase the trait in population 1 but not in population 2, then

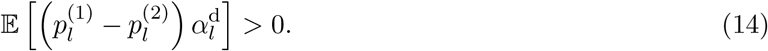

as directional selection causes systematic changes in allele frequencies across the loci *l* underlying variation in the trait under selection (e.g., Hayward and Sella 2022). Importantly, this form of selection can occur even if the mean phenotype of the two populations does not change (Harpak and Przeworski 2021; Yair and Coop 2022). Similarly, phenotype-biased migration, where, say, individuals with a higher value of the phenotype tend to migrate from population 2 to population 1, can also create a positive association between effect sizes and allele frequency differences (Eq. 14).

Unlike the case of neutral genetic drift in the two populations, where the sign of the LD between two alleles is independent of their effect sizes, the effect-size-correlated associations driven by selection or phenotype-biased migration can add up across loci, and thus lead to substantial, systematic biases in estimates of allelic effect sizes. This systematic genetic confounding would also substantially inflate the average squared effect size estimate and thus measures of the genetic variance tagged by SNPs.

In addition, these systematic sources of genetic confounding can generate genetic correlations between traits with no overlap in their sets of casual loci—i.e., with no pleiotropic relationship. This will occur if two traits have both experienced selection or biased migration along the same axis. To take a concrete example, if people tend to migrate to cities in part based on traits 1 and 2, then these traits will become genetically correlated. If this axis is explicitly included as a covariate in the GWAS, then its influence on estimates of heritability and genetic correlations will be removed. However, its influence will not be removed by inclusion of genetic principal components or the relatedness matrix, if this axis (here, city vs. non-city) is not a major determinant of genome-wide relatedness at non-causal loci (Vilhjálmsson and Nordborg 2013). Nor will LD score regression control for this influence, as the selection- or migration-driven differentiation of a variant along the axis will be correlated with the extent to which it tags long-range causal variants involved in either trait. This effect on LD score regression is similar to that discussed above for assortative mating (Border et al. 2022a,b). Thus, like assortative mating, selection and phenotype-biased migration along unaccounted-for axes of population stratification can generate genetic correlations between traits. These selection- and migration-driven correlations should not necessarily be viewed as spurious, since genetic correlations should include those that arise from systematic long-range LD, but they complicate the interpretation of population-level genetic correlations as evidence for pleiotropy.

Again, these issues largely vanish in family-based studies, although phenotype-biased migration can cause transient differences in cis- and trans-LD that lead to biases in family-based estimates of direct effects (Eqs. 6 and 7).

### 3.3 Admixture

When populations that have previously been separated come into contact, alleles from the same ancestral population remain associated with each other in the admixed population until they are dissociated by recombination. If allele frequencies had diverged between the ancestral populations, this ‘ancestry disequilibrium’ can translate to cis-LD between loci affecting a trait (Nei and Li 1973), potentially confounding GWASs performed in the admixed population. More generally, long range LD will be an issue when there is genetic stratification and ongoing migration between somewhat genetically distinct groups.

For concreteness, we again consider a simple model where two populations have been separated for some time, allowing allele frequencies to diverge between them. The populations then come into contact and admix in the proportions *A* and 1 – *A*. We assume that mating is random with respect to ancestry in the admixed population.

Suppose that, just before admixture, the frequencies of the focal allele at a given locus *k* were 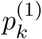 and 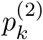 in the two populations. Then the initial degree of cis-LD between loci λ and *l* in the admixed population is given by Eq. (11), weighted by the proportions in which the populations admix:

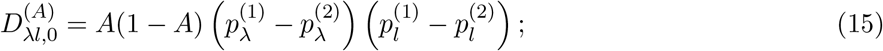

see, e.g., Pfaff et al. (2001). This cis-LD subsequently decays at a rate *c*_λ*l*_ per generation, so that, *t* generations after admixture,

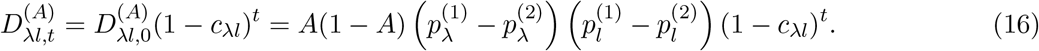

Because we assume that mating is random in the admixed population, the trans-LD is zero in every generation after admixture: 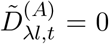. Note that the decay of cis-LD in an admixed population will be slowed if individuals mate assortatively by ancestry, because the trans-LD generated by assortative mating is continually converted by recombination to new cis-LD (as in our assortative mating model above; see Zaitlen et al. (2017) for more discussion of this point in the context of population admixture).

#### Allele frequency divergence due to drift

How do these patterns of LD affect a population GWAS? If allele frequency differences between populations arose from neutral drift, they will be independent of effect sizes at causal loci and across loci, and therefore will not contribute, on average, a systematic directional bias to effect size estimates. However, they will inflate the average squared effect size estimate, by a smaller amount than for a population GWAS performed when the populations were still separated (because of the elimination of trans-LD under random mating in the admixed population). Moreover, this amount will decline in the generations after admixture as the remaining cis-LD is eroded by recombination (Eq. 16; Fig. 5). We quantify these effects in Appendix A3.3 (see also Pfaff et al. 2001; Rosenberg and Nordborg 2006; Zaitlen et al. 2014; Lee and Lee 2023b).

Although within-family GWASs were not genetically confounded when the populations were separate (because cis- and trans-LDs were equal, as discussed above), they become genetically confounded in the admixed population, as all trans-LD is eliminated by random mating in the admixed population, leaving an excess of cis-LD relative to trans-LD that biases effect size estimates (Eqs. 6 and 7). As in the case of the population GWAS, these biases will be zero on average if allele frequency differences between the ancestral populations were due to drift. However, after admixture, they will still inflate the average squared effect size estimate (and thus the variance of effect size estimates), which will thereafter decline in subsequent generations as the cis-LD is gradually broken down by recombination (Eq. 16; Fig. 5).

In comparing the average squared effect size estimate in a population and a family-based GWAS, we observe that the value in the population GWAS rapidly declines to approximately the same level as the value in the within-family GWAS, despite the former having started at a much higher level in the initial admixed population (Fig. 5). The explanation is that LD between unlinked loci confounds effect size estimation in the population GWAS but not the within-family GWAS, such that (i) the average squared effect size estimate from the population GWAS is initially much higher than that from a within-family GWAS, because it is inflated by LD between many more pairs of loci, and (ii) the average squared effect size estimate from the population GWAS declines more rapidly, because LD between unlinked loci is broken down more rapidly than LD between linked loci.

#### Allele frequency divergence due to selection or phenotype-biased migration

In addition to drift, and as discussed above, selection and phenotype-biased migration can generate systematic, signed (effect-size correlated) LD, which would lead to systematic cis-LD in the descendent admixed population. These would lead to larger inflations of genetic variance and genetic correlations than would be expected had allele frequency divergence between the ancestral populations been due to drift alone, and would complicate interpretations of genetic correlations as being due to pleiotropy. Moreoever, if the admixed population is more than a few generations old such that LD between unlinked loci but not linked loci has largely been broken down, then population- and family-based estimates of these quantities might be similar.

#### Spurious genetic correlations due to confounding in population-based PGSs

Factors other than selection and phenotype-biased migration can also generate non-pleiotropic genetic correlation signals in family-based studies of admixed populations. In fact, the use of confounded population GWAS effect sizes can be sufficient. As an example of the confounding of genetic correlations in admixed populations due to a confounded GWAS for one trait, consider the GIANT-GWAS height polygenic score. Owing to confounding within Europe (Berg et al. 2019; Sohail et al. 2019), the height PGS showed large differences between Northern Europeans and sets of individuals sampled in other locations, such as the African 1000 genomes samples (Martin et al. 2017). This confounding generated a spurious, systematic correlation between height effect sizes and allele frequency differences across populations, with height-increasing alleles that are more common among Northern Europeans being assigned larger effects (Berg et al. 2019). As a result, in a PGS constructed from these effect size estimates, larger PGS values are predictive of greater North European ancestry. Now imagine a sibling-based study performed in a sample with recently admixed ‘European’ and ‘non-European’ ancestry—African Americans, for example. An individual with a larger value than their sibling for the GIANT height PGS will, on average, carry more ‘European’ ancestry. In African Americans, there will also be a systematic association of lighter skin pigmentation with recent ‘European’ ancestry, and selection on skin pigmentation will have driven a signed difference in allele frequencies between European and West African ancestors. Putting these observations together, the GIANT height PGS, being predictive of the degree of European ancestry, may well be predictive of skin pigmentation differences between African American sibling pairs (Eq. 10), leading to the naive and incorrect conclusion that height and skin colour are causally linked. In reality, this result would reflect the fact that alleles predicted to increase height and alleles that affect skin color are in systematic effect-signed admixture LD, as in Eq. (15), as a consequence of stratification-biased effect size estimates from the GIANT European GWAS.

### 3.4 Stabilizing selection

Stabilizing selection—selection against deviations from an optimal phenotypic value—is thought to be common (Sella and Barton 2019), and has recently been argued to be consistent with the genetic architectures of many human traits (Simons et al. 2022). By disfavoring individuals with too many or too few trait-increasing alleles, stabilizing selection generates negative cis-LD among alleles with the same directional effect on the trait (Bulmer 1971). Thus, stabilizing selection will attenuate GWAS effect size estimates at genotyped loci that tag these causal loci.

To quantify these biases, we consider the model of Bulmer (1971, 1974), in which a large number of loci contribute to variation in a trait under stabilizing selection, with the population having adapted such that the mean trait value is equal to the optimum. Under this model, stabilizing selection rapidly reduces variance in the trait by generating negative cis-LD among trait-increasing alleles. If we make the simplifying assumption that all loci have equal effect sizes, then the equilibrium reduction in trait variance, – *d*\* (where *d*\* < 0), can be calculated as a function of the genic variance *V_g_*, the environmental noise *V_E_*, the strength of stabilizing selection *V_S_/V_P_* (scaled according to the phenotypic variance *V_P_*), and the harmonic mean recombination rate, 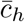, among loci underlying variation in the trait (Bulmer 1974; Appendix A3.4).

Under these same assumptions, we calculate in Appendix A3.4 the average per-locus attenuation bias in effect size estimates induced by stabilizing selection, 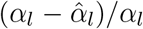. In a population GWAS, this attenuation bias is approximately

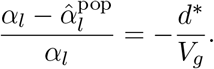

In a within-family GWAS, the average proportionate bias is approximately

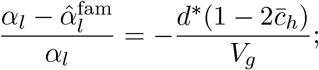

i.e., smaller than in a population GWAS by a factor of 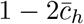.

Thus, the bias in effect size estimation can be calculated given estimates of the phenotypic variance and heritability of the trait, the harmonic mean recombination rate, and the strength of stabilizing selection (Appendix A3.4). In the Methods, making some simplifying assumptions about the genetic architecture of the trait in question, we calculate an approximate value 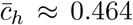 for humans. Using this value, Fig. 6 shows the average proportionate reduction in GWAS effect size estimates for various strengths of stabilizing selection and heritabilities of the trait. The range of selection strengths was chosen to match that inferred for human traits by Sanjak et al. (2018).

**Figure 6:**
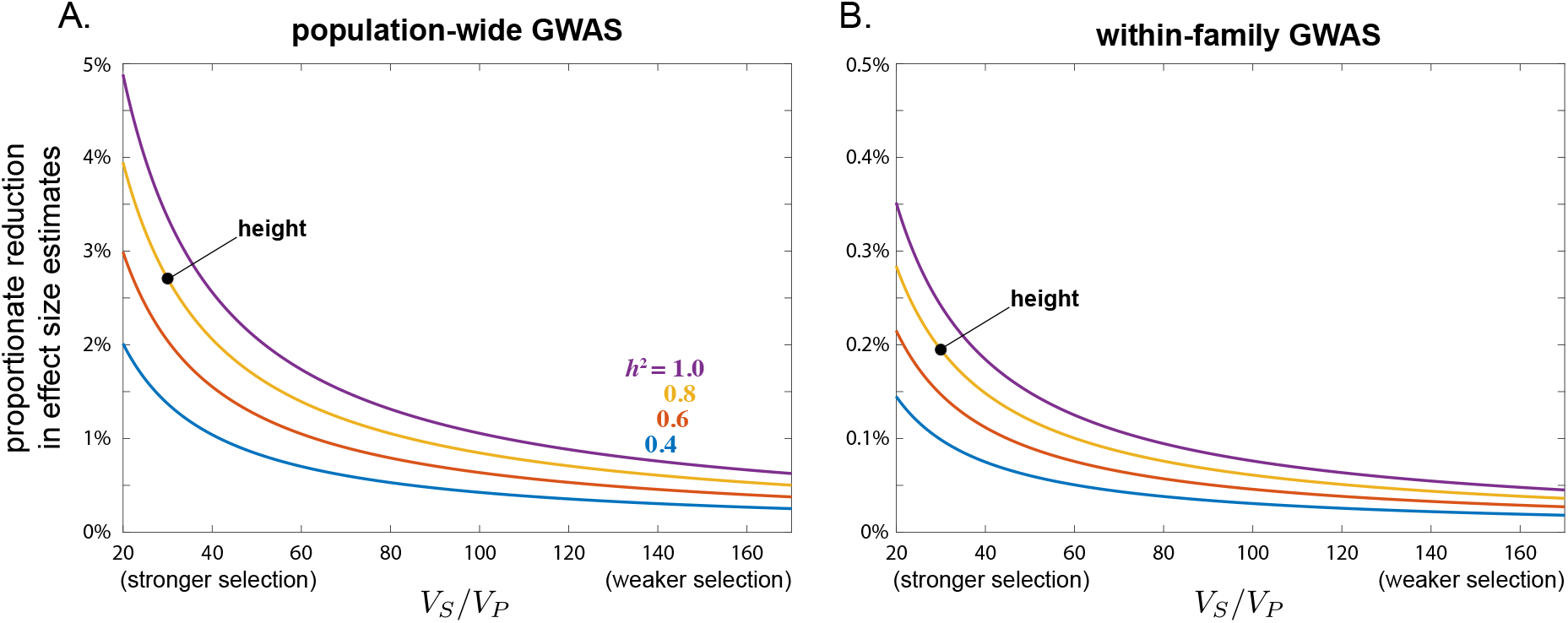
Stabilizing selection attenutates GWAS effect size estimates. The calculations displayed here assume that genetic variation in the trait is contributed by 1,000 loci of equal effect spaced evenly along the human genome. Stabilizing selection is stronger if the width of the selection function scaled by the phenotypic variance, *V_S_/V_P_*, is smaller. The placement of the point for human height assumes a heritability of 0.8 and a strength of stabilizing selection of *V_S_/V_P_* = 30, as estimated by Sanjak et al. (2018). Details of these calculations can be found in Appendix A3.4. Note the different scales of the y-axes in A and B.

Attenuation of effect size estimates is larger if stabilizing selection is stronger or if the trait is more heritable. Taking height as an example, heritability is ~0.8, *V_P_* ≈ 7cm^2^, and Sanjak et al. (2018) estimate a sex-averaged strength of stabilizing selection of *V_S_*/*V_P_* ≈ 30. From these values, we calculate that a population GWAS would systematically underestimate effect sizes at loci that causally influence height by about 3% on average, in the absence of other sources of LD (Fig. 6A). More generally, within the range of reasonable strengths of stabilizing selection inferred by Sanjak et al. (2018), we calculate average attenutations of population-based effect size estimates of up to 5% for highly heritable traits (*h*^2^ ≈ 1) under strong stabilizing selection (*V_S_/V_P_* ≈ 20), down to 0.25% for less heritable traits (*h*^2^ ≈ 0.4) under weak stabilizing selection (*V_S_*/*V_P_* ≈ 170) (Fig. 6A).

Given the estimate 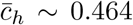, the proportionate bias that stabilizing selection induces in within-family GWASs is expected to be a fraction 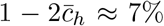 that in population-based GWASs. Thus, for height, a within-family GWAS would underestimate effect sizes by only about 0.2% on average (Fig. 6B).

The quantitative importance of these biases will vary by application. In situations where the goal is gene discovery, for example, 5% reductions in effect size estimates are unlikely to flip the statistical significance of variants with large effects on a trait. However, the attenuations in effect size estimates caused by stabilizing selection are systematic across loci, and therefore could substantially affect aggregate quantities based on these estimates. For example, the range of average reductions in population effect size estimates calculated above for human traits would translate to reductions in naive estimates of SNP-based heritabilities of between 0.5% and 10% (~6% in the case of height). If effect sizes are estimated by within-family GWAS, on the other hand, the reductions in these SNP-based heritability estimates would be much smaller.

As a further example, by generating negative LD between alleles with the same directional effect on the trait, the impact of stabilizing selection opposes, and therefore masks, the genetic impact of assortative mating (Brown et al. 2016). A practical consequence is that stabilizing selection will tend to attenuate estimates of the strength of assortative mating based on GWAS effect sizes, which often use cross-chromosome correlations of polygenic scores (e.g., Yengo et al. 2018; Yamamoto et al. 2023). In humans, the phenotypic correlation among mates for height has been measured at about ~0.25 (Stulp et al. 2017). In Appendix A3.4, we calculate that estimates of this correlation based on cross-chromosome correlations in PGSs will be biased downwards by about 20%, to ~0.20, because of stabilizing selection on height. Were assortative mating weaker, or stabilizing selection stronger, the genetic impact of assortative mating would be masked to an even greater extent (Appendix A3.4).

As in our analysis of assortative mating above, if stabilizing selection ceases in some generation, the negative LD that built up during the period of stabilizing selection will decay over subsequent generations, rapidly for pairs of loci on different chromosomes and more slowly for linked pairs of loci. Patterns of selection on human traits have changed over time—for example, the strength of stabilizing selection on birth weight has relaxed (Ulizzi and Terrenato 1987). In general, therefore, patterns of confounding reflect a composite of contemporary and historic processes.

### 3.5 Sibling indirect effects

Indirect effects of siblings’ genotypes on each other’s phenotypes are known to be a potential source of bias in sibling-based GWASs (Fletcher et al. 2021; Young et al. 2022), and can be measured and corrected only if, in addition to sibling genotypes, parental genotypes are also available (Kong et al. 2018; Young et al. 2022). To generate intuition for their impact on GWASs, we consider a simple model of indirect sibling effects in the absence of G×E interactions and other confounding effects, focusing on a single-locus model for simplicity. We suppose that the indirect effect of an individual’s phenotype on their sibling’s phenotype is β, so that the phenotypes of two siblings *i* and *j* can be written

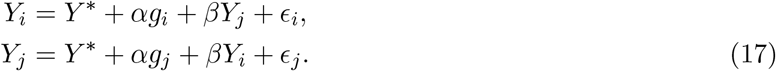

Taking their difference and rearranging, we find that

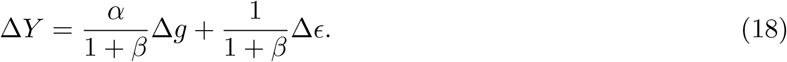

Therefore, in the absence of genetic confounding and G×E interactions, a sibling-based association study would return an effect size estimate of

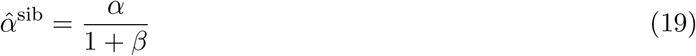

on average. Thus, if sibling indirect effects are synergistic (*β* > 0), they lead to underestimation of the direct genetic effect at the locus. In contrast, if sibling indirect effects are antagonistic (*β* < 0), they lead to overestimation of the direct genetic effect.

How would a population GWAS be affected by the same sibling indirect effects? Sibling *i*’s phenotype can be written

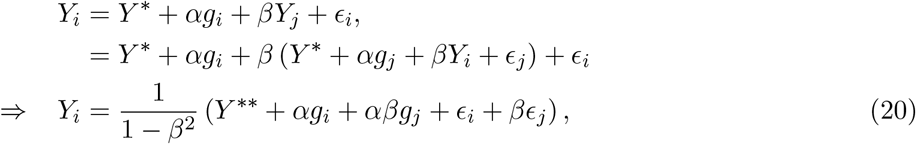

where *Y*\*\* = (1 + *β*)*Y*\*. Therefore, if we were to randomly choose one sibling from each sibship and estimate the effect size at the locus using a population association study across families, we would obtain

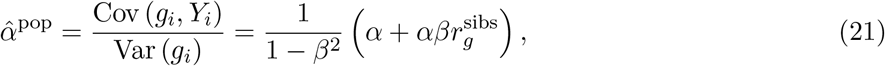

where 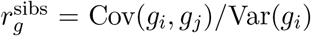 is the genotypic correlation between sibs at the locus. Sibling indirect effects alter the effect size estimate in a population GWAS via two channels. The first is through the factor 1/(1 – *β*^2^) in Eq. (21), which reflects second-order feedbacks of an individual’s phenotype on itself, via the sibling. Since 1/(1 – *β*^2^) > 1, these feedbacks act to exacerbate the effects of causal alleles. For example, if sibling indirect effects are antagonistic (*β* < 0), then a sibling with a large trait value will tend to indirectly reduce the trait value of their sibling, which in turn will indirectly further increase the trait value of the focal individual. This channel therefore pushes population GWASs towards overestimating the magnitude of direct genetic effects.

The other channel by which sibling indirect effects can influence a population GWAS is driven by the genotypic correlation among siblings, and is easiest to understand if we assume that sibling indirect effects are weak (*β*^2^ ≪ 1). In this case, 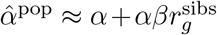. Since the genotypic correlation 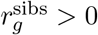, this channel of sibling indirect effects has the opposite effect to the one it has on a sibling GWAS: if sibling indirect effects are synergistic (*β* > 0), the population GWAS overestimates the direct genetic effect at the locus, while if sibling indirect effects are antagonistic (*β* < 0), the population GWAS underestimates the direct genetic effect. The reason for this difference is that a sibling GWAS is based on siblings whose genotypes differ at the focal locus, and whose genotypic values are therefore anticorrelated. If sibling indirect effects are synergistic (*β* > 0), they will tend to attenuate the phenotypic differences between such siblings, and therefore attenuate effect size estimates. In contrast, because siblings’ genotypes are positively correlated across the entire population, synergistic sibling indirect effects (*β* > 0) will tend to exacerbate phenotypic differences across families, leading a population GWAS to overestimate effect sizes.

### 3.6 Gene-environment (GxE) and gene-gene (GxG) interactions

Up to this point, we have assumed that alleles’ direct effects do not vary across environments or genetic backgrounds. To generate intuition for the influence of G×E (and G×G) interactions on population and family-based GWAS designs, we restrict our focus to a single causal locus, assuming no genetic confounding and no indirect effects of siblings. To incorporate G×E interactions, we allow the effect size of the alleles at the locus to depend on the family environment. The phenotype of individual *i* in family *f* is

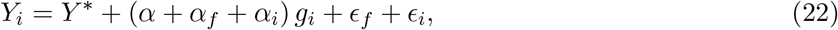

where we arbitrarily define *α_f_* and *α_i_* such that their population means are zero: 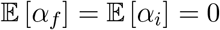. *α* is then the average causal effect of the allele were it randomized across individuals from different families in our sample. *α_f_* is the deviation of this effect in family *f* due to their environment, and *α_i_* is an individual deviation which we assume to be independent of *i*’s genotype; both *α_f_* and *α_i_* can be thought of as random slopes in a mixed model. Note that *α_f_* can reflect the interaction of alleles with the family’s external environment (as we have framed it here) or with the family’s genetic background (a G×G interaction).

If we perform a sibling GWAS by taking pairs of full siblings *i* and *j* in family *f* and regressing the difference in their phenotypes Δ*Y_f_* = *Y_i_* – *Y_j_* on the difference in their genotypes at the focal locus Δ*g_f_* = *g_i_* – *g_j_*, we obtain an effect size estimate

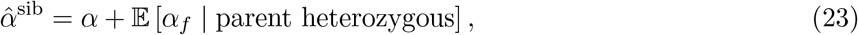

where the second term—the deviation of the family-based estimate 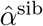 from *α*—is the average family deviation conditional on a parent being heterozygous at the focal locus (Appendix A4). The intuition is that, because only heterozygous parents contribute the genetic variation among siblings on which our effect size estimate is based, if these heterozygous parents are non-randomly distributed across environments, then the family-based GWAS samples values from a distribution of family effects *α_f_* that is different to the overall population distribution.

We can compare this estimate from a sibling GWAS to one from a population GWAS, again under the assumption of no genetic confounding or indirect effects from siblings:

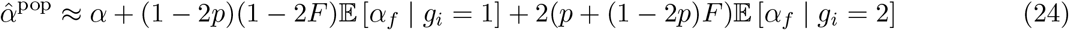

where *p* is the frequency of the focal variant and *F* is the inbreeding coefficient at the locus (Appendix A4). The approximation holds if *F* is small. Note that Eq. (24) conditions on the number of focal alleles carried by the sampled individual, whereas Eq. (23) conditions instead on the parental genotype.

Like the family-based estimate, the population-based effect size estimate is distorted when heterozygotes are not randomly distributed over family backgrounds (*E* [*α_f_* | *g_i_* = 1] ≠ 0) as well as when homozygotes are not randomly distributed across family backgrounds (when *E* [*α_f_* | *g_i_* = 2] ≠0). Thus, effect size estimates from both family- and population-GWAS can differ from the genetic effects that would be estimated if genotypes were randomly distributed across interacting family backgrounds, and these distortions will in general not be the same aross population and family-based study designs.

As noted above, because of current sample size constraints in family-based studies, a common strategy is to calculate the association of population-based PGSs and phenotypic differences among family members. In the absence of confounding, it is clear from Eq. (10) that the influence of G×E interactions on the covariance of sibling differences in PGSs and trait values would depend on the average value of 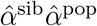 across loci. Thus the slope of the PGS in this regression could be affected if, on average, the alleles at casual loci tagged by genotyped variants in the PGS are more often found in environments that suppress (or enhance) their effects. G×E interactions across many loci have been suggested by some recent studies (Mostafavi et al. 2020; Zhu et al. 2022), but their quantitative impact on differences between population and family-based GWASs remains unknown.

An allele’s effect could also systematically differ across families (*α_f_*) if it is involved in epistatic interactions with alleles at other loci in the genome (G×G). By analogy to our G×E model above, epistatic interactions would lead to biases in family-based GWASs if parents who are heterozygous at the focal study locus tend to have systematically different genotypes at loci that interact epistatically with the focal locus, relative to the population distribution of such genetic backgrounds.

Up to this point, we have also ignored parent-offspring interactions as a possible source of bias in family-based studies. Following the the same logic above, interactions between parents’ and offsprings’ alleles will result in family GWAS estimates that are the average effect of the focal allele in an offspring conditional on the genetic background of a heterozygous parent. Thus, again, a non-random distribution of genetic backgrounds in heterozygous parents is a potential source of bias.

One way that heterozygous parents might exhibit a non-random distribution of genetic backgrounds is via trait-based assortative mating, which could therefore modify the way that epistasis and parent-offspring interactions influence effect size estimation in a family-based GWAS relative to a population GWAS and relative to the true average population effect.

A final, overarching complication is that the individuals participating in a population GWAS are not a random subset of the population(s) from which they are drawn (Fry et al. 2017; Pirastu et al. 2021; Tyrrell et al. 2021), and families enrolled in GWASs can be even less representative of the population as a whole (Mostafavi et al. 2020; Benonisdottir and Kong 2022). These participation biases can potentially lead to systematic differences between the distributions of genotypes and interacting environments experienced by the population, the GWAS sample, and participants in a family-based study.

## 4 Discussion

It has long been recognized that population GWASs in humans can be biased by environmental and genetic confounding (Lander and Schork 1994; Vilhjálmsson and Nordborg 2013). Currently, population GWASs attempt to control for these confounds by focusing on sets of individuals that are genetically more similar and by controlling for population stratification. However, these controls are imperfect and are not always well defined. For example, controlling for genome-wide patterns of population stratification based on common alleles does not control for the genetic and environmental confounding of rare variants (Zaidi and Mathieson 2020). Work on genetic confounding has uncovered increasing evidence that assortative mating may be leading to large biases in estimates of direct genetic effects and to large genetic correlations for a number of traits (Yengo et al. 2018; Border et al. 2022b,a); moreover, it can often be unclear whether genetic signals of assortative mating are due to trait-based mate choice or some other more general form of genetic confounding (e.g., Haworth et al. 2019). Additionally, while we have focused primarily on genetic confounding, for a number of traits there are also signals of residual environmental confounding in GWAS signals (Selzam et al. 2019; Mostafavi et al. 2020; Okbay et al. 2022; Abdellaoui et al. 2022). Thus, subtle and often interwoven forms of genetic and environmental confounding remain a major issue in many GWASs (Young et al. 2022), compromising the interpretation of GWAS effect size estimates and downstream quantities such as SNP heritabilities and genetic correlations.

Effect size estimates from within-family GWASs are less affected by these various confounds. In the absence of G×E interactions, they are not subject to environmental confounding across families, because the environments of family members are effectively randomized with respect to within-family genetic transmission. As we have shown, family-based estimates should also suffer substantially less from genetic confounding, because genetic transmission at unlinked loci (but not linked loci) is randomized by independent assortment of chromosomes in meiosis. Nonetheless, family-based GWASs can suffer from residual genetic confounding as well as sibling indirect effects and G×E/G×G interactions; they also raise a number of conceptual problems that we discuss below.

### Sources of genetic confounding

Genetic confounding is caused by long-range LD between loci that affect the trait or traits under study. To illustrate the potential for genetic confounds to bias GWAS effect size estimates, we have considered several sources of long-range LD. Some of these—assortative mating, selection on GWAS traits, and phenotype-biased migration—can cause systematic directional biases in GWAS effect size estimates. Others, such as neutral population structure, cause random biases that influence the variance of effect size estimates and related quantities. Assortative mating and neutral population structure have received considerable theoretical attention in the GWAS literature (e.g., Rosenberg and Nordborg 2006; Yengo et al. 2018; Border et al. 2022a,b). Here, we have further outlined how both selection and phenotyped-biased migration can drive systematic genetic confounding that may not be well accounted for by current methods of controlling for stratification.

We wish to emphasize stabilizing selection in particular as a potential source of systematic confounding in GWASs. Stabilizing selection has been well studied in the quantitative genetics literature but less so in the context of GWASs, despite its expected ubiquity. By selecting for compensating combinations of trait-increasing and trait-decreasing alleles, stabilizing selection generates negative LD between alleles with the same directional effect on the trait (Bulmer 1971, 1974), and can therefore bias GWAS effect size estimates downwards. While the potential for stabilizing selection to confound effect size estimation has been noted (e.g., Brown et al. 2016; Yair and Coop 2022; Li et al. 2023), the resulting biases have not, to our knowledge, been quantified. Our calculations suggest that these downward biases could, for some human traits, be as large as 5% systematically across all causal loci in population GWASs. While biases of this magnitude are unlikely to compromise some goals of GWASs, such as gene discovery, they could be quantitatively problematic for other GWAS aims, such as estimation of SNP heritabilities and the strength of assortative mating. Moreover, while our results pertain to (a particular model of) stabilizing selection, many kinds of selection generate LD between genetically distant loci—in fact, only multiplicative selection among loci does not (Bürger 2000, pgs. 50 and 177). Therefore, the general result that selection can generate genetic confounding will hold more broadly.

For a given genotyped locus in a GWAS, there is no bright line between local ‘tagged’ LD and long-range confounding LD, and one reasonable objection to the approach taken here is that that we have used an arbitrary definition of the causal loci that are locally tagged by a genotyped locus (*L*_local_ in Eq. 2). All of the sources of genetic confounding that we have considered generate LD among causal loci both within and across chromosomes. Under these models, the within-chromosome LD that is generated is, in a sense, a continuation of the LD generated across chromsomes (moving from a recombination rate = 0.5 to ≤ 0.5). Thus, while investigators may prefer some looser definition of ‘local’ when thinking about genotyped GWAS loci as tag SNPs, to extend that definition to include all loci on the same chromosome as the SNP would, by reasonable interpretation, be to include confounding into the desired estimator.

The extent to which the absorption of genetic confounding in estimated effect sizes is a problem depends on the application. In the case of polygenic prediction, absorbing environmental effects, indirect effects, the effects of untyped loci throughout the genome can help to improve prediction accuracy, although this does come at a cost to interpretability. For GWAS applications focused on understanding genetic causes and mechanisms, the biases in effect size estimates and spurious signals of pleiotropy among traits generated by genetic confounding will be more problematic.

### Indirect genetic effects

Family GWASs are often interpreted as providing the opportunity to ask to what extent parental genotypes (or other family genotypes) causally affect a child’s phenotype (‘genetic nurture’; Kong et al. 2018). Viewed in this way, the association between untransmitted parental alleles and the child’s phenotype would seem, at first, a natural estimate of indirect genetic effects.

In practice, however, if the population GWAS suffers from genetic and environmental confounds, then the estimated effects of untransmitted alleles will absorb that confounding in much the same way that estimates of direct genetic effects from a population GWAS do (Eq. 8; Shen and Feldman 2020). For example, in the case of assortative mating, a given untransmitted allele is correlated with alleles that were transmitted both by this parent and by their mate, and these transmitted alleles can directly affect the offspring’s phenotype. Thus, while family-based estimates of direct genetic effects benefit from the randomization of meiosis and from controlling for the environment, family-based estimates of indirect genetic effects lack both of these features and should be interpreted with caution. Indeed, recent work using parental siblings to control for grandparental genotypes has shown that little of the estimated ‘indirect genetic effect’ may be causally situated in parents (Nivard et al. 2022). With empirical estimates of indirect genetic effects potentially absorbing a broad set of confounds (Demange et al. 2022; Young et al. 2022), and few current studies of indirect effects having designs that allow such confounding to be disentangled, it is premature—and potentially invalid—to interpret associations of untransmitted alleles causally in terms of indirect genetic effects (Wolf et al. 1998). Rather, they should be treated agnostically in terms of ‘non-direct’ effects.

### Direct genetic effects

Mendelian segregation provides a natural randomization experiment within families (Fisher 1952), and so crosses in experimental organisms and family designs have long been an indispensable tool to geneticists in exploring genetic effects and causation. Growing concerns about GWAS confounding and the increasing availability of genotyped family members have led to a return of family-based studies to the association study toolkit (Young et al. 2019). Family-based estimates of direct genetic effects are often interpreted as being unbiased and discussed in terms of the counterfactual effect of experimentally substituting one allele for another (Morris et al. 2020; Brumpton et al. 2020; Young et al. 2022).

As we have shown, family-based GWASs are indeed less subject to confounding than population-based GWASs: in the presence of genetic and environmental confounding, the family-based estimate of the effect size at a given locus provides a much closer approximation to the true effects of tightly linked causal loci than a population-based estimate does. The family-based estimate is not biased by environmental variation across families and avoids the correlated effects of the many causal loci that lie on other chromosomes. Still, the family-based estimate does absorb the effects of non-local causal loci on the same chromosome, and so cannot truly be said to be free of genetic confounding. Rather than considering a single allele being substituted between individuals, a better experimental analogy for the effect size estimate would be to say that we are measuring the mean effect of transmission of a large chunk of chromosome surrounding the focal locus, potentially carrying many causal loci.

In addition, while within-family GWASs offer these advantages, in other ways, they move us further away from the questions about the sources and causes of variation among unrelated individuals that motivate population GWASs in the first place. Indeed, the presence of confounding introduces a number of conceptual issues in moving from within-family GWAS to the interpretation of differences among individuals from different families (Coop and Przeworski 2022a,b). For example, in the presence of genetic confounding, the effect of a causal allele of interest will depend on a set of weights: its LD to many other causal alleles. In estimating the direct effect of the allele, family-based approaches weight these LD terms differently to population-based approaches, which, we argue, can complicate the interpretation of these estimates. For example, when previously isolated populations admix, same-ancestry alleles will be held together in long genomic blocks until these are broken up by recombination, which will happen very quickly for alleles on different chromosomes but more slowly for alleles on the same chromosome. A few generations after admixture, therefore, cross-chromosome ancestry LD will largely have dissipated, but contiguous ancestry tracts will still span substantial portions of chromosome lengths. Since both population and within-family GWASs are similarly confounded by the same-chromosome LD, their mean squared effect sizes will be similar in this case (Fig. 5). Bearing in mind that the LD resulting from admixture is not present in the source populations, it becomes unclear which weighting of ancestry LD is appropriate if we want to interpret the resulting effect size estimates as direct effects. As this example illustrates, while family-based GWASs are a useful device for dealing with confounding, it is not always obvious how to interpret the quantities that they measure.

A number of additional complications arise when, to compensate for the small effect sizes of individual loci, researchers combine many SNPs into a polygenic score (PGS) and study the effects of PGSs within families (or use them as instruments in Mendelian randomization analyses). For one, SNPs are usually chosen for inclusion in the PGS on the basis of their statistical significance in a population GWAS. This approach prioritizes SNPs whose effect size estimates are amplified (or even wholly generated) by confounding (for an example of how this leads to residual environmental confounding in applications of sibling-based effect size estimates, see Zaidi and Mathieson 2020). Second, the weights given to SNPs that are included in the PGS absorb the effects of confounding, and this confounding is heterogeneous across SNPs. Thus, when we study the correlates of trait-A PGS differences between siblings in the presence of GWAS confounding, we are not observing the average phenotypic outcomes of varying the genetic component of trait A between siblings. Rather, we are varying a potentially strangely-weighted set of genetic correlates of trait A.

An observation that a population GWAS PGS is predictive of phenotypic differences among siblings demonstrates that the PGS SNPs tag nearby causal loci, but beyond that, interpretation is difficult. Notably, if there is cross-trait assortative mating for traits A and B, but no pleiotropic link between the traits, then some of the SNPs identified as significant in a GWAS on trait A may be tightly linked to loci that causally affect trait B but not trait A. If these loci are included in the trait-A PGS, then when we study the effect of variation in the trait-A PGS on sibling differences, we are accidentally absorbing some components of the variation in trait B across siblings. Thus, we might observe a correlation between the trait-A PGS and differences in trait B between siblings, and this correlation may be lower than is observed at the population level, without there existing any pleiotropic (or causal) link between A and B. These effects can be exacerbated if the two traits have different genetic architectures (Figure 4). Instead of using a set of SNPs and weights from a population GWAS, genetic correlations between traits due to pleiotropy could be estimated from the correlation of effect sizes estimated within families (Howe et al. 2022). Given current sample size constraints in family-based studies, the confidence intervals on these estimates are large. Moreover, significant family-based correlations need not reflect pure pleiotropy, since, as we have shown, they are not completely free of genetic confounding due to intra-chromosomal LD.

Also complicating the interpretation of family-based effect size estimates are various types of interactions. Indirect effects between siblings can bias family estimates of direct genetic effects (Eq. 19; Young et al. 2019; Fletcher et al. 2021; Young et al. 2022) in ways that are conceptually different from the biases they introduce to population-based estimates (Eq. 21). These sibling effects can potentially be addressed with fuller family information (e.g., parental genotypes in addition to sibling genotypes; Kong et al. 2018; Young et al. 2022).

As we have further shown here, G×E (and G×G) interactions can also complicate the interpretation of family-based effect size estimates. The reason is that, even if we were to know the causal alleles for a trait of interest, what we estimate by measuring their associations with phenotypic differences within families is not analogous to the counterfactual effects of experimentally substituting alleles in random individuals. Instead, we are necessarily restricting our focus to the effect of their transmission from heterozygous parents. If heterozygous parents tend to experience different environments or carry different genetic backgrounds than homozygotes do, within-family designs will tell us about direct effects in these particular environments or genetic backgrounds, rather than in the population as a whole. Thus, although the ongoing shift towards family-based studies is motivated by concerns about confounding, with different alleles experiencing different environmental and genetic backgrounds, family-based studies can be influenced by conceptually similar issues of confounding in the presence of G× E and G×G interactions. Such interactions are difficult to reliably identify and measure, but there are a growing number of potential examples from GWASs (Tropf et al. 2017; Barcellos et al. 2018; Young et al. 2018b; Mostafavi et al. 2020; Patel et al. 2022). The interaction issues raised here echo a set of conceptually distinct concerns about the interpretation of average treatment effects in other contexts (Słoczyński 2022), reinforcing the need for care in interpreting such estimates as informative about causes across heterogeneous groups.

In summary, family-based studies are a clear step forward towards quantifying genetic effects, with large-scale family studies carrying the potential to resolve long-standing issues in human genetics. However, these designs come with their own sets of caveats, which will be important to understand and acknowledge as family-based genetic studies become a key tool in the exploration of causal effects across disparate fields of study.

## Acknowledgements

We thank Jeremy Berg, Doc Edge, Arbel Harpak, Hanbin Lee, Molly Przeworski, and members of the Coop lab for helpful discussions and feedback on earlier drafts. Funding was provided by the National Institutes of Health (NIH R35 GM136290 awarded to GC) and a Branco Weiss fellowship to CV.

## Methods

All simulations were carried out in SLiM 4.0 (Haller and Messer 2019). Code is available at github.com/cveller/confoundedGWAS.

For the purpose of carrying out sibling association studies in our simulations, we assumed a simple, monogamous mating structure: each generation, each female and each male is involved in a single mating pair, and each mating pair produces exactly two offspring (who are therefore full siblings). To maintain the precisely even sex ratio required by this scheme, we assumed that a quarter of mating pairs produce two daughters, a quarter produce two sons, and half produce a son and a daughter. Population sizes were chosen to ensure that these numbers of mating pairs were whole numbers, and mating pairs were permuted randomly each generation before assigning brood sex ratios (to ensure that no artifact was introduced by SLiM’s indexing of individuals).

Each generation, per-locus effect size estimates were calculated for both population-wide and sibling GWASs. The former were calculated as the regression of trait values on per-locus genotypes, while the latter were calculated as the regression of sibling differences in trait values on sibling differences in per-locus genotypes.

In all simulations, the total population size was *N* = 40,000.

### Assortative mating

For our general cross-trait assortative mating setup, traits 1 and 2 are influenced by variation at sets of bi-allelic loci *L*_1_ and *L*_2_ respectively. The effect sizes of the reference allele at locus *l* on traits 1 and 2 are *α*_l_ and *β*_l_ respectively. An individual’s polygenic score (PGS) is then 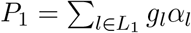 for trait 1 and 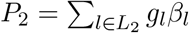 for trait 2. In all the scenarios we simulated, traits had heritability 1, so that individuals’ trait values are the same as their PGSs.

Our aim is to simulate a scenario where assortative mating is based on females’ values for trait 1 and males’ values for trait 2, such that, across mating pairs, the correlation of the mother’s PGS for trait 1, 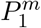, and the father’s PGS for trait 2, 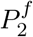, is a constant value *ρ* (in all of our simulations, *ρ* = 0.2). To achieve this, we use an algorithm suggested by Zaitlen et al. (2017): At the outset, we choose an accuracy tolerance *ε* such that, if by some assignment of mates the correlation of their PGSs falls within *ε* of the target value *ρ*, we accept that assigment. Each generation in which assortative mating occurs, we rank females in order of their PGSs for trait 1, and males in order of their PGSs for trait 2. We then calculate the PGS correlation across mating pairs, *ρ*_0_, if females and males were matched according to this ranking. If this (maximal) correlation is smaller than the upper bound of our target window (*ρ*_0_ < *ρ* + *ε*, which very seldom occurred in our simulations), then females and males mate precisely according to their PGS rankings and we move on to the next generation. If, instead, *ρ*_0_ exceeds *ρ* + *ε*, then we follow the following iterative procedure until we have found a mating structure under which the correlation of PGSs falls within *ε* of the target value *ρ*.

First, we choose initial ‘perturbation sizes’ *ξ*_0_ and *ξ*_1_ = 2*ξ*_0_. Suppose that, in iteration *k* of the procedure, the perturbation size is *ξ_k_* and the chosen mating structure leads to a correlation among mates of *ρ_k_*. If |*ρ_k_* – *ρ*| < *ε*, we accept the mating structure and move on to the next generation. Otherwise, we choose a new perturbation size *ξ*_*k*+1_: (i) if *ρ*_*k*–1_, *ρ_k_* > *ρ*, then *ξ*_*k*+1_ = 2*ξ_k_*; (ii) if *ρ*_*k*–1_ > *ρ* > *ρ_k_* or *ρ*_*k*–1_ < *ρ* < *ρ_k_*, then *ξ*_*k*+1_ = (*ξ*_*k*–1_ + *ξ_k_*)/2; (iii) if *ρ*_*k*–1_, *ρ_k_* < *ρ*, then *ξ*_*k*+1_ = *ξ_k_*/2. Once we have chosen *ξ*_*k*+1_, for each individual we perturb their PGS (trait 1 for females; trait 2 for males) by a value chosen from a normal distribution with mean 0 and standard deviation *ξ*_*k*+1_, independently across individuals. We then rank females and males according to their perturbed PGSs, and calculate the correlation *ρ*_*k*+1_ of their true PGSs if they mate according to this ranking. (Since, in our experience, there can be substantial variance in the *ρ*_*k*+1_ values that result from this procedure, we repeat it 5 times and choose the mating structure that produces the value of *ρ*_*k*+1_ closest to the target value *ρ*.) We then decide if another iteration—i.e., another perturbation size *ξ*_*k*+2_—is required.

### Fig. 2. Cross-trait assortative mating for traits with the same genetic architecture

In the simulations displayed in Fig. 2, *ρ* = 0.2, and traits 1 and 2 have identical but non-overlapping genetic architectures: *L*_1_ and *L*_2_ are distinct sets of 500 loci each, with *α_l_* = 1 and *β_l_* = 0 for *l* ∈ *L*_1_, and *α_l_* = 0 and *β_l_* = 1 for *l* ∈ *L*_2_. Loci in *L*_1_ and *L*_2_ alternate in an even spacing along the physical (bp) genome. Fig. 2A shows results for the ‘single chromosome’ case where the recombination fraction between adjacent loci is *c* = 1/999 in both sexes (such that the single-chromosome genome receives, on average, one crossover per transmission). Fig. 2B shows results for the case where recombination fractions between loci are calculated from the human female and male linkage maps generated by Kong et al. (2010). In both cases, we assumed no crossover interference.

At each locus, the initial frequency of the reference allele was 1/2, with reference alleles assigned randomly across diploid individuals and independently across loci such that, in expectation, Hardy-Weinberg and linkage equilibrium initially prevail. The assortative mating algorithm above was run for 19 generations, with a target correlation *ρ* = 0.2, a tolerance parameter *ε* = *ρ*/100, and an initial perturbation size 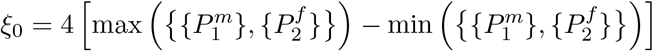. Thereafter, assortative mating was switched off, with mating pairs (still monogamous) being chosen randomly.

### Fig. 3. Same-trait assortative mating

The algorithm we followed to ensure assortative mating of a given strength was the same as that for Fig. 2 above, but here traits 1 and 2 are identical. 1,000 loci underlie variation in the trait, and are evenly spread along the physical genome. The effect size of the reference allele at each locus was drawn from a normal distribution with mean 0 and standard deviation 1, independently across loci. The initial frequency of the reference allele at each locus was drawn, independently across loci, from a uniform distribution on [*MAF*, 1 – *MAF*]; in our simulations, we chose a minimum minor allele frequency of *MAF* = 0.1. Since here we are interested in quantifying the mean squared effect size estimate, which is directionally affected by drift-based local LD that may not be present in our initial configuration, we allowed 150 generations of random mating before switching on assortative mating (only the final 20 generations of this random mating burn-in are displayed in Fig. 3). Assortative mating occurred for 19 generations, after which random mating occurred for a further 20 generations.

### Fig. 4. Cross-trait assortative mating for traits with different architectures

We again followed a similar procedure to that for Fig. 2 above, but now, while traits 1 and 2 have distinct genetic bases, the numbers of loci contributing variation to traits 1 and 2 are |*L*_1_| = 100 and |*L*_2_| = 1,000. Trait-1 loci are placed evenly along the physical genome, with trait-2 loci then evenly spaced among the trait-1 loci; we used the human linkage map for these simulations. At both trait-1 and trait-2 loci, the initial frequency of the focal allele was drawn from a uniform distribution on [*MAF*, 1 – *MAF*], with *MAF* = 0.1. At trait-2 loci, true effect size were randomly drawn from a normal distribution with mean zero and standard deviation 1; at trait-1 loci, true effect sizes were randomly drawn from a normal distribution with mean zero and standard deviation 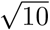, so that traits 1 and 2 have equal genic variances. After a burn-in of 150 generation of random mating, assortative mating was switched on. We performed a population GWAS at the end of the period of random mating and after 20 generations of assortative mating. These GWASs were performed across 1,000 replicate trials, with the effect size estimates then pooled across trials. From these, we estimated the densities of the absolute values of effect size estimates using Matlab’s kernel density estimator ksdensity, specifying that the support of the distributions be positive.

### Fig. 5. Population structure and admixture

We wished first to simulate a situation where two populations of size *N*/2 have been separated for a length of time such that the value of *F_ST_* between them is some predefined level (in our case, a mean *F_ST_* per locus of 0.1). To do so without having to run the full population dynamics of two allopatric populations for a prohibitively large number of generations, we simply assigned allele frequencies to achieve the desired level of *F_ST_*. We assumed 1,000 loci spread evenly over the physical genome. At each locus *l*, we chose an ‘ancestral’ frequency 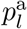 for the reference allele independently from a uniform distribution on [*MAF*, 1 – *MAF*], with *MAF* = 0.2. We then perturbed this allele frequency in populations 1 and 2 by independent draws from a normal distribution with mean 0 and variance 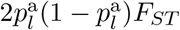; if a perturbed allele frequency fell below 0 or above 1, we set it to 0 or 1 respectively. The population dynamics described above, with monogamous mating pairs chosen randomly, were then run for 50 generations.

In generation 50, the two populations merge, forming an admixed population of size *N*. The same population dynamics, with monogamous mating pairs chosen randomly, were then run for a further 50 generations.

### Fig. 6. Stabilizing selection

To calculate the bias in GWAS effect size estimation caused by stabilizing selection, we must first calculate the harmonic mean recombination rate. We focus on humans, and consider only the autosomal genome. The set of loci underlying variation in the trait is *L*, which we apportion among the 22 autosomes according to their physical (bp) lengths (as reported in GRCh38.p11 of the human reference genome; https://www.ncbi.nlm.nih.gov/grc/human/data?asm=GRCh38.p11). For each chromosome, we spread its allotment of loci evenly over its sex-averaged genetic (cM) length, using the male and female linkage maps produced by Kong et al. (2010). (We use genetic lengths instead of physical lengths because, were we to spread loci evenly over the physical lengths of the chromosomes, some pairs of adjacent loci on some chromosomes might have a sex-averaged recombination fraction of 0, in which case the harmonic mean recombination rate would be undefined.) For each pair of linked loci, the recombination rate between them was estimated separately from the male and female genetic distance between them using Kosambi’s map function (Crow 1990). Pairs of loci on separate chromosomes have a recombination fraction of 1/2. With the sex-averaged recombination fraction *c_ll′_* thus calculated for every pair of loci (*l*, *l′*), the harmonic mean recombination fraction was calculated as 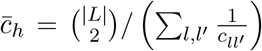, where 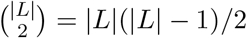 is the number of pairs of distinct loci in *L*.

Performing this calculation with |*L*| = 1,000 loci, we obtain an estimate of 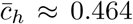 for human autosomes. Substituting this estimate into Appendix Eqs. (A.87) and (A.88) then defines the curves plotted in Figs. 6A and 6B respectively.

---

**Figure S1:**
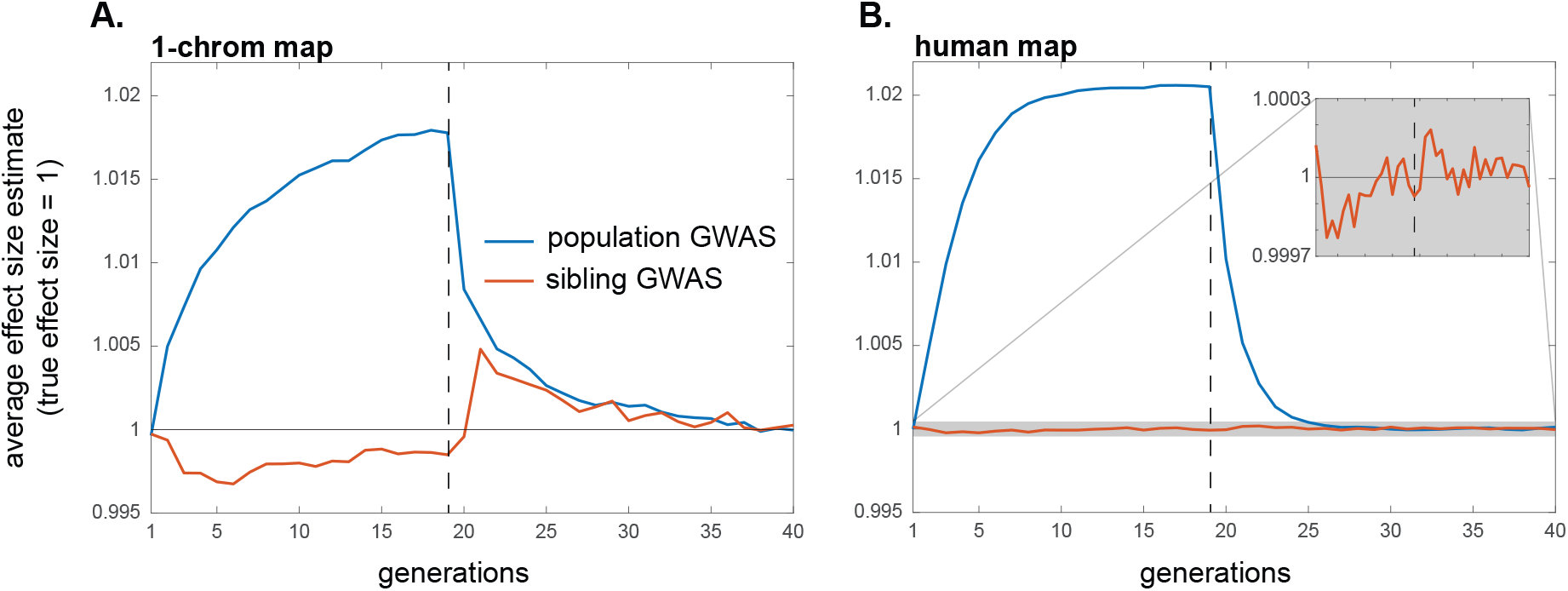
Cross-trait assortative mating influences effect size estimates at loci that affect the study trait, although this influence is second-order relative to that on effect size estimates at loci that do not affect the study trait but do affect the other trait involved in assortative mating (note the scale of the y-axis). Simulations are the same as in Fig. 2.

## A1 Genetic confounding in population and family-based GWAS designs

### A1.1 The model

Under the general additive model we have studied, an individual’s value for trait Y is

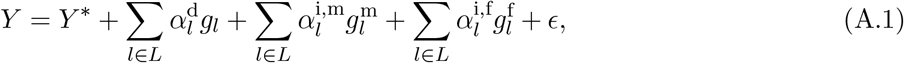

where *g_l_* is the number of focal alleles at locus *l* carried by the individual, 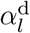 is the direct genetic effect on the trait value of the focal allele at *l* (which we assume to be positive, without loss of generality), 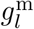 and 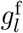 are the numbers of copies of the focal allele at locus *l* carried by the individual’s mother and father respectively, and 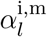 and 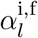 are the indirect genetic effects of the focal allele at *l* via the mother’s and father’s genotype respectively. *ϵ* is the environmental disturbance, with mean zero, and *Y*\* is the expected trait value of the offspring of parents who carry only trait-decreasing alleles.

It will be useful to expand Eq. (A.1) in terms of the individual’s and the individual’s parents’ maternally and paternally inherited genotypes:

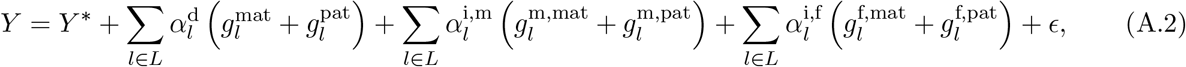

where 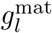 is the number of focal alleles at locus *l* that the individual inherited maternally, 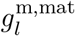 is the number of focal alleles at *l* that the individual’s mother inherited maternally, etc.

### A1.2 Population GWAS

If we perform a standard population GWAS at a genotyped locus λ, the estimated effect of the focal allele at λ on the trait Y is

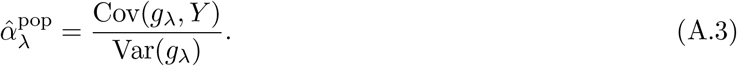

Here, Var(*g*_λ_) is the genotypic variance at λ among sampled individuals, equal to 2*p*_λ_(1 – *p*_λ_)(1 + *F*_λ_), where *p*_λ_ is the frequency of the focal allele at λ and *F*_λ_ is the coefficient of inbreeding at λ. For example, if λ is at Hardy-Weinberg equilibrium, then Var(*g*_λ_) = 2*p*_λ_(1 – *p*_λ_); if, instead, the population is divided into several populations, in each of which Hardy-Weinberg equilibrium obtains at λ but between which the frequency of the focal variant differs, then Var(*g*_λ_) = 2*p*_λ_(1 – *p*_λ_)(1 + *F*_*ST*,λ_), where *F*_*ST*,λ_ is the value of *F_ST_* at locus λ.

The covariance term in Eq. (A.3) expands out to

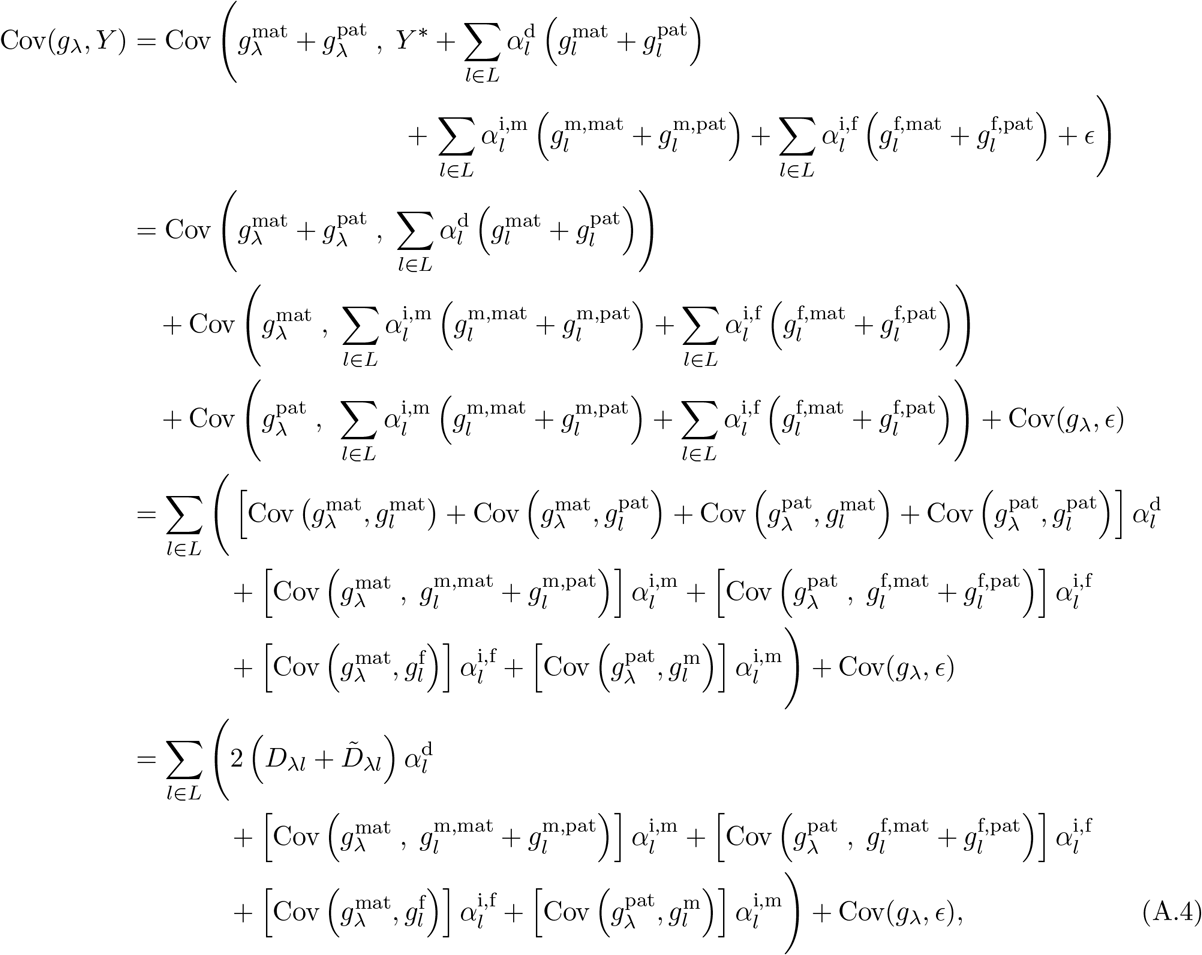

where *D*_λ*l*_ and 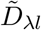 are the degrees of cis- and trans-linkage disequilibrium between the focal alleles at loci λ and *l* in the GWAS sample. Since 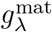 equals 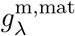 or 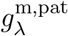 with equal probability, 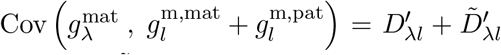, and similarly, 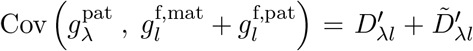 (here, 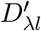 and 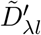 are the LDs in the parents of the sample, assumed to be equal across mothers and fathers). Since maternal transmission is independent of paternal genotype, and vice versa, 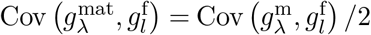 and 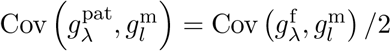. So

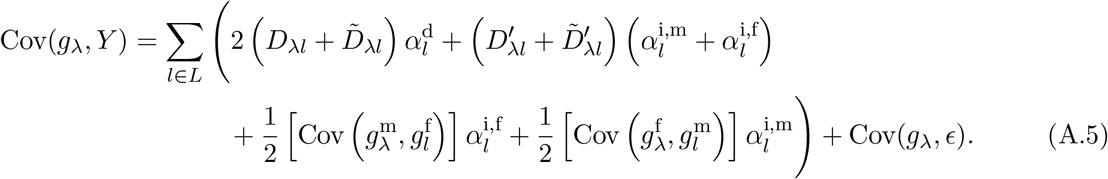

If 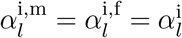, then

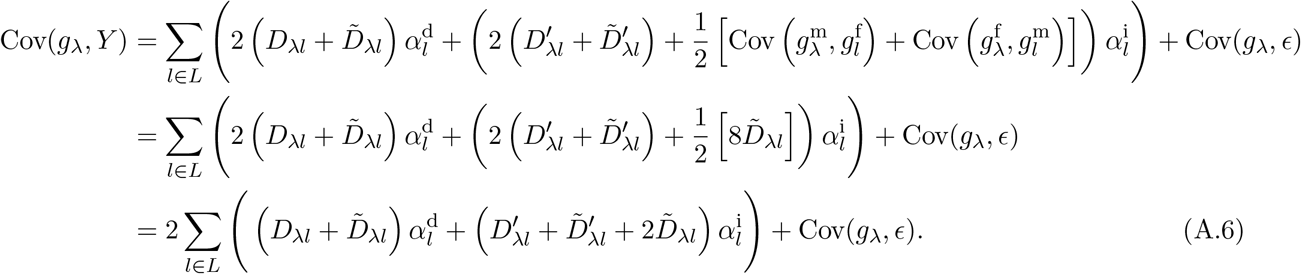

In the second line of Eq. (A.6), we have used the fact that covariances across parents translate to covariances across maternal and paternal genomes in the offspring. Note, however, that 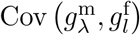 and 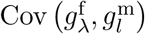 need not, in general be equal—e.g., they will not be so under sex-based cross-trait assortative mating—which is why we could not apply a similar simplification to Eq. (A.5).

Dividing Eq. (A.6) by Var(*g*_λ_), and recognizing that, for *l* ∈ *L*_local_, *c*_λ*l*_ ≈ 0, we recover Eq. (3) in the Main Text.

### A1.3 Sibling GWAS

Consider two full siblings. Let 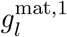 and 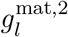 indicate whether sib 1 and sib 2 respectively inherited the focal (trait-increasing) allele from their mother at locus *l*. Let 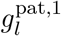 and 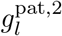 be analogous indicators for paternal transmission. Write 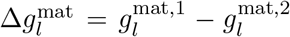 and 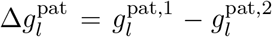. Since maternal and paternal transmission are independent, 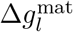 and 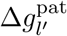 are independent for all pairs of loci *l* and *l′* (including *l* = *l′*). The difference in the two siblings’ genotypic values at locus *l* is 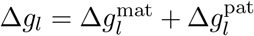. From Eq. (A.1), the difference in their trait values is

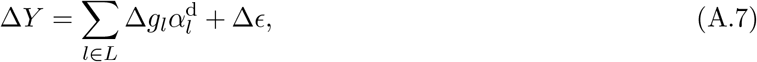

where Δ*ϵ* is the difference in the environmental disturbances experienced by the two siblings. Notice that the indirect effects cancel out of Eq. (A.7), since the parental genotypes are the same for the two siblings. So, in a sib-GWAS for trait Y, the estimated effect size at λ is

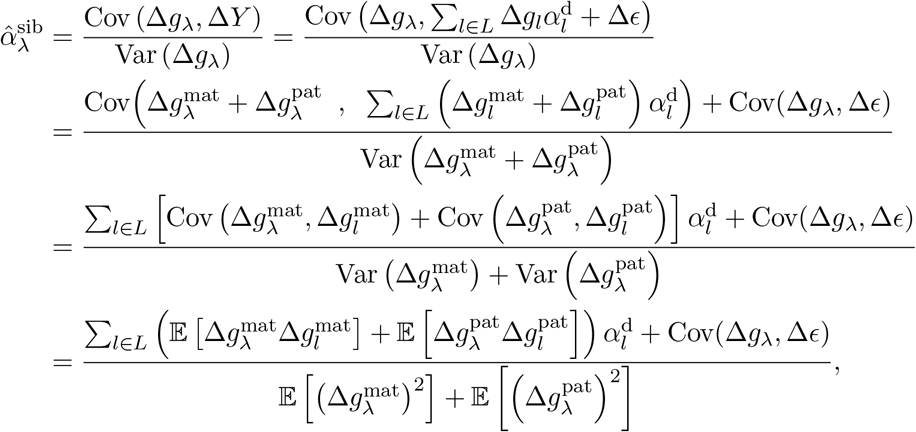

since 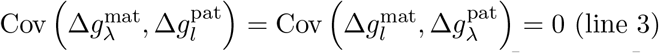 (line 3) and 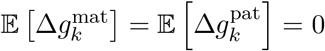 for all loci *k* (line 4). The denominator 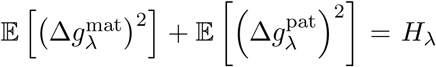, the fraction of parents in the family GWAS sample who are heterozygous at locus λ. The only non-zero contributions to 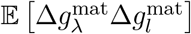 and 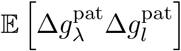 come from parents who are heterozygous at both λ and *l*. Such parents are either ‘coupling’ double-heterozygotes carrying the focal alleles at λ and *l* in coupling phase (i.e., inherited from the same parent), or ‘repulsion’ double-heterozygotes carrying the focal alleles at λ and *l* in repulsion phase (inherited from different parents). Among parents, let the fractions of coupling and repulsion double-hets for loci λ and *l* be 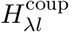 and 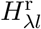 respectively. If the recombination rate between the loci is 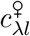 in females and 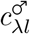 in males, then

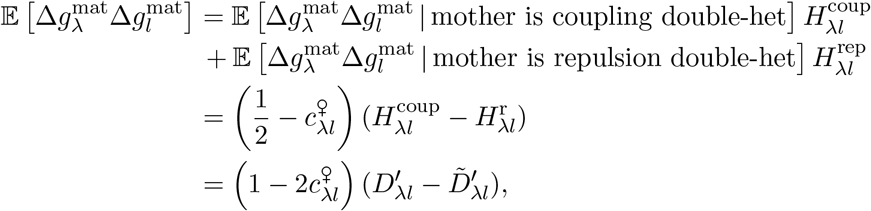

since 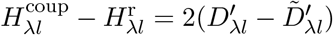, where 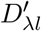 and 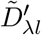 are the cis- and trans-LD between the focal/trait-increasing alleles at λ and *l* among parents.. Similarly,

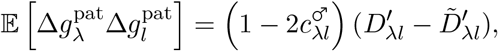

So

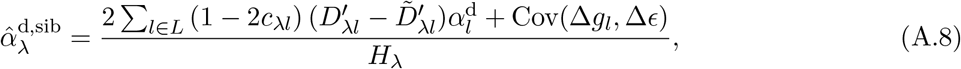

where *c*_λ*l*_ is the sex-averaged recombination fraction between λ and *l*. Since Cov(Δ*g_l_*, Δ*ϵ*) = 0, and recognizing that, for *l* ∈ *L*_local_, *c*_λ*l*_ ≈ 0 and 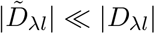 in expectation, we recover Eq. (7) in the Main Text.

### A1.4 Indirect effects: transmitted vs. untransmitted alleles

In Eq. (A.2), 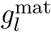 represents the allele that was transmitted maternally from among the set of maternal alleles 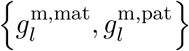. Thus, if the maternally transmitted allele was the grandmaternal allele (with probability 1/2, and in which case 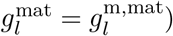), then the untransmitted allele at locus *l* is the grandpaternal allele, with genotypic value 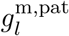. To make this distinction clear, we write 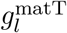 for the genotypic value of the maternally transmitted allele at locus *l*, and 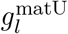 for the maternally untransmitted allele at locus *l*. Similarly, 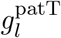 and 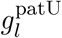 represent the paternally transmitted and untransmitted alleles at *l*. The transmitted and untransmitted genotypes are 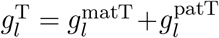 and 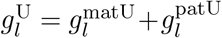 respectively.

#### Estimating direct effects

The regressions of the trait value on the transmitted and untransmitted genotypes are

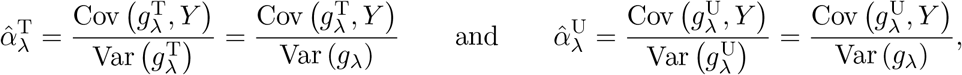

where we have used the fact that, since transmission at λ is random, 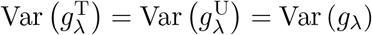. The estimate of the direct effect of the focal variant at λ is then

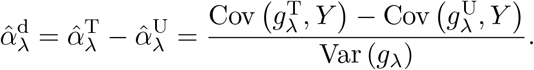

We have

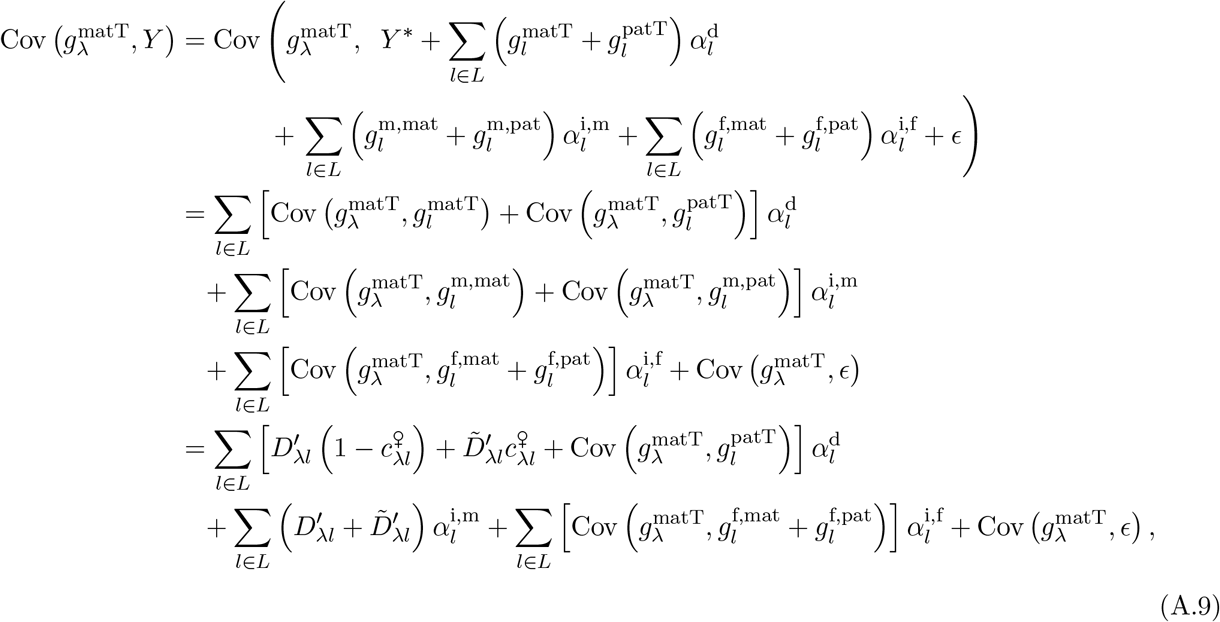

and

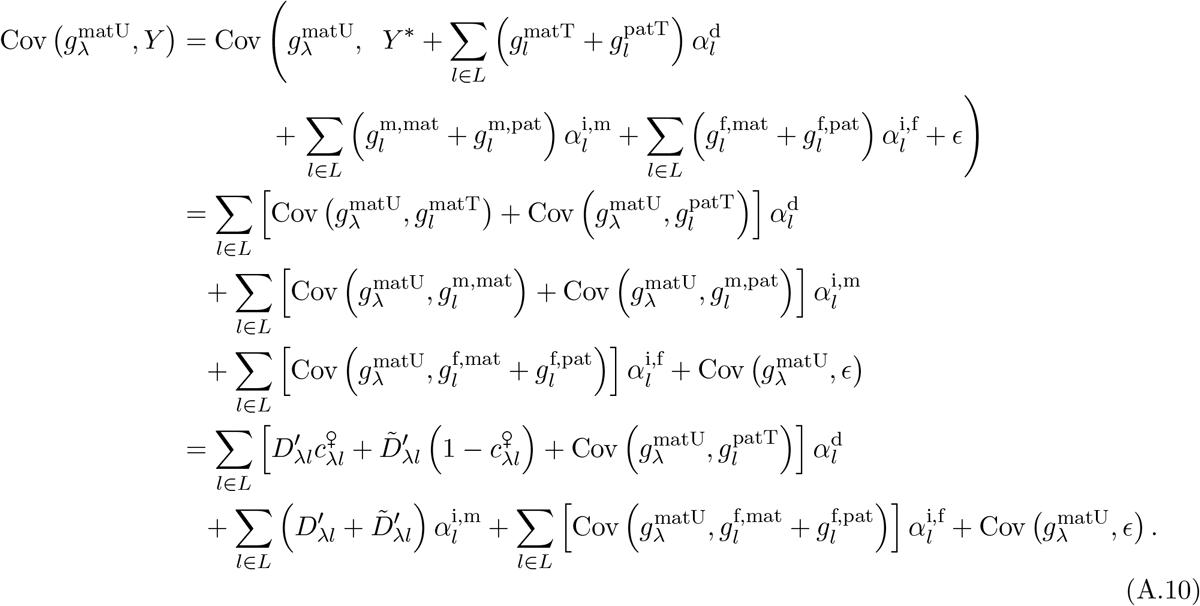

Since 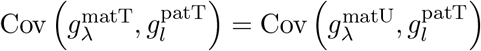 and 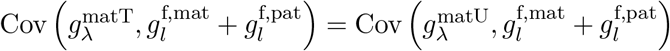,

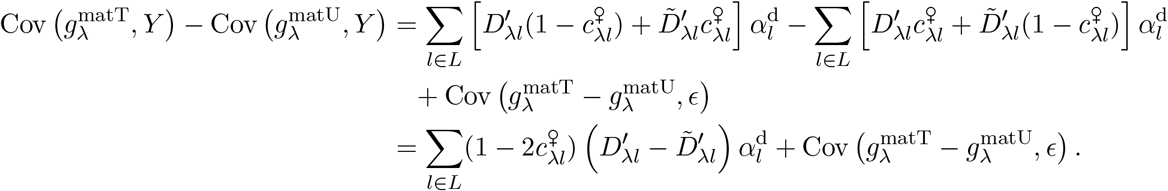

Similarly,

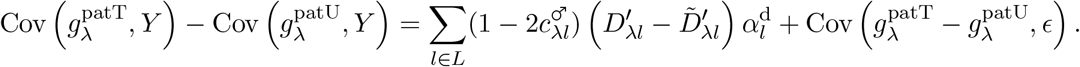

Since 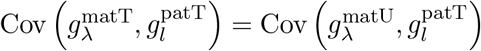 and 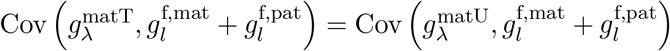,

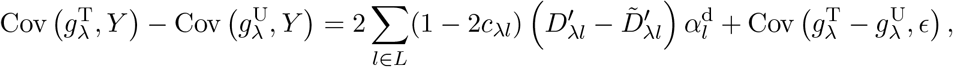

where *c*_λ*l*_ is the sex-averaged recombination fraction between λ and *l*. Therefore, the transmitted-untransmitted regression coefficient at locus λ is

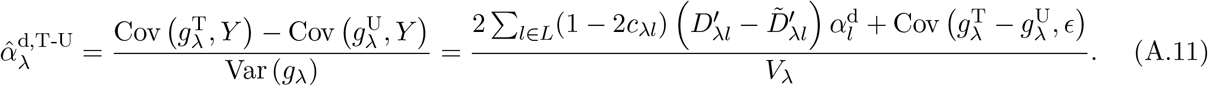

#### Estimating indirect effects

The coefficient in the regression of the trait value *Y* on the untransmitted genotype 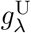 at locus λ, 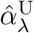, has sometimes been considered to provide an estimate of the indirect ‘family’ effect of the focal variant at 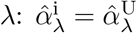. From Eq. (A.10) and its analog for the paternally untransmitted allele,

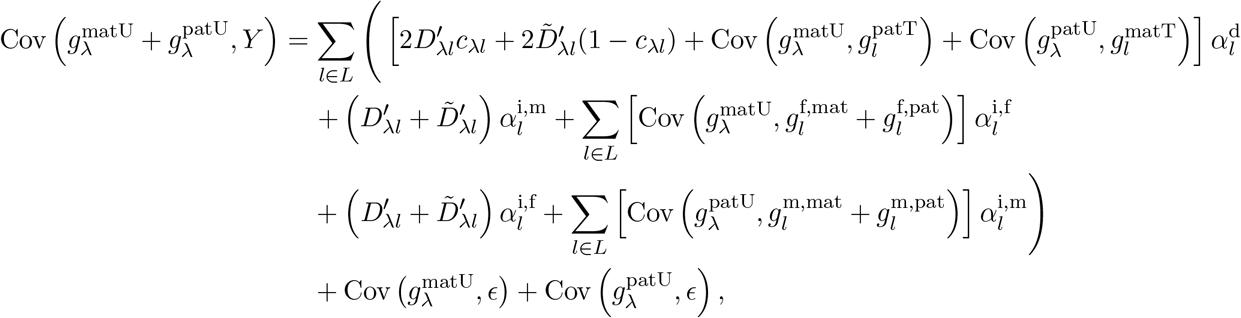

where *c*_λ*l*_ is the sex-averaged recombination fraction. In this expression,

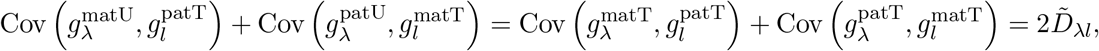

while

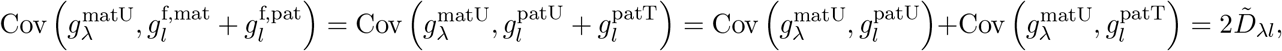

and, similarly, 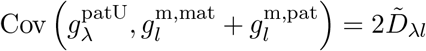. So

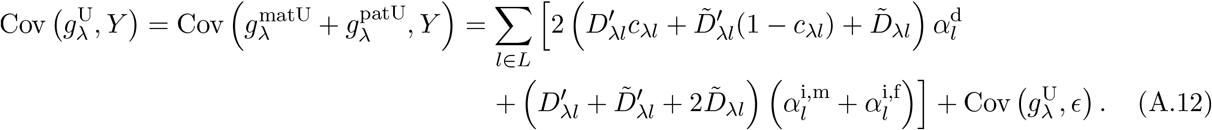

If we assume that indirect effects via the maternal and paternal families are equal 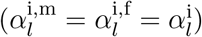, then Eq. (A.12) simplifies further to

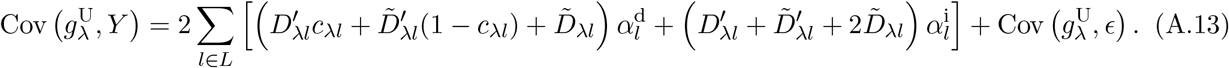

In this case, the estimate of the indirect effect of the focal allele at λ is

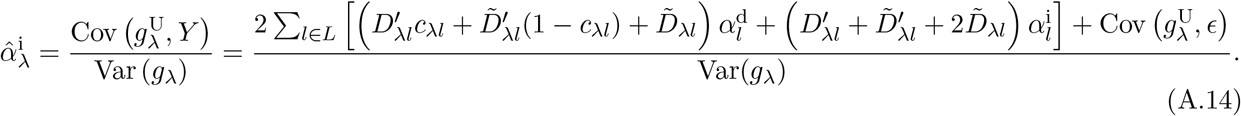

## A2 Polygenic scores and their phenotypic correlations

Suppose that we have estimated effect sizes 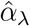 at a set of genotyped loci λ ∈ Λ using a population GWAS for trait 1. For each individual, we can then compute a polygenic score:

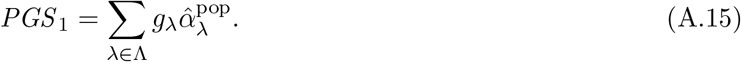

PGSs are often treated as predictions of individuals’ genetic values for traits. In this regard, we might therefore be interested in the covariance across the population between the PGS for a trait and individuals’ values for that trait: Cov(*PGS*_1_, *Y*_1_). Additionally, if PGSs are treated as predictions of genetic values of traits, then we might be interested in how the PGS calculated for one trait covaries with the value of another trait: Cov(*PGS*_1_, *Y*_2_). Such covariances might be informative of genetic correlations between traits, or pleiotropy of the alleles underlying genetic variation in the traits. We focus on the two-trait covariance, since it nests the single-trait covariance as a special case. If the total set of loci causally underlying variation in traits 1 and 2 is *L*, then the population covariance between the PGS for trait 1 and the value of trait 2 is

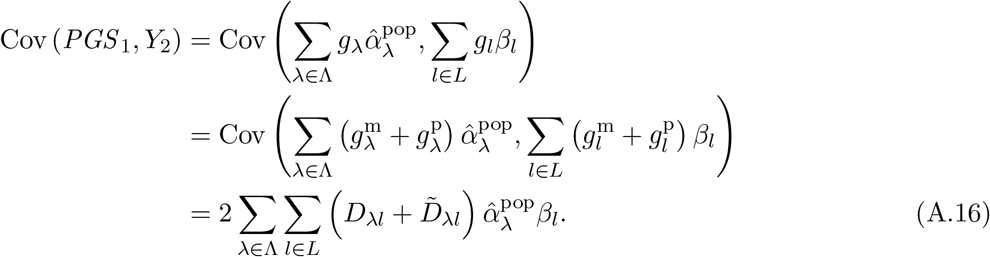

The effect size estimates from the population GWAS for trait 1 are

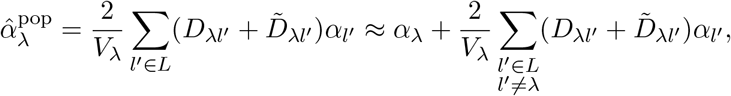

and so Eq. (A.16) is, in general,

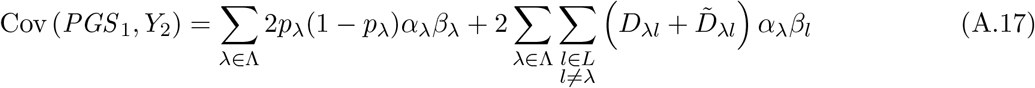

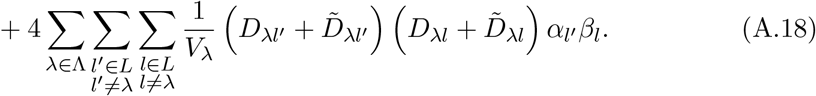

In a family-based study, we might instead be interested in the covariance between siblings’ differences in the trait-1 population PGS and their differences in trait 2. We can write this covariance in our model as

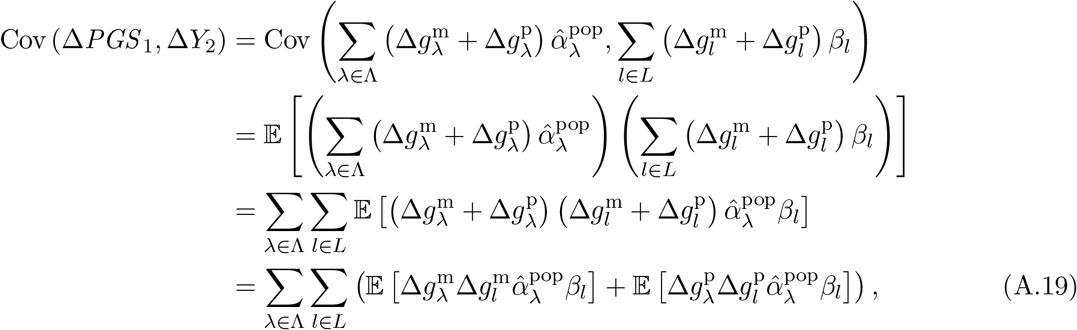

since maternal and paternal transmission are conditionally independent. Focusing on maternal transmission, and writing 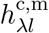 and 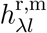 for the events that the mother is respectively a coupling and a repulsion heterozygote at loci λ and *l*, with 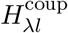 and 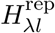 their associated probabilities (which are assumed to be the same for mothers and fathers),

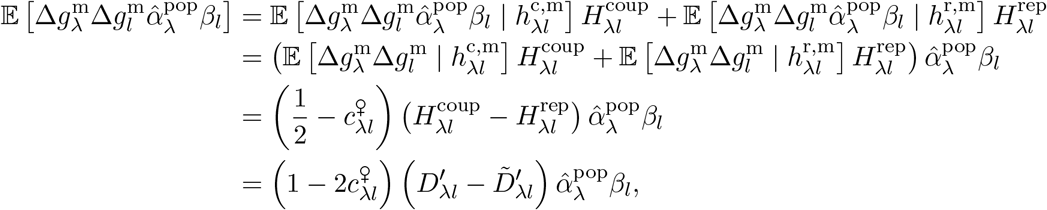

with 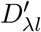 and 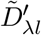 measured in the parents. Similarly,

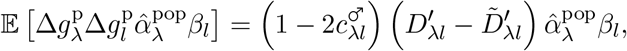

and so Eq. (A.19) becomes

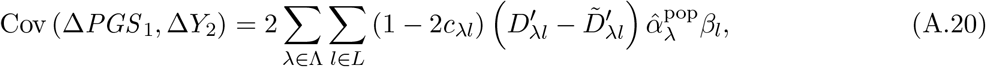

where *c*_λ*l*_ is the sex-averaged recombination fraction between λ and *l*.

Before we substitute the population GWAS estimates 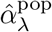 into Eq. (A.20), it is worth considering what value this expression would take if effect sizes were correctly estimated at every study locus, 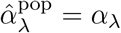. In this case, Eq. (A.20) becomes

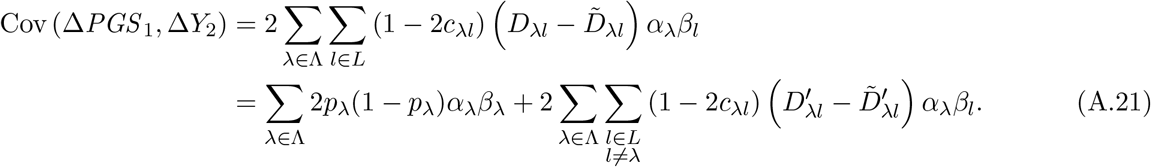

If the two traits are distinct, then the first term in Eq. (A.21) is the genic covariance of traits 1 and 2 across the set of study loci (more precisely, tagged locally by the study loci), and reflects systematic pleiotropy at these loci; this term would, for example, be positive if alleles tend to have same-direction effects on traits 1 and 2. If we were studying only one trait, then *α*_λ_ = *β*_λ_, and the first term would be the genic variance of the trait across study loci, 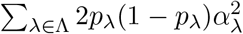. The second term in Eq. (A.21) is an effect of linkage disequilibria between study loci and the loci that are causal for trait 2; these LDs are absorbed by the PGS because the PGS is a sum across loci. In the absence of such LDs, or in cases where the cis- and trans-LDs are equal so that 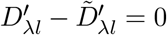, Eq. (A.21) would equal the genic variance in the single-trait case and the genic covariance in the two-trait case.

The effect size estimates from a population GWAS are in fact

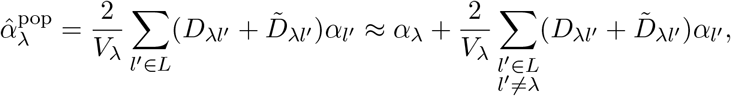

*D*_λ*l′*_ and 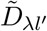 are measured in the sample. We assume these to be equal to the values in parents in the family-based GWAS, 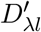 and 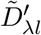, and so the value taken by Eq. (A.20) is

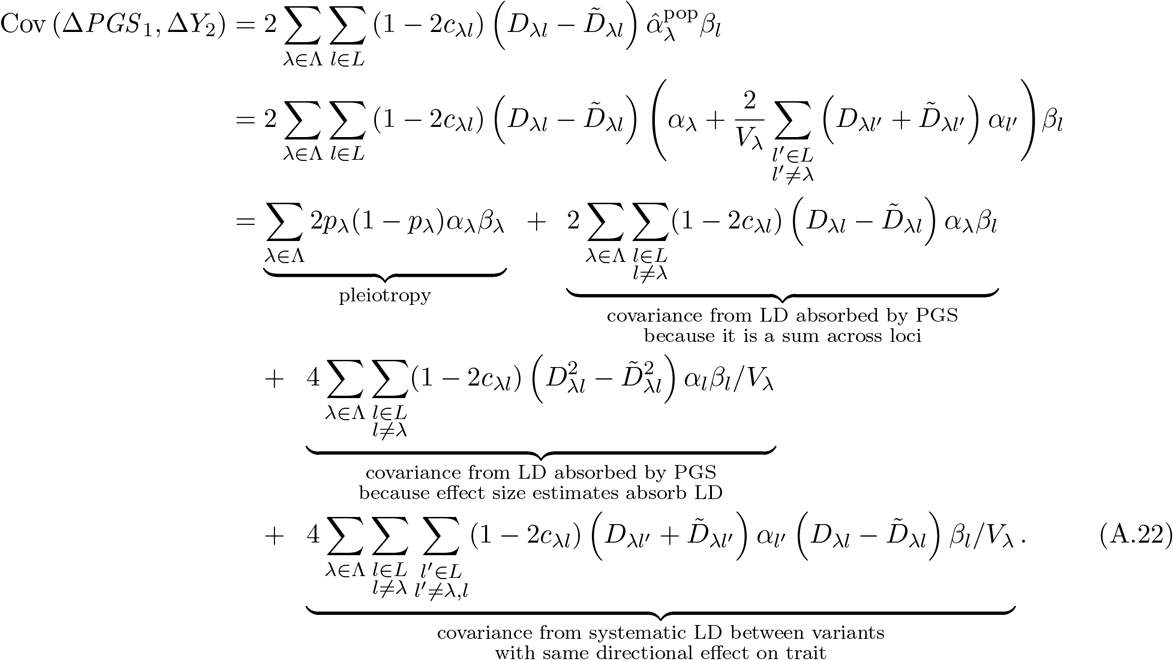

In the absence of genetic confounding 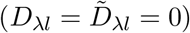 or, more generally, if genetic stratification is such that the cis- and trans-LDs are equal 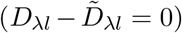, then Eq. (A.22) simplifies to the SNP-tagged genic covariance between traits 1 and 2:

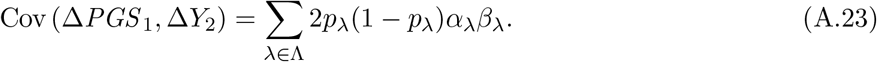

If traits 1 and 2 are the same, then this is simply the SNP-tagged genic variance of the trait: 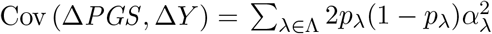.

Eq. (A.22) simplifies somewhat if we focus on a single trait (*α_l_* = *β_l_*) and assume that there is no trans-LD 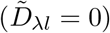; in this case,

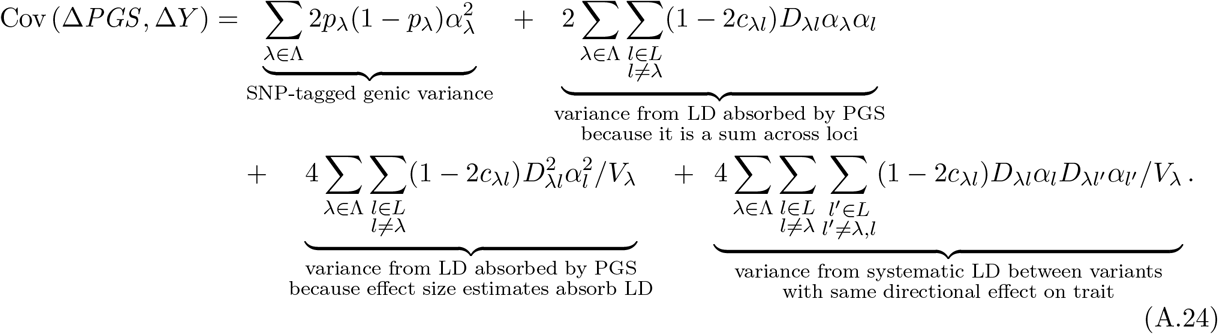

## A3 Sources of genetic confounding

The calculations above reveal that genetic confounds in GWAS designs can depend on long-range LD in the sample and among parents of the sample. Here, we consider several possible sources of long-range LD.

### A3.1 Assortative mating

If there is a constant correlation among mates for their values of two traits, then a genetic equilibrium will eventually be achieved. In this equilibrium, for any pair of loci *l* and *l′*, the trans-LD 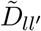 will be constant. Call this constant value 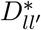, and suppose that the recombination fraction between the loci is *c_ll′_*. With 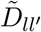 constant across generations, the balance of its conversion into cis-LD (at rate *c_ll′_*, per generation) and the destruction of cis-LD by recombination (at rate *c_ll′_*, per generation) will result in an equilibrium level of cis-LD equal to the degree of trans-LD: 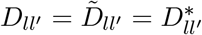 (e.g., Crow and Felsenstein 1968).

The value of 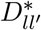, will, in general, depend in a complicated way on the strength of effects of *l* and *l′* on the traits upon which assortative mating is based and on the linkage relations of these loci to one another and to other causal loci. However, while it is therefore difficult to calculate the individual equilibrium LD terms 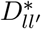, we can in some cases calculate weighted sums of these terms across locus pairs.

Let the set of loci that influence one or both traits be *L*, and let *α_l_* be the effect size of the focal variant at locus *l* on trait 1 and *β_l_* its effect on trait 2 (the analyses below also apply to same-trait assortative mating, setting *α_l_* = *β_l_*). Recall the notation 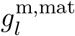 and 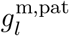 for a mother’s maternally and paternally inherited genotype at locus *l*, with 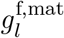 and 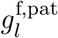 a father’s analogs. The mother’s breeding value for trait 1 is

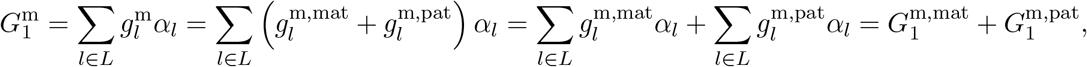

and, similarly, her breeding value for trait 2 is

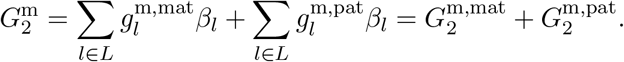

The father’s breeding values for the two traits are

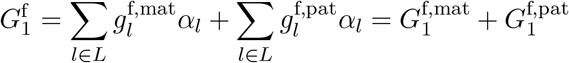

and

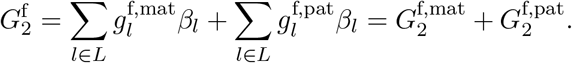

We assume that individual trait values equal the breeding values plus environmental disturbances that are uncorrelated with the breeding values:

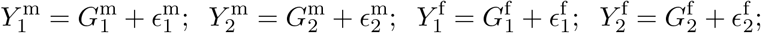

where

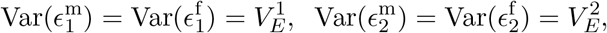

and

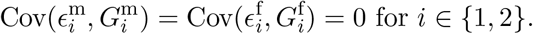

#### A3.1.1 Same-trait assortative mating, or cross-trait assortative mating that is symmetric with respect to sex

We first consider the case where the strength of assortative mating between two traits, as measured by their correlation coefficient across mating pairs, is equal in the female-male and male-female directions. Notice that this scenario covers same-trait assortative mating. In the case of cross-trait assortative mating, it could occur if assortative mating arises by mechanisms other than direct female (or male) mating preferences.

We assume that there is a constant correlation *ρ* among mating pairs for their phenotypic values of traits 1 and 2. In equilibrium, this will translate to a constant correlation *ρ_G_* between their breeding values as well (e.g., Felsenstein 1981). To calculate *ρ_G_*, we first note that, because assortative mating is based on phenotypic values and not breeding values per se, if we know the phenotypes of a pair of mates, we obtain no further information about the similarity of their breeding values; that is,

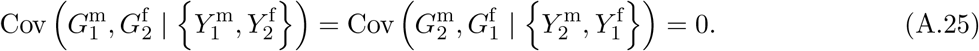

For the same reason, if we know the phenotypic values of two mates, then the trait-2 value of the male does not offer any information on the female’s trait-1 breeding value beyond that already offered by the female’s trait-1 phenotype, and vice versa; that is,

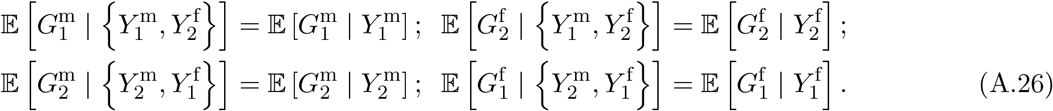

If *Y*_1_ and *G*_1_, and similarly *Y*_2_ and *G*_2_, are bivariate normal, then

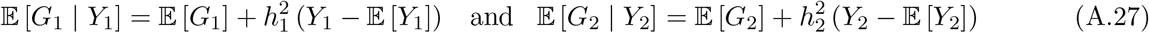

where 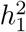 and 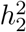 are the heritabilities of traits 1 and 2, respectively.

From the law of total covariance,

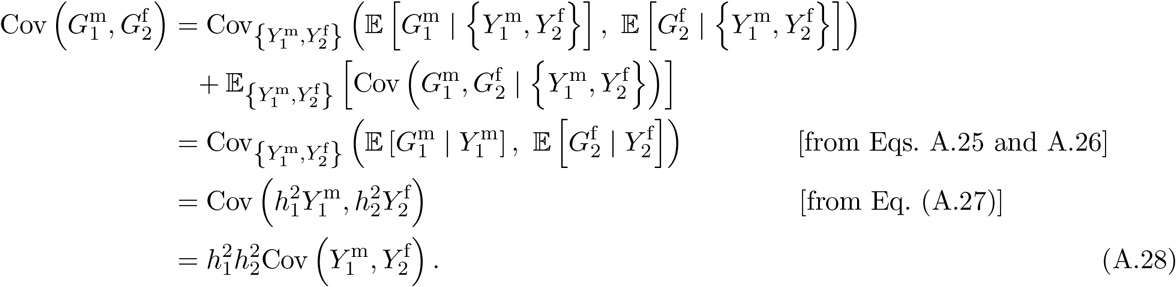

Similarly, 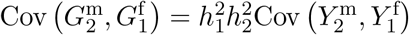.

Let *V*^1^ and *V*^2^ be the phenotypic variances of traits 1 and 2, and 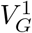 and 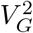 their additive genetic variances, assumed to be the same across the sexes. Given the calculations above, the correlation among mates for their breeding values of traits 1 and 2, *ρ_G_*, can be written

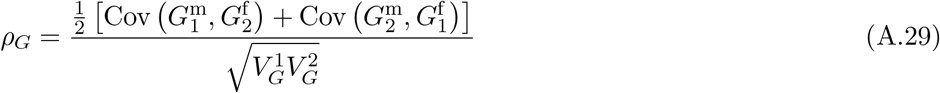

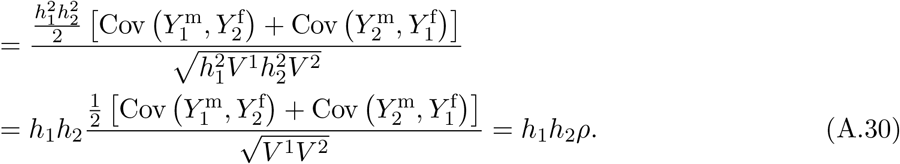

When traits 1 and 2 are the same, we have *ρ_G_* = *h*^2^*ρ*, a standard result (e.g., Wright 1921; Felsenstein 1981).

Expanding the numerator of Eq. (A.29),

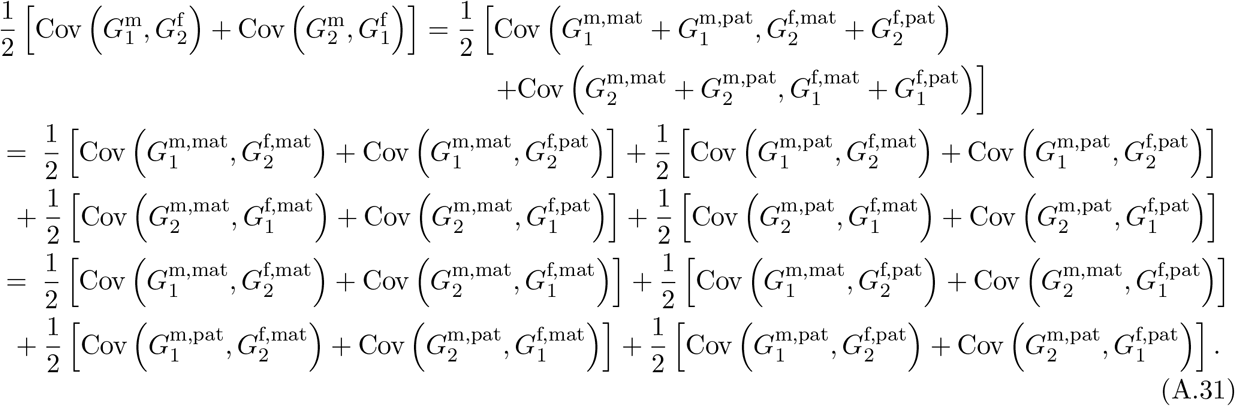

But

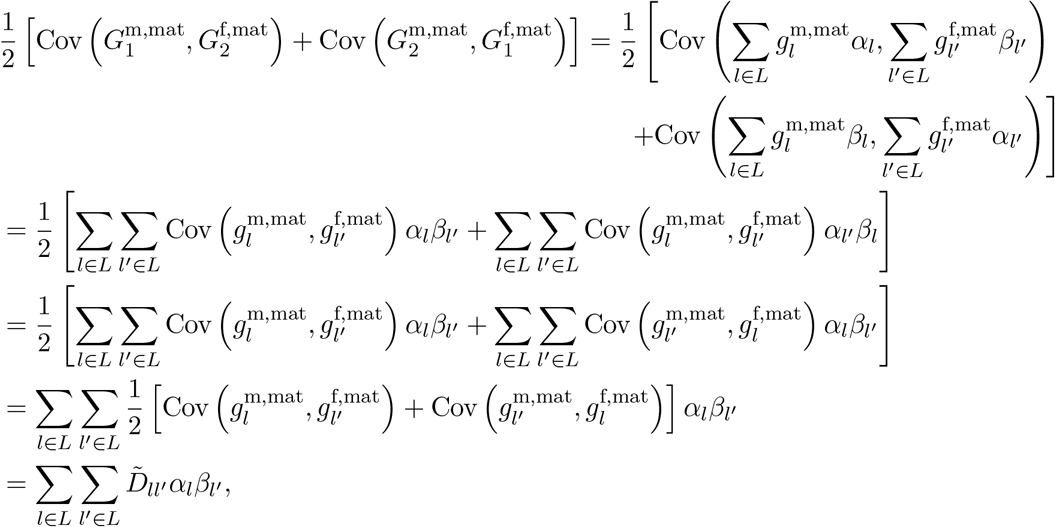

since grandmaternal and grandpaternal alleles are transmitted to the offspring with equal probability, independently across maternal and paternal transmission. The three additional terms in Eq. (A.31) likewise each amount to 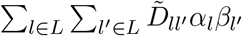, and so

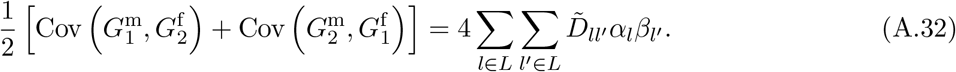

Noting that the trans-covariance at a given locus 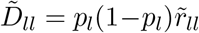, where 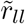 is the within-locus correlation (equal to the inbreeding coefficient at the locus), we can split Eq. (A.32) into within- and between-locus terms:

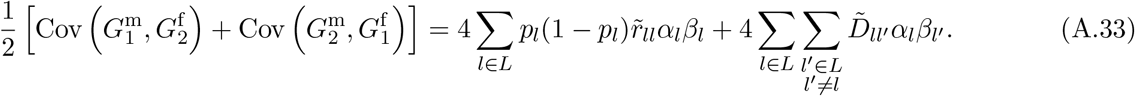

In the denominator of Eq. (A.29),

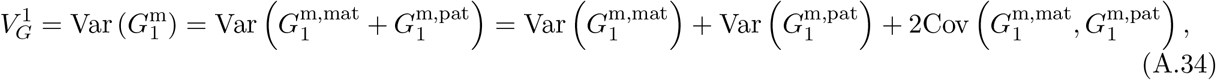

Expanding the first term,

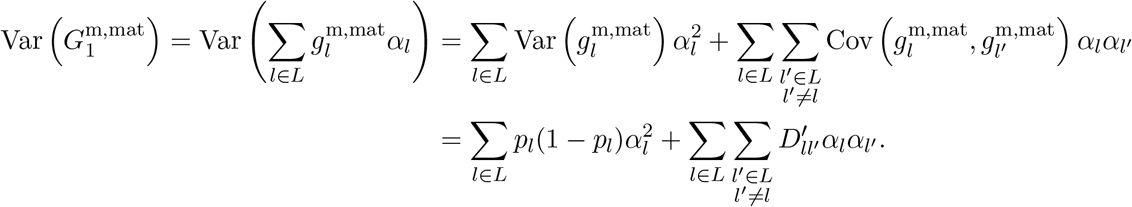

Similarly, the second term is

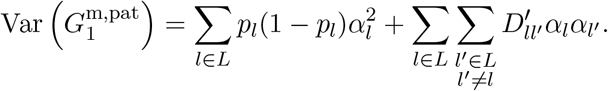

The third, covariance term in Eq. (A.34) is

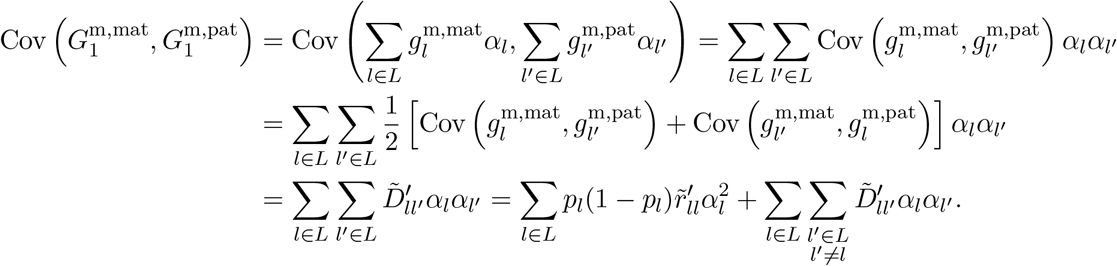

Putting these together in Eq. (A.34),

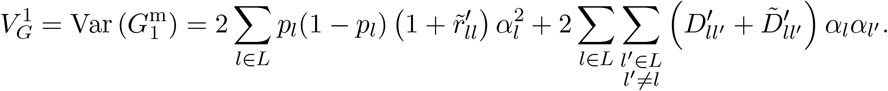

Similarly,

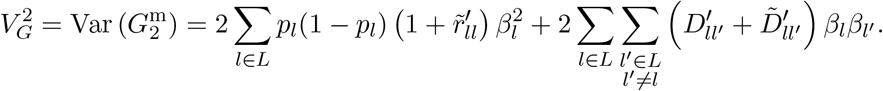

In equilibrium, 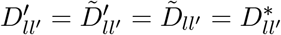, for *l* ≠ *l′*, and 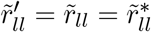, so

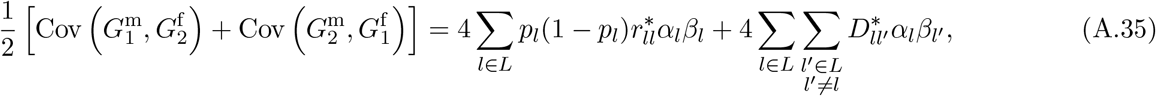

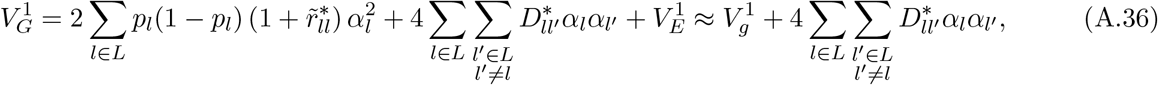

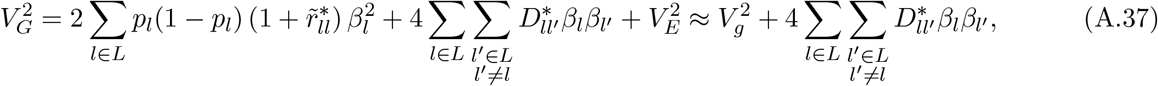

where 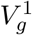 and 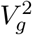 are the genic variances of traits 1 and 2, and the approximations come from the fact that, under assortative mating for a polygenic trait, the sum of the ~|*L*|^2^ cross-locus trans-LD terms 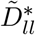, dominates the sum of the |*L*| within-locus trans-LD terms 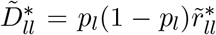 (Crow and Kimura 1970, Ch. 4). Eq. (A.29) in equilibrium is therefore

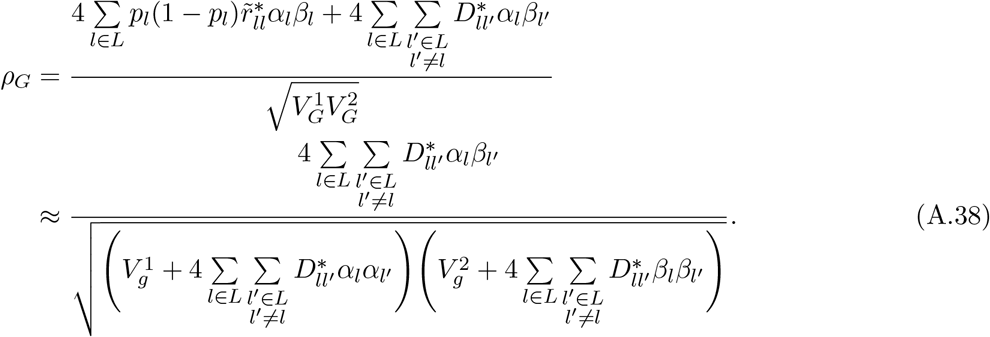

We now consider some special cases.

##### Same-trait assortative mating with equal effect sizes

In the case of same-trait assortative mating, *α_l_* = *β_l_*, so Eq. (A.38) simplifies to

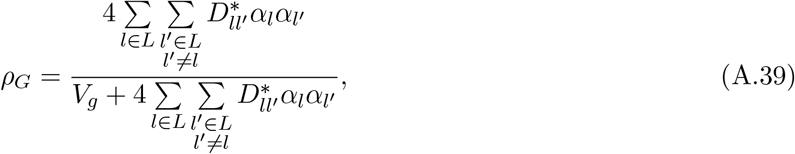

from which

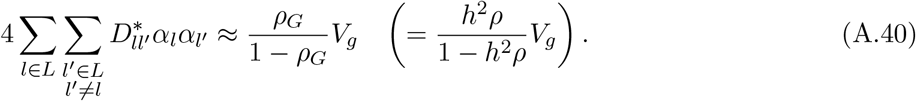

Since, in equilibrium, 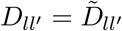, this expression can also be written

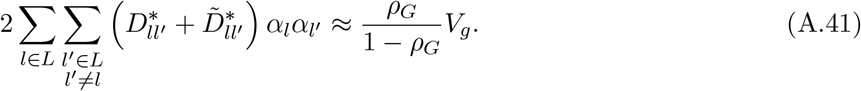

Because the additive genetic variance 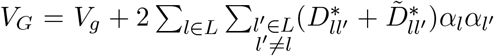, Eq. (A.41) can also be written

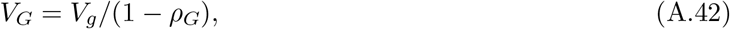

which is a classic result (e.g., Wright 1921; Crow and Kimura 1970, Ch. 4).

If we make the further assumption that effect sizes are the same across loci (*α_l_* = *α* for all *l* ∈ *L*), then Eq. (A.41) becomes

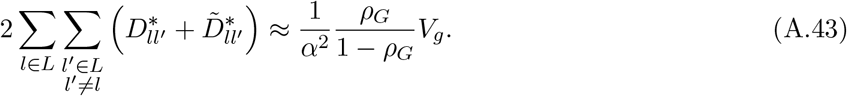

In a population association study at locus *l*, assuming no indirect effects and no sources of genetic confounding other than assortative mating, the effect size estimate is

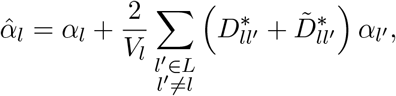

so that the proportionate bias in the effect size estimate at *l* is

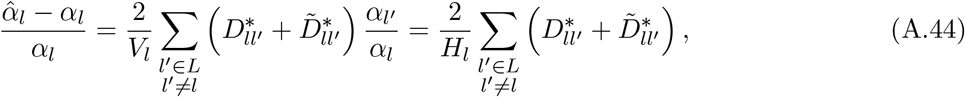

since *α_l′_*, = *α_l_* by assumption and *V_l_* ≈ *H_l_* = 2*p_l_*(1 – *p_l_*) because assortative mating does not substantially increase within-locus homozygosity (Crow and Kimura 1970, Ch. 4). The average proportionate bias across loci is then

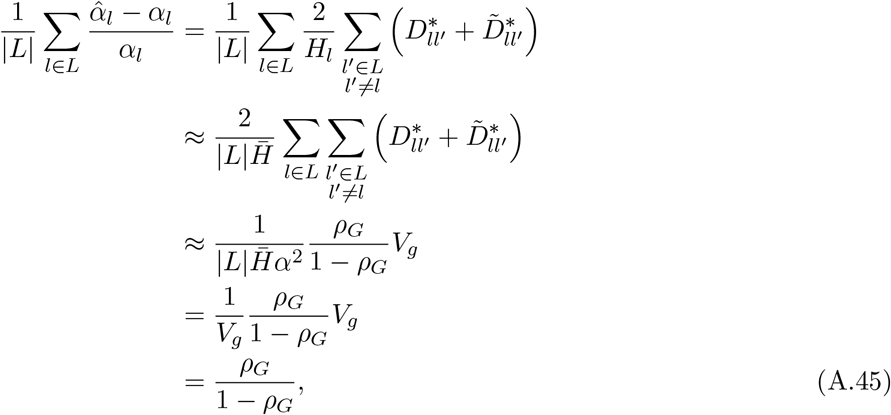

where we have used Eq. (A.43) and have assumed that minor allele frequencies do not differ widely across loci. Since *ρ_G_* = *h*^2^*ρ*, where *ρ* is the phenotypic correlation among mates and *h*^2^ = *V_G_*/*V_P_* is the heritability of the trait, Eq. (A.45) can also be written

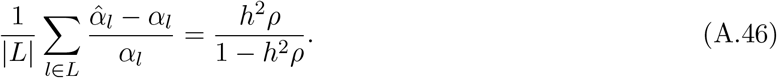

##### Sex-symmetric cross-trait assortative mating with distinct genetic bases and equal effect sizes

In the case of cross-trait assortative mating, if the sets of loci underlying the two traits, *L*_1_ and *L*_2_, are distinct, then *α*_l_ ≠ 0 ⟹ *β_l_* = 0 and *β_l_* ≠ 0 ⟹ *α_l_* = 0. In this case, Eq. (A.38) becomes

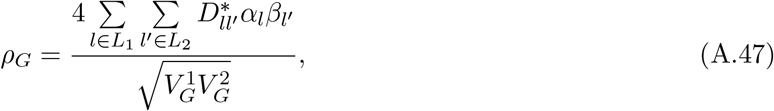

from which

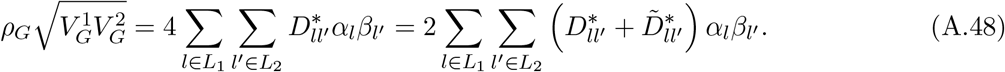

Because assortative mating is cross-trait, the LDs that assortative mating induces across *L*_1_ and *L*_2_ will dominate the second-order LDs induced within *L*_1_ and within *L*_2_. Therefore, 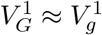 and 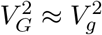.

The effect size estimate at a locus *l* ∈ *L*_1_ in a population GWAS on trait 2 is

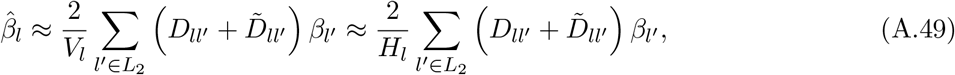

while the true effect size *β_l_* is zero, since *l* ∉ *L*_2_. In equilibrium, the average effect size estimate, and thus the average deviation of these estimates from the true values, is therefore

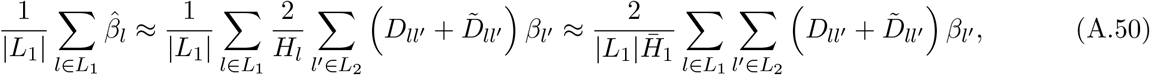

where we have assumed that minor allele frequencies are not very different across *L*_1_ (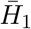 is the average heterozygosity in *L*_1_). If we further assume that effect sizes at causal loci are equal for each trait (*α_l_* = *α* for all *l* ∈ *L*_1_ and *β_l′_*, = *β* for all *l′* ∈ *L*_2_), then Eq. (A.50) can be written

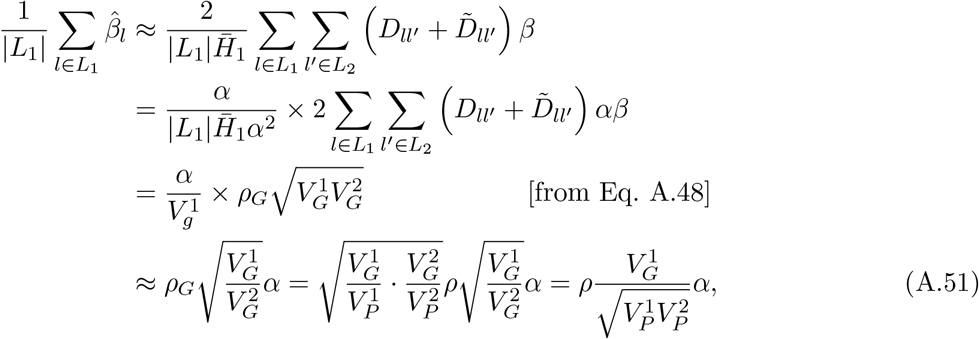

recalling from Eq. (A.30) that 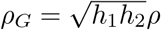.

In the further special case where both the genetic and the phenotypic variances of the two traits are equal, then so are the heritabilities of the two traits. In this case, Eq. (A.51) simplifies to

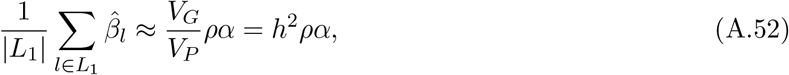

where *h*^2^ is the common heritability of the two traits.

##### Sex-symmetric cross-trait assortative mating for traits with different genetic architectures

Eq. (A.52) reveals an interesting role for genetic architecture in the bias that cross-trait assortative mating can generate in population association studies performed at non-causal loci. Suppose, as we did in deriving Eq. (A.52), that the two traits on which assortative mating is based have the same genetic and phenotypic variances, *V_G_* and *V*, and therefore also the same heritabilities, *h*^2^. We shall make the further assumption that the traits have the same genic variance, *V_g_*. Assume further that the sets of loci underlying traits 1 and 2, *L*_1_ and *L*_2_, have similar mean heterozygosities 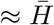. Normalize the effect size sizes at loci causal for trait 2 to *β* = 1, so that the traits’ common genic variance is 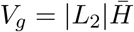.

Suppose that we now perform a population GWAS for trait 2. At loci that are causal for trait 2 (*l* ∈ *L*_2_), we will estimate effect sizes accurately: 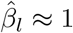 (there will be a small positive second-order bias, of order *ρ*^2^, since the locus *l* ∈ *L*_2_ comes into positive LD with loci *l′* ∈ *L*_1_, which in turn have come into positive LD with loci *l″* ∈ *L*_2_).

At loci that are causal for trait 1 (*l* ∈ *L*_1_), and which therefore have no effect on trait 2, we will estimate effect sizes on average as given by Eq. (A.52): 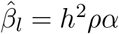.

How does the number of loci underlying variation in trait 1, |*L*_1_|, affect this biased estimate of their effect on trait 2? For the genic variance of trait 1 to be the same as that of trait 2, 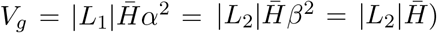, and so we must have *α*^2^ = |*L*_2_ |/|*L*_1_|. Substituting this into the average effect size estimate at non-causal loci, 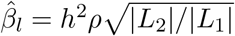.

So, the average effect size estimate at causal loci *l* ∈ *L*_2_ is 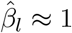, while the average effect size estimate at non-causal loci *l* ∈ *L*_1_ is 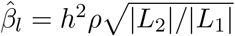. How do these two quantities compare? If the number of loci underlying the two traits is the same, *L*_1_ = *L*_2_, and effect size estimates at non-causal loci are smaller than those at causal loci by a factor of about *h*^2^*ρ*. However, if there are more loci underlying trait 2 than underlying trait 1—i.e., if trait 1 has a more concentrated genetic architecture *L*_1_ < *L*_2_—then the effect size estimates at non-causal loci will be closer to those at causal loci. Indeed, if trait 1 has a sufficiently concentrated architecture relative to trait 2, specifically, if *L*_1_ < *h*^4^*ρ*^2^L_2_, then the effect size estimates at non-causal loci will, on average, be larger in magnitude than effect size estimates at causal loci.

More generally, the calculations above suggest that, in a more realistic scenario where effect sizes vary across loci, the trait-2 GWAS distribution of magnitudes of effect size estimates at trait-1 loci (non-causal) will overlap more with the distribution of magnitudes of effect size estimates at trait-2 loci (causal) if the genetic architecture of trait 1 is more concentrated (Fig. 4). This will lead to a greater number of trait 1 loci being identified as statistically significantly associated with trait 2 in the trait-2 GWAS.

#### A3.1.2 Cross-trait assortative mating that is asymmetric with respect to sex

We now consider the case where the strength of assortative mating between two traits, as measured by their correlation coefficient across mating pairs, is not equal in the female-male and male-female directions. This is clearest in the case of an active mate preference exhibited by one sex for some phenotype exhibited by the other sex.

To study this case, we make several simplifying assumptions. First, we assume that the genetic bases of variation in the two traits are distinct: *α*_l_ ≠ 0 ⇔ *β*_l_ = 0. Second we assume that there is only one active direction of assortative mating: female trait 1 and male trait 2. That is, conditional on the mother’s breeding value for trait 1 and the father’s breeding value for trait 2, there is no correlation between the mother’s breeding value for trait 2 and the father’s breeding value for trait 1:

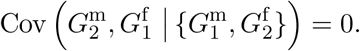

Suppose that there is a constant correlation *ρ_G_* between mothers’ breeding values for trait 1 and fathers’ breeding values for trait 2:

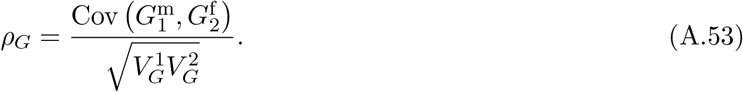

To study the genetic consequences of this assortment, we need to know the average bi-directional correlation among mates for traits 1 and 2 (Eq. A.29). Since traits 1 and 2 will come into a positive genetic correlation via assortative mating of female trait 1 and male trait 2, there will also be a positive covariance between mothers’ breeding values for trait 2 and fathers’ breeding values for trait 1, which we can express using the law of total covariance:

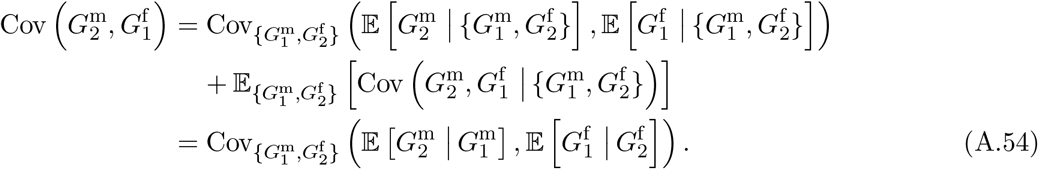

If 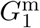 and 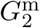 are bivariate normal (more generally, if 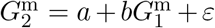, with 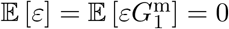), then

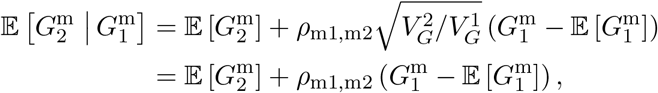

where 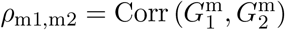 is the genetic correlation between traits 1 and 2 in mothers, and where we have assumed that the two traits have equal variance. Similarly, if 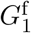 and 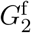 are bivariate normal, then

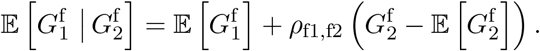

Substituting these expressions into Eq. (A.54),

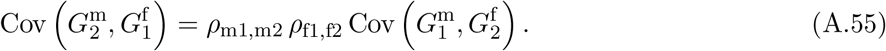

But, in our case, *ρ*_m1,m2_ = *ρ*_f1,f2_, the common value of which we shall call *ρ*_12_, and so the average bi-directional correlation is

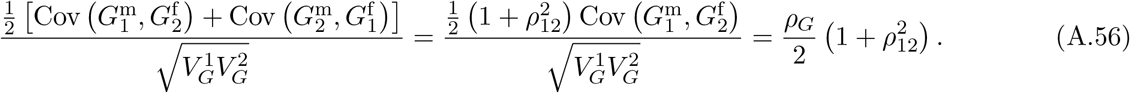

Given this value, the calculations of the effect of assortative mating on the weighted sums of cis- and trans-covariances, and thus on the additive genetic variance, proceed as for the case of symmetric assortative mating above.

Assuming the genetic bases of the two traits to be distinct, we may substitute the average bi-directional correlation, 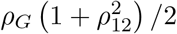, into Eq. (A.48) to find

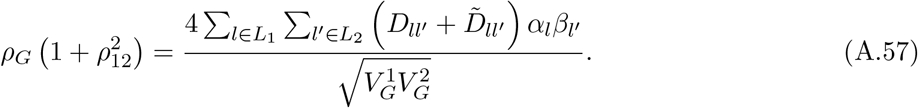

But

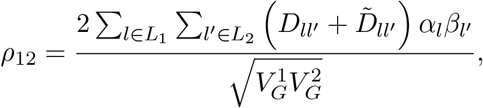

and so Eq. (A.57) can be written as the quadratic equation 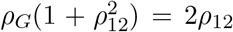, the relevant solution to which is 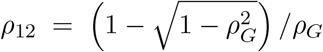. If *ρ_G_* is small, we use the first-order Taylor approximation 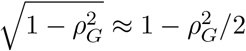 to find

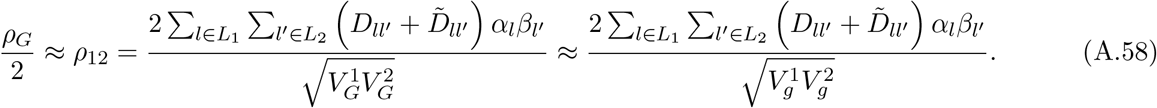

In the particular scenario we have simulated in Fig. 2, 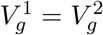, *α_l_* = 1 for all *l* ∈ *L*_1_, and *β_l_* = 1 for all *l* ∈ *L*_2_, so Eq. (A.58) further simplifies to

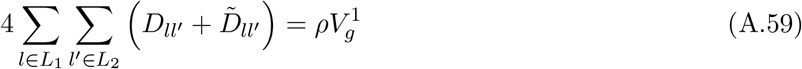

In a population association study for trait 2 performed at a locus *l* ∈ *L*_1_ (so that *β_l_* = 0),

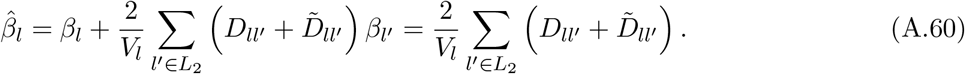

Across loci in *L*_1_, the average estimate is

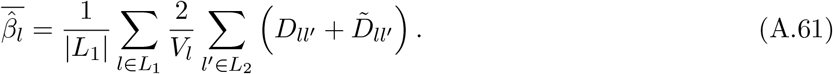

In our simulations, *p_l_* ≈ 1/2 for all *l* so that *V_l_* ≈ 2*p_l_*(1 – *p_l_*) = 1/2, and |*L*_1_| = |*L*_2_| = 500, so 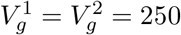. Under this configuration,

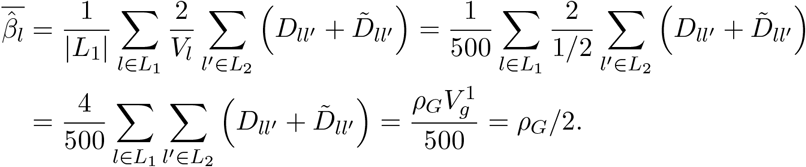

The trait we simulated is genetic, with heritability 1, and so *ρ_G_* = *ρ*, the phenotypic correlation among mates. We chose a strength of assortative mating of *ρ* = 0.2, and so, in equilibrium, the average effect size estimate at non-causal loci should be approximately 0.1, which is indeed the case in Fig. 2.

##### Sex-asymmetric cross-trait assortative mating for traits with different genetic architectures

For the case where the numbers of loci underlying traits 1 and 2 differ, and noting that the ‘effective’ correlation among mates in the sex-asymmetric case is approximately half that in the sex-symmetric case (Eq. A.58), we can perform a similar back-of-the-envelope calculation as in the sex-symmetric cross-trait assortative mating case above to find that, when effect sizes are constant across trait-1 loci and constant across trait-2 loci (though differing across traits 1 and 2), the effect size estimates at trait-1 (non-causal) loci in a trait-2 population GWAS is, on average, a fraction 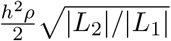 of the estimates at trait-2 (causal) loci.

Thus, more generally, when the number of loci underlying trait 1 is small relative to the number of loci underlying trait 2, the distribution of magnitudes of effect size estimates at trait-1 loci in a trait-2 GWAS can overlap substantially with the distribution of magnitudes of effect size estimates at trait-2 loci (Fig. 4), causing variants at these non-causal trait-1 loci to show up as significant in the trait-2 GWAS.

#### A3.2 Population structure

In the model we have considered, with results displayed in Fig. 5, there are initially two isolated populations of equal size. The frequency of the focal variant at locus *l* is 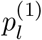 in population 1 and 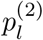 in population 2, so that its overall frequency is 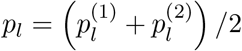. A population GWAS at locus λ returns an effect size estimate

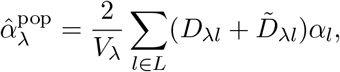

where *D*_λ*l*_ and 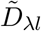 are calculated across both populations and are generally nonzero because of allele frequency differences between the two populations at loci λ and *l* (Nei and Li 1973). In our case,

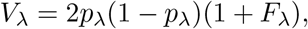

and

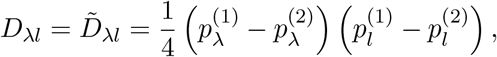

so

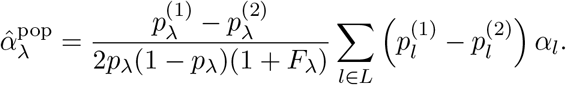

Squaring this and multiplying by 2_*p*λ_(1 – *p*_λ_),

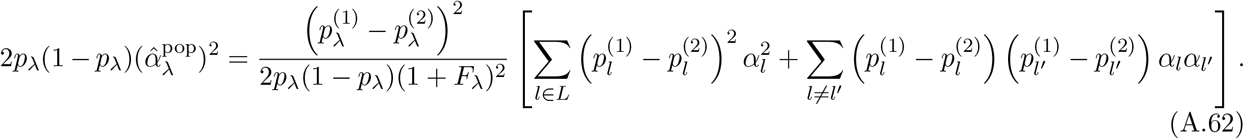

##### Neutral allele frequency divergence

If allele frequency divergence between the two populations is neutral, frequency changes at different loci are independent of one another and of effect sizes, so the second term in square brackets above is zero in expectation. In addition, because Hardy-Weinberg equilibrium obtains within each population, non-zero expected values of *F*_λ_ derive only from allele frequency differences between the populations, so that *F*_λ_ = *F*_*ST*,λ_ in expectation. Therefore,

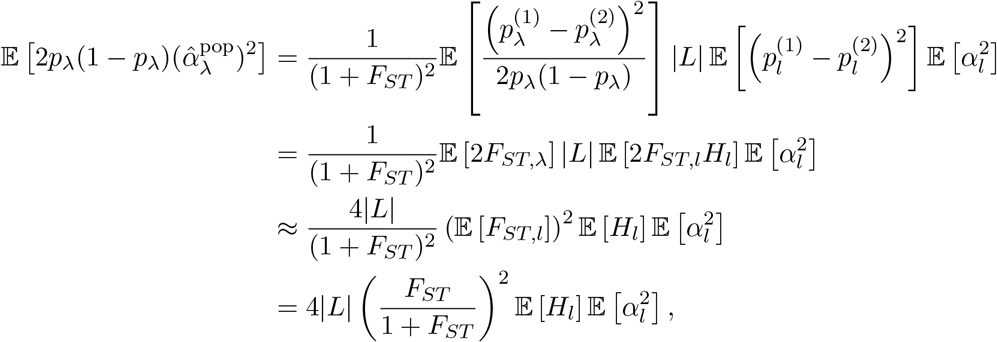

where *H_l_* = 2*p_l_*(1 – *p_l_*). If the ancestral allele frequency at *l* was 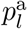, then 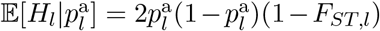, and so 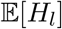 is calculated using the law of iterated expectations by averaging this quantity over the ancestral distribution of allele frequencies: 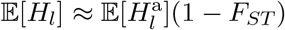, where 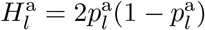. So

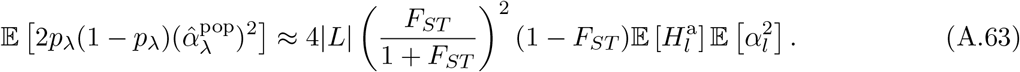

##### Selection and phenotype-biased migration

Above, in calculating the mean heterozygosity-weighted value of 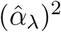 under neutral frequency divergence between populations, we assumed that in Eq. (A.62) the second term in the square brackets was zero, i.e., that the effect-size-signed population allele frequency difference was uncorrelated across loci. Howevever, when selection or phenotype-biased migration acts, this will no longer be true. For example, if higher genetic values of the trait were favoured in population 1 relative to population 2, then selection will on average have driven a mean shift such that 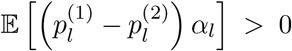. This in turn will drive systematic positive covariances between terms 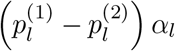 and 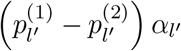, and as these covariances are summed over all pairs of loci in Eq. (A.62), the resulting inflation of the average squared effect size estimate (and other genome-wide summaries) could be quantitatively substantial.

##### More general population stratification

Given a sample of *N* individuals, the sample cis-LD between two markers λ and *l* can be written generally as

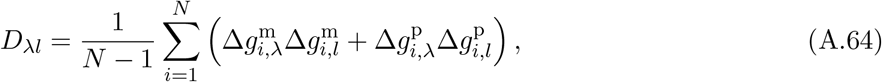

where 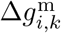 and 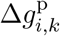 are the deviations of individual *i*’s maternal and paternal focal allele count at locus *k* from their mean frequencies. The trans-LD between λ and *l* is

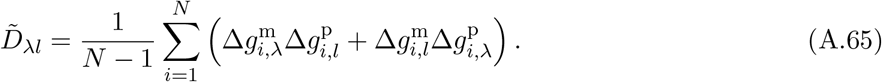

These cis- and trans-LD terms are equal only if

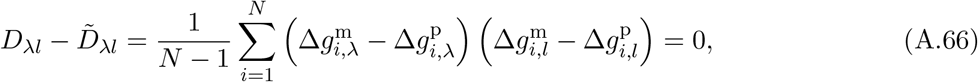

i.e., if the maternal and paternal alleles at the one locus are exchangeable with respect to deviations of the allelic state at the other locus.

We might often be concerned with stratification along some specific axis of variation in our sample. Call this axis *v*, with every individual having a value along *v*, with mean zero across individuals (for example, in our two population case above, the vector *v* could be 1 for population 1 and −1 for population 2). The covariance of the maternal allele at locus *l* with the vector *v* is proportional to 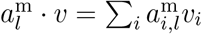. So the contribution of LD along this axis to the difference in cis- and trans-LD is

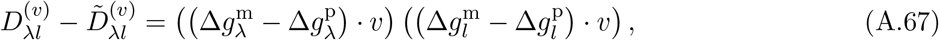

which is zero only if the maternal and paternal genotypes at the two loci are exchangeable with respect to each other along the axis *v*.

#### A3.3 Admixture

Suppose that two previously isolated populations admix in proportions *A* and 1 – A, with subsequent random mating in the admixed population. Following the notation in the Section A3.2 above, before admixture, the frequency of the focal variant at locus *l* was 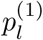 in population 1 and 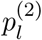 in population 2, so that its overall frequency in the admixed population is 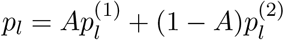.

When the two populations admix, trans-LD between all pairs of loci disappears in expectation, owing to random mating in the admixed population: 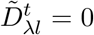 for any pairs of loci λ and *l* and for any number of generations *t* after admixture. However, cis-associations between alleles that were more prevalent in one ancestral population than in the other will be retained as cis-LD in the admixed population until these associations are eroded by recombination. The initial degree of cis-LD between loci λ and *l* in the admixed population is

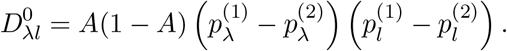

When *t* generations have elapsed since admixture, this cis-LD will have been eroded by recombination to

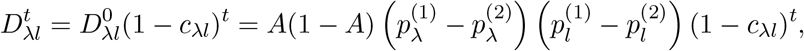

where *c_λl_* is the sex-averaged recombination rate between λ and l. Therefore, *t* generations after admixture, a population association study at λ returns an effect size estimate

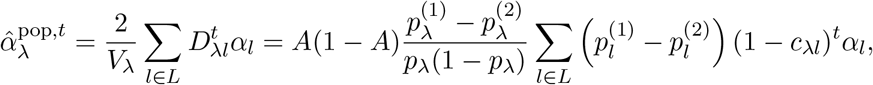

while a sibling-based association study at λ returns

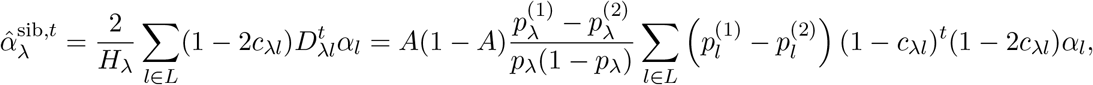

where we have substituted *V*_λ_ = *H*_λ_ = 2*p*_λ_(1 – *p*_λ_) owing to random mating in the admixed population. Squaring the population estimate and multiplying by 2*p*_λ_(1 – *p*_λ_),

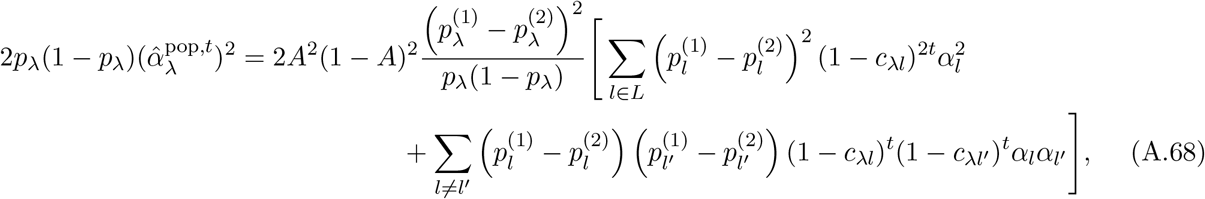

while the heterozygosity-weighted squared sibling effect size is

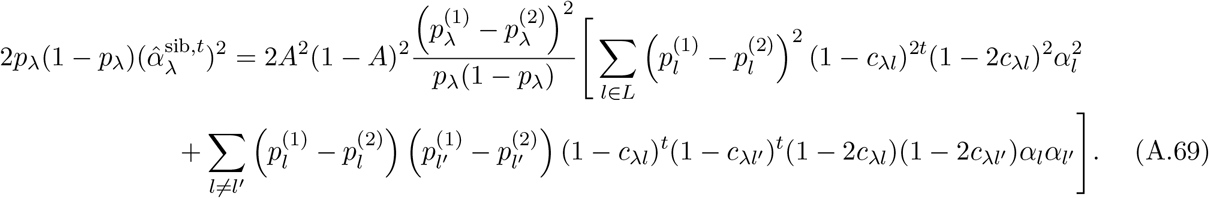

##### Neutral allele frequency divergence

If allele frequency divergence between the two populations was neutral, then frequency changes at different loci are independent of one another, of effect sizes, and of recombination rates (assuming the loci are sufficiently far apart), so the second terms in square brackets in Eqs. (A.68) above is zero in expectation, so that

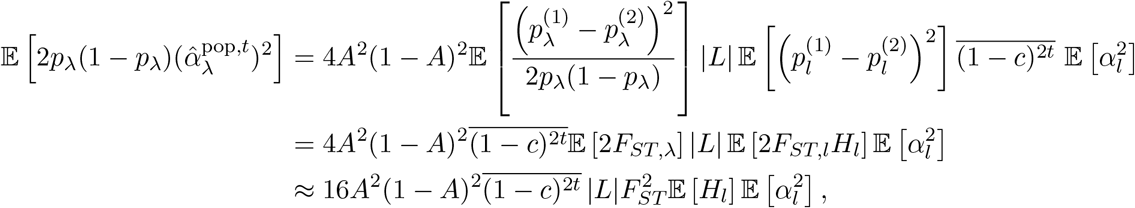

where 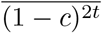 is the average value of 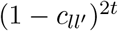 taken across all pairs of loci *l*, *l*′.

Similarly, under drift in the ancestral populations, the average squared sibling-based effect size estimate can be simplified to

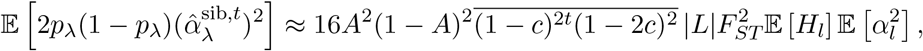

where 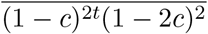 is the average value of 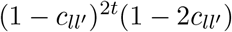 taken across all pairs of loci *l*, *l*′.

##### Selection and phenotype-biased migration

As in the case of population structure, selection and phenotype-biased migration in the ancestral populations can drive systematic positive covariances between the terms 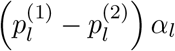 and 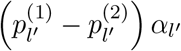 in Eqs. (A.68) and (A.69) above, so that the second terms in square brackets in these equations do not cancel in expectation as they did under neutral divergence between the ancestral populations. Again, as these covariances are summed over all pairs of loci in Eqs. (A.68) and (A.69), the resulting inflation of the average squared effect size estimate and other genome-wide summaries could be substantial.

#### A3.4 Stabilizing selection

We consider the model of Bulmer (1971, 1974), in which a very large number of loci contribute variation to a trait under stabilizing selection. We assume that the distribution of trait values is centered on the optimal value *Y*\*, and that the relative fitness of an individual with trait value *Y* is exp (−(*Y* – *Y*\*)^2^/2*V_S_*), where *V_S_*, the width or ‘variance’ of this gaussian selection function, governs the strength of stabilizing selection, with larger *V_S_* values implying weaker selection. Under this model, selection acts to reduce the phenotypic variation each generation; if the trait value is normally distributed with variance *V_P_*, then selection reduces the within-generation phenotypic variance by an amount

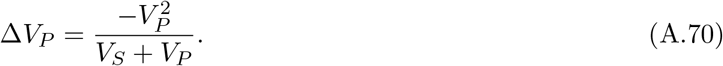

How much of this reduction carries over to the offspring generation then depends on the heritability of the trait.

Owing to the large number of loci in this model, the buildup of LD among them occurs on a faster timescale than the change in allele frequencies at individual loci. Assuming the loci to have equal effect sizes, Bulmer (1974) showed that the overall reduction in the phenotypic variance due to stabilizing selection, *d*, rapidly approaches a quasi-equilibrium value that approximately satisifes

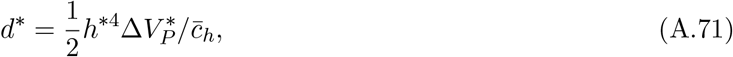

where *h*\*^2^ is the heritability of the trait in this equilibrium and 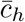 is the harmonic mean of the recombination rates amongst all pairs of loci. On this rapid timescale, the reduction in variance is due to LD among the loci underlying the trait; in fact,

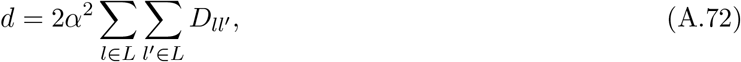

where *α* is the common per-locus effect size and 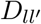 is defined with respect to the trait-increasing alleles at *l* and *l*′. The individual linkage disequilibria 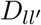, in expectation, are proportional to the inverse recombination rates 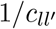. Writing

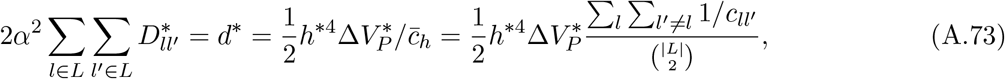

where 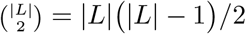 is the number of pairs of distinct loci in *L*, it is apparent that

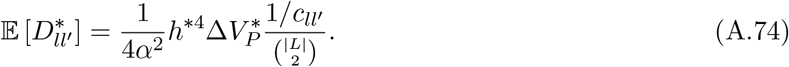

Henceforth we deal only with equilibriuj quantities and therefore drop the star superscript for neatness. The phenotypic variance *V_P_* can be written *V_P_* = *V_G_* + *V_E_* = *V_g_* + *d* + *V_E_*, where *V_G_* is the additive genetic variance, *V_g_* is the genic variance, and *V_E_* is the variance due to the environment. Eqs. (A.70) and (A.71), together with the definition of heritability *h*^2^ = *V_G_/V_P_*, define a quadratic equation in *d*:

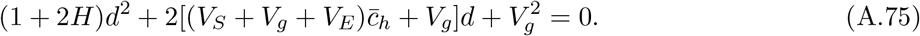

Eq. (A.75) matches Eq. (10) in Bulmer (1974), with Bulmer’s parameter *c* replaced by 1/2*V_S_*. For ease of reference in what follows, we write Eq. (A.75) in the standard form *ad*^2^ + *bd* + *c* = 0. The roots are

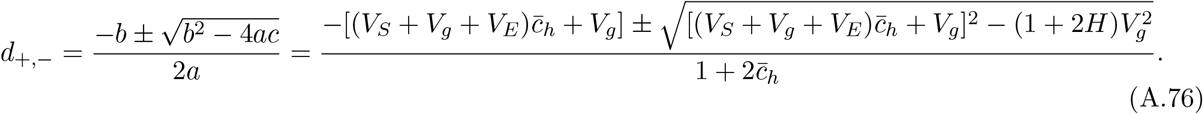

To see which of these roots is the relevant one, we first note that the roots are both real, since the requirement for this is

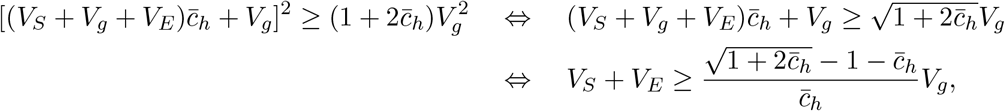

and 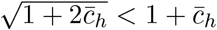 for 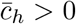, while *V_S_* + *V_E_* > 0. Furthermore, since *b* > 0 and 4*ac* > 0, both roots are in fact negative, with *d*_−_ < *d*_+_ < 0. Now note that

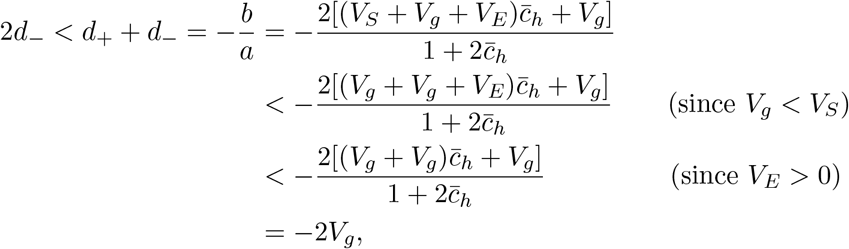

i.e., *V_g_* + *d*_−_ < 0. But then if the relevant root were *d* = *d*_−_, 0 ≤ *V_G_* = *V_g_* + *d*_−_ < 0, a contradiction. So the relevant root is in fact

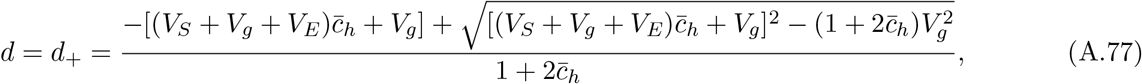

from which

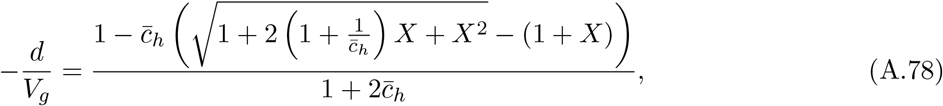

where 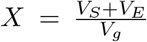. Since, in the absence of selection, *V_G_* = *V_g_*, Eq. (A.78) gives the proportionate reduction in the additive genetic variance due to selection.

From Eq. (A.72), 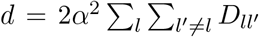, and, since 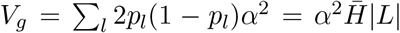, with |*L*| the number of loci and 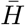 the average heterozygosity across them, we have

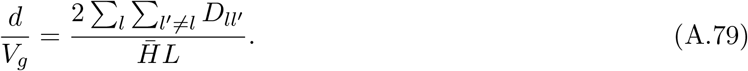

In a population association study performed at locus *l*, the effect size estimate is

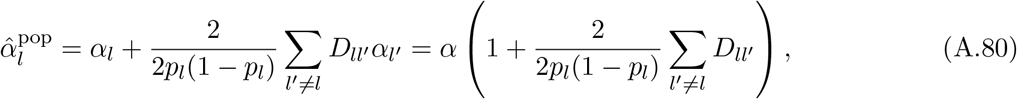

so that the proportionate error is

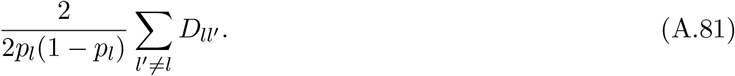

The mean proportionate error across loci is therefore

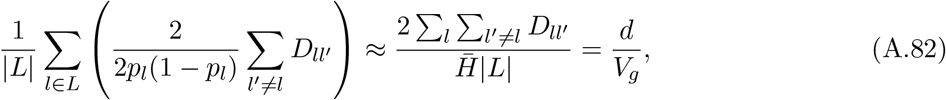

from Eq. (A.79), and assuming that the heterozygosities do not vary much across loci. That is, the average proportionate bias to effect size estimation that stabilizing selection induces is approximately equal to the proportionate reduction in the additive genetic variance, which is given in general form by Eq. (A.78).

In a within-family association study performed at locus *l*, the effect size estimate is

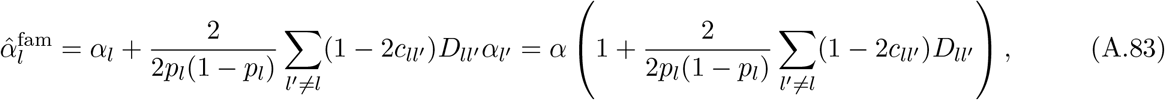

so that the proportionate error is

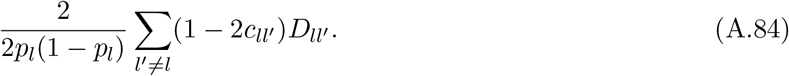

The mean proportionate error across loci is therefore

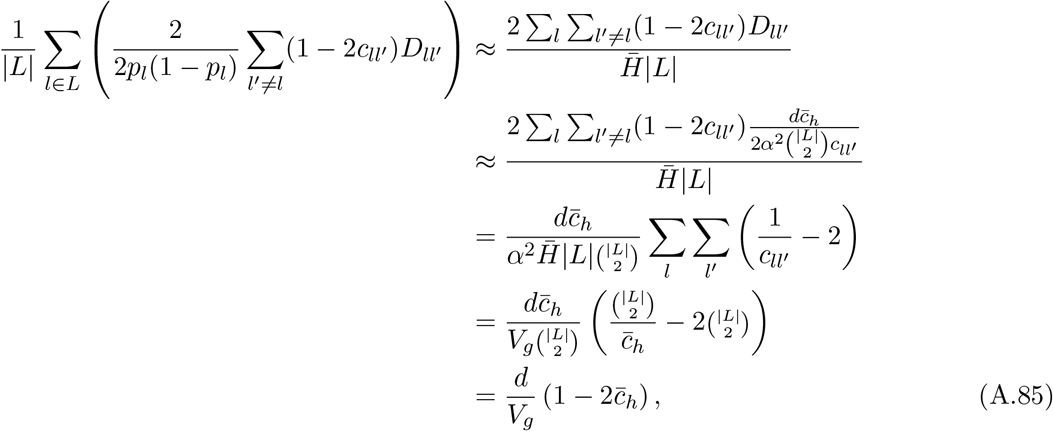

where we have used Eq. (A.74) in the second line. Therefore, the mean error in the within-family GWAS is smaller in magnitude than that in a population GWAS by a factor 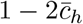.

If ~1,000 loci underlie variation in the trait (and all contribute approximately the same variation), 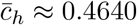 in humans (see Methods), and so the average bias that stabilizing selection induces in within-family GWASs will be about 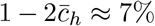 that in population GWASs. If ~10,000 loci underlie variation in the trait, 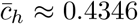, and so the bias in within-family GWASs will be about 13% that in population GWASs.

The calculations above give the average proportionate bias to GWAS estimates in terms of the basic parameters of the model, *V_g_, V_E_, V_S_*, and 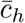. Often, however, not all of these parameters will be measurable. For example, human height appears to be under stabilizing selection (Sanjak et al. 2018), is highly heritable, and this heritability is believed to be underlain largely by *direct* genetic effects (Lee et al. 2018). However, it is difficult to directly measure the genic variance in height *V_g_* because not all causal loci will be assayed in association studies—and, moreover, even if they were, effect size estimation at these causal loci would be biased by the genetic confounds that we have studied in this paper. However, the phenotypic variance in height *V_P_* can obviously be measured, and the heritability of height *h*^2^ can also be measured using classical methods rather than effect size estimation in association studies. The strength of stabilizing selection on height can also be measured (Sanjak et al. 2018). From *V_P_* and *h*^2^, the additive genetic variance *V_G_* can be estimated (*V_G_* = *h*^2^*V_P_*).

This example suggests that, in many applications, it might be useful to be able to estimate the equilibrium value of *d* using *V_G_* (or *V_P_*), *V_E_, V_S_*, and 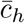, even though *V_G_* (and *V_P_*), in the model we have considered, is a state variable influenced by the state variable of primary interest, *d*. This is straight-forward: returing to our use of a star superscript to denote equilibrium values, if we treat *V_G_* and *V_P_* as their equilibrium values 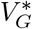 and 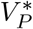, Eq. (A.71) can be estimated directly, and also simplifies to

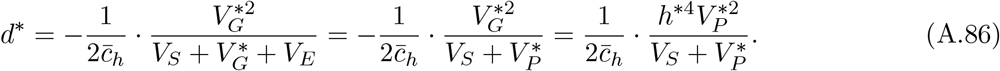

The proportionate bias in a population GWAS, given by Eq. (A.82), can similarly be estimated from *h*^2^, *V_P_, V_S_*, and 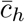, by first observing that

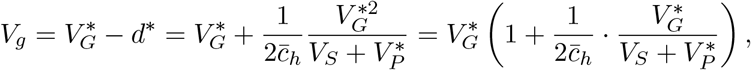

so that Eq. (A.82) can be written

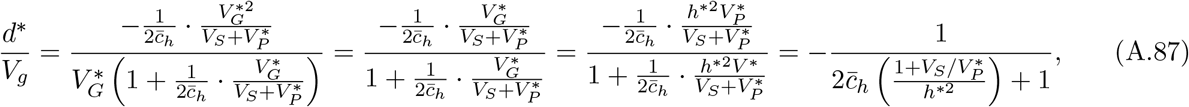

which reveals that the proportionate bias depends only on 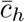, *h*\*^2^ and the scaled inverse strength of selection, 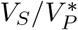.

From Eq. (A.85), the proportionate bias in a within-family GWAS is then approximately

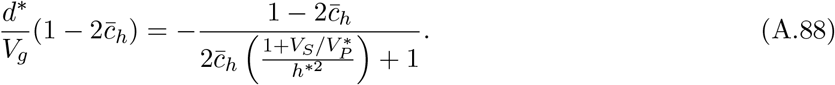

##### Stabilizing selection attenuates estimates of the strength of assortative mating based on cross-chromosome PGS correlations

Recently, the strength of assortative mating has been estimated based on measurement of the correlation of polygenic scores across distinct sets of chromosomes (e.g., Yengo et al. 2018; Yamamoto et al. 2023). Were assortative mating acting in isolation, such correlations would be due entirely to the positive cis- and trans-LDs among same-effect alleles created by assortative mating. Since stabilizing selection, acting in isolation, generates negative cis-LDs among same-effect alleles, it will attenuate the positive cis-LDs generated by assortative mating, and therefore reduce the correlation in PGSs among distinct sets of chromosomes, leading to underestimates of the strength of assortative mating if this effect is not taken into account.

To quantify this attenuation, we first calculate the strength of (positive) cross-chromosome LDs expected under assortative mating alone; then we calculate the strength of (negative) cross-chromosome LDs expected under stabilizing selection alone; then, assuming these LDs to be generated independently of one another—so that the LDs generated under the joint action of assortative mating and stabilizing selection are the sums of the LDs expected under these forces alone—we calculate how much stabilizing selection attenuates the correlation in PGSs across distinct sets of chromosomes.

##### Cross-chromosome correlations in PGSs

The number of autosomes in the haploid set is *n* (= 22 in humans). Label the set of loci on chromosome *k* that contribute variation to our trait of interest *L_k_*; the overall set of loci underlying variation in the trait is *L* = {*L*_1_, *L*_2_,…, *L_k_*}. We divide the chromosomes into distinct sets *K*_1_ and *K*_2_ (e.g., *K*_1_ could be the set of odd numbered chromosomes and *K*_2_ the even). Let *L*^(1)^ and *L*^(2)^ be the sets of causal loci on the chromosomes in *K*_1_ and *K*_2_ respectively (i.e., 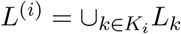).

Suppose that we have accurately estimated effect sizes at all loci *l* ∈ *L*. For each individual, we then calculate a polygenic score for *K*_1_ and for *K*_2_:

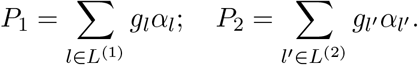

We are interested in the correlation in the population between *P*_1_ and *P*_2_, and in particular, how this correlation is affected by assortative mating and stabilizing selection for the focal trait. The correlation can be written

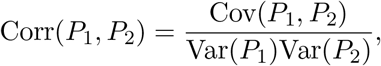

with

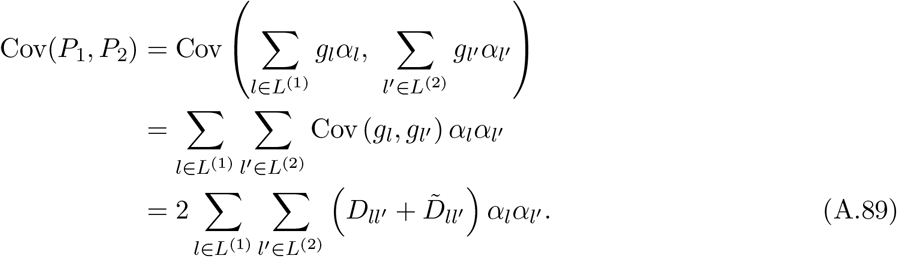

Since, to make progress in the case of stabilizing selection, we will assume effect sizes to be equal across loci, we make that assumption now, so that

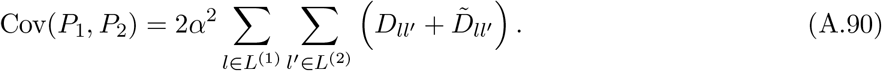

Since every pair of loci (*l,l*′) across *L*^(1)^ and *L*^(2)^ are by definition unlinked, under many processes (including assortative mating and stabilizing selection), the values of 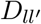 and 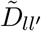 will not differ much in expectation across locus pairs, in equilibrium. Therefore, we may approximate 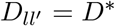 and 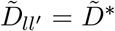 for all *l* ∈ *L*^(1)^ and *l*′ ∈ *L*^(2^, so that Eq. (A.90) simplifies further:

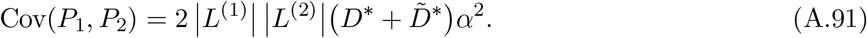

##### Assortative mating alone

Under assortative mating with equal effect sizes across loci, in equilibrium, LDs are approximately equal across locus pairs, regardless of the recombination rate between them; moreover, cis- and trans-LDs are equal (see above). Therefore, to calculate 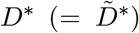, we simply apportion the total LD given by Eq. (A.40) among individual locus pairs:

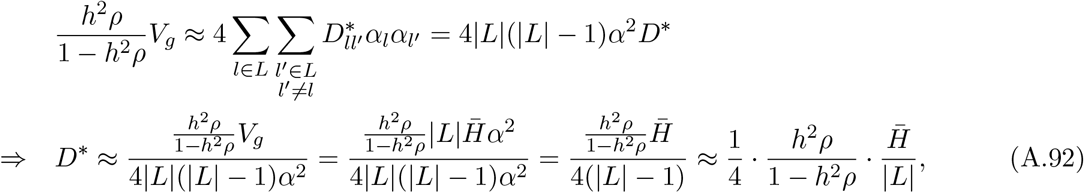

when |*L*| is large. Similarly,

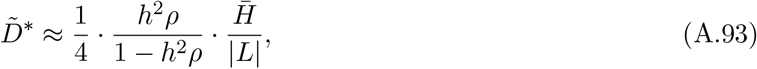

so that the overall contribution of assortative mating to the covariance in Eq. (A.91) is proportional to

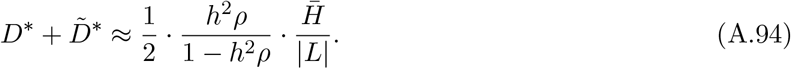

##### Stabilizing selection alone

Under stabilizing selection, the total amount of negative cis-LD is given by Eq. (A.87):

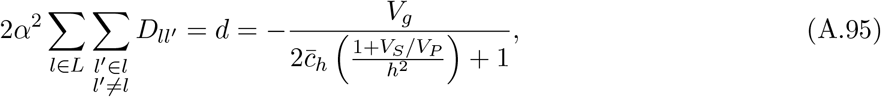

where we have dropped the equilibrium ‘*’ markers. This expression does not easily decompose into terms from individual locus pairs. However, if we assume that stabilizing selection is relatively weak 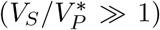 and that the recombination process is such that the harmonic mean recombination rate 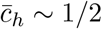 (as is the case in humans), Eq. (A.95) can be approximated by

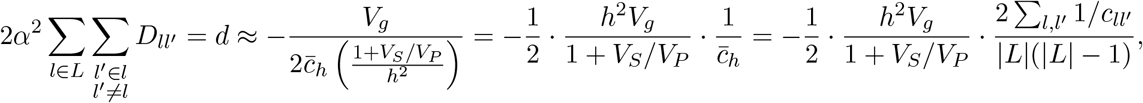

from which we infer that, in expectation,

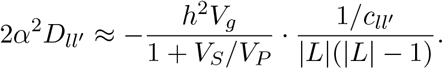

Therefore, for unlinked *l* and *l*′ 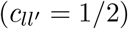, in expectation,

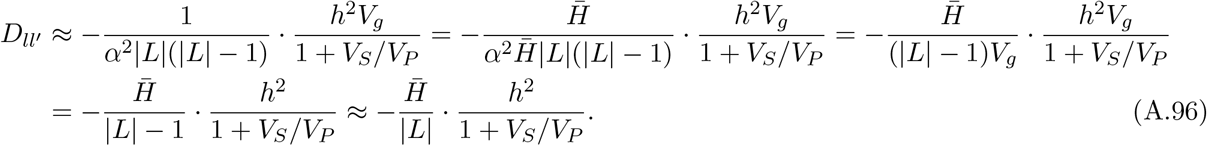

Stabilizing selection does not systematically generate trans-LD, so, in expectation, 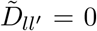. Therefore, under stabilizing selection alone, the contribution of an unlinked locus pair to the covariance in Eq. (A.91) is

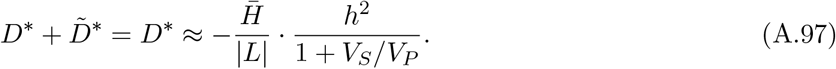

##### How much does stabilizing selection attenuate the signal of assortative mating?

Comparing Eqs. (A.94) and (A.97), we find that the proportionate attenuation of assortative mating’s effect (in isolation) by the action of stabilizing selection is

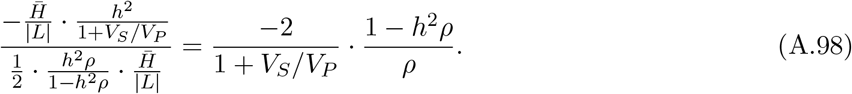

For example, in the case of human height (*h*^2^ ~ 0.8), the signal of assortative mating (strength *ρ* ~ 0.25) is attenuated by stabilizing selection (strength *V_S_/V_P_* ~ 30) by a proportionate amount of approximately 20%. That is, one might measure by other means (e.g., the phenotypic correlation among mates, together with an estimate of the heritability of height) that the strength of assortative mating is *ρ* = 0.25, but estimating this strength from cross-chromosome PGS correlations without accounting or correcting for stabilizing selection on height would yield 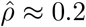, 20% smaller than the true value.

## A4 One-locus GxE

We study the phenotypic model in Eq. (22), with the phenotype of individual *i* in family *f* given by

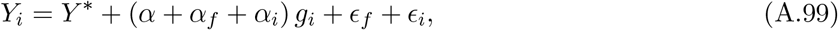

where, across the population, 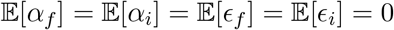, and *α_i_*, *ϵ_f_*, and *ϵ_i_* are all independent of *g_i_*.

### Sibling GWAS

Let *i* and *j* be siblings in family *f*, and define Δ*Y_f_* = *Y_i_* – *Y_j_*, Δ_*g_f_*_ = *g_i_* – *g_j_*, and Δ*ϵ_f_* = *ϵ_i_* – *ϵ_j_*. A sibling association study returns an effect size estimate

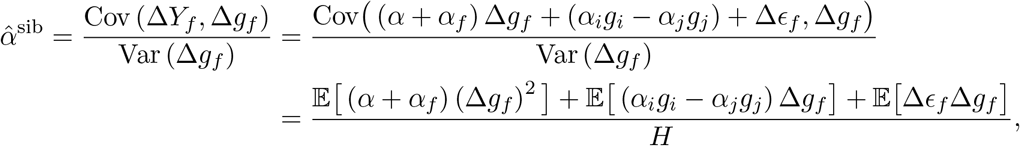

where *H* is the fraction of parents who are heterozygous at the focal locus. Since *α_i_*, *α_j_*, *ϵ_i_*, and *ϵ_j_* are genotype-independent perturbations, 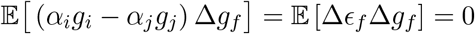, and so

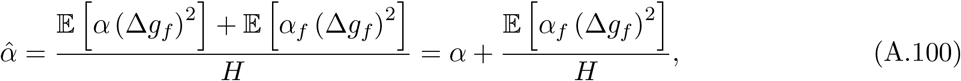

which deviates from *α* by an amount 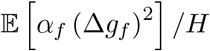.

Let 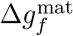 and 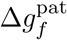 be the difference in the genotypes of the siblings in family *f* due to maternal and paternal transmission. Because of the independence of maternal and paternal transmission in a given family, the term additional to *α* in Eq. (A.100) can be split into 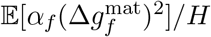 and 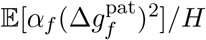, which we can analyze separately.

If the mother is heterozygous, then 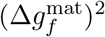 equals 1 with probability 1/2 and 0 with probability 1/2; if the mother is homozygous, then 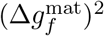 is 0. Therefore, denoting by *h*^m^ the event that the mother is heterozygous,

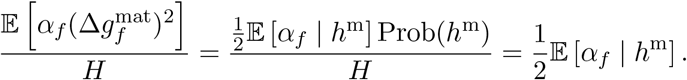

The same holds for paternal transmission, and so the deviation of the family-based estimate 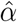 from *α* is

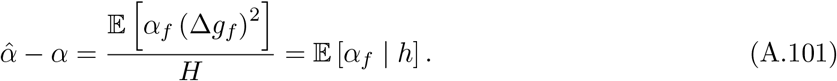

That is, quite intuitively, if the average G×E effect *α_f_* is different in the families of heterozygous parents than in the population as a whole, then limiting estimation to the offspring of heterozygous parents will be problematic.

### Population GWAS

Under the same one-locus model, a population association study returns an effect size estimate of

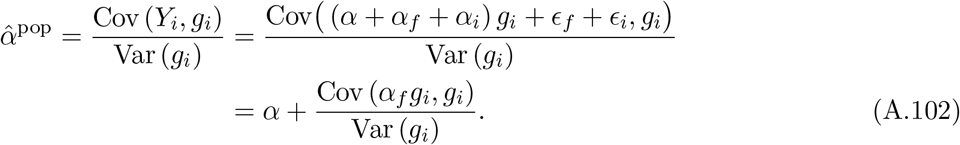

We can immediately see from Eq. (A.102) that if the family environments are randomized across genotypes, such that *α_f_* and *g_i_* are independent (implying Cov(*α_f_g_i_, g_i_*) = 0), then the population estimate will coincide with *α*.

To calculate the deviation of the population estimate from *α* in the general case, let *F* be the inbreeding coefficient at the locus. Then Var(*g_i_*) = 2*p*(1 – *p*)(1 + *F*), where *p* is the frequency of the focal variant, and the frequency of heterozygotes is *f*_1_ = 2*p*(1 – *p*)(1 – *F*) while the frequencies of the two homozygotes are *f*_0_ = (1 – *p*)^2^ + p(1 – *p*)*F* (zero focal alleles) and *f*_2_ = *p*^2^ + *p*(1 – *p*)*F* (two focal alleles). The covariance term in Eq. (A.102) can then be written

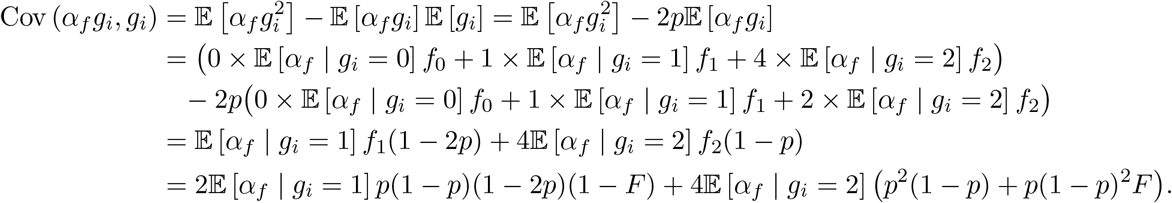

The deviation of the population-based estimate from *α* is therefore

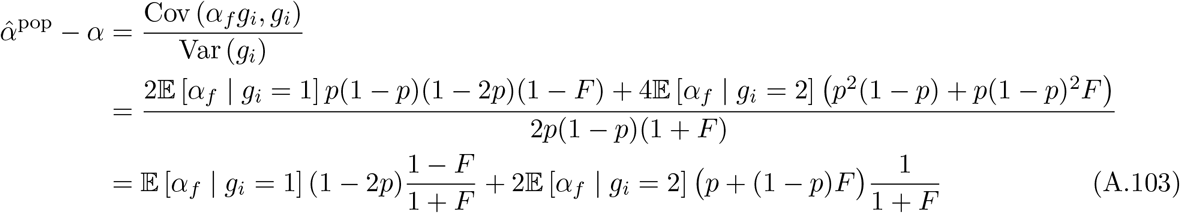

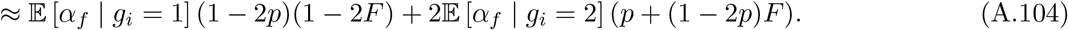

The approximation holds when *F* is small.

An interesting special case is where homozygotes for the focal allele and heterozygotes have the same distribution of environments, so that 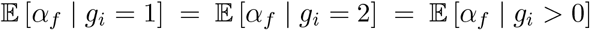. In this case, Eq. (A.103) simplifies to

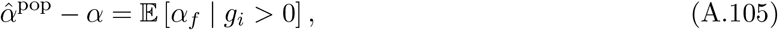

which reveals that, if individuals who carry the focal allele tend to experience different environments to individuals who do not carry the focal allele, then the population GWAS estimate will deviate from the average effect under true randomization, *α*. Moreover, in this case, if 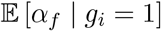 and 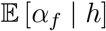 are the same—that is, if the mean environment of heterozygous offspring is the same as that for heterozygous parents—then the sibling and population-based effect size estimates are the same.

## Footnotes

1 By the Frisch-Waugh-Lovell theorem (Greene 2018, pg. 36), Eq. (3) is the estimate one obtains by first regressing both the trait and the focal-locus genotype on the PCs, collecting the residuals from these regressions (which can be thought of as trait and focal-locus genotype values stripped of whatever signal the PCs captured), and regressing the residuals from the trait regression on the residuals from the genotype regression.

## Notes

### Competing Interest Statement

The authors have declared no competing interest.

## References

Abdellaoui, A., Dolan, C. V., Verweij, K. J., and Nivard, M. G. (2022). Gene–environment correlations across geographic regions affect genome-wide association studies. Nature Genetics, 54(9):1345–1354.

Abecasis, G. R., Cardon, L. R., and Cookson, W. O. C. (2000). A general test of association for quantitative traits in nuclear families. American Journal of Human Genetics, 66(1):279–292.

Allison, D. B. (1997). Transmission-disequilibrium tests for quantitative traits. American Journal of Human Genetics, 60(3):676–690.

Atwell, S., Huang, Y. S., Vilhjálmsson, B. J., Willems, G., Horton, M., Li, Y., Meng, D., Platt, A., Tarone, A. M., Hu, T. T., et al. (2010). Genome-wide association study of 107 phenotypes in *Arabidopsis thaliana* inbred lines. Nature, 465(7298):627–631.

Barcellos, S. H., Carvalho, L. S., and Turley, P. (2018). Education can reduce health differences related to genetic risk of obesity. Proceedings of the National Academy of Sciences, 115(42):E9765–E9772.

Benonisdottir, S. and Kong, A. (2022). The genetics of participation: method and analysis. bioRxiv, doi: https://doi.org/10.1101/2022.02.11.480067.

Berg, J. J., Harpak, A., Sinnott-Armstrong, N., Joergensen, A. M., Mostafavi, H., Field, Y., Boyle, E. A., Zhang, X., Racimo, F., Pritchard, J. K., et al. (2019). Reduced signal for polygenic adaptation of height in UK Biobank. eLife, 8:e39725.

Border, R., Athanasiadis, G., Buil, A., Schork, A., Cai, N., Young, A., Werge, T., Flint, J., Kendler, K., Sankararaman, S., W, D. A., and A, Z. N. (2022a). Cross-trait assortative mating is widespread and inflates genetic correlation estimates. Science, 378(6621):754–761.

Border, R., O’Rourke, S., de Candia, T., Goddard, M. E., Visscher, P. M., Yengo, L., Jones, M., and Keller, M. C. (2022b). Assortative mating biases marker-based heritability estimators. Nature Communications, 13(1):1–10.

Brown, B. C., Price, A. L., Patsopoulos, N. A., and Zaitlen, N. (2016). Local joint testing improves power and identifies hidden heritability in association studies. Genetics, 203(3):1105–1116.

Brumpton, B., Sanderson, E., Heilbron, K., Hartwig, F. P., Harrison, S., Vie, G. Å., Cho, Y., Howe, L. D., Hughes, A., Boomsma, D. I., et al. (2020). Avoiding dynastic, assortative mating, and population stratification biases in Mendelian randomization through within-family analyses. Nature Communications, 11(1):3519.

Bulik-Sullivan, B. (2015). Relationship between LD score and Haseman-Elston regression. BioRxiv, doi: https://doi.org/10.1101/018283.

Bulik-Sullivan, B. K., Finucane, H. K., Anttila, V., Gusev, A., Day, F. R., Loh, P.-R., Duncan, L., Perry, J. R., Patterson, N., Robinson, E. B., et al. (2015a). An atlas of genetic correlations across human diseases and traits. Nature Genetics, 47(11):1236–1241.

Bulik-Sullivan, B. K., Loh, P.-R., Finucane, H. K., Ripke, S., Yang, J., Patterson, N., Daly, M. J., Price, A. L., and Neale, B. M. (2015b). LD Score regression distinguishes confounding from polygenicity in genome-wide association studies. Nature Genetics, 47(3):291–295.

Bulmer, M. G. (1971). The effect of selection on genetic variability. American Naturalist, 105(943):201–211.

Bulmer, M. G. (1974). Linkage disequilibrium and genetic variability. Genetics Research, 23(3):281–289.

Bürger, R. (2000). The Mathematical Theory of Selection, Recombination, and Mutation. Wiley, Chichester, UK.

Coop, G. and Przeworski, M. (2022a). Lottery, luck, or legacy. A review of “The Genetic Lottery: Why DNA matters for social equality”. Evolution, 76(4):846–853.

Coop, G. and Przeworski, M. (2022b). Luck, lottery, or legacy? The problem of confounding. A reply to Harden. Evolution, 76(10):2464–2468.

Crow, J. F. (1990). Mapping functions. Genetics, 125(4):669–671.

Crow, J. F. and Felsenstein, J. (1968). The effect of assortative mating on the genetic composition of a population. Eugenics Quarterly, 15(2):85–97.

Crow, J. F. and Kimura, M. (1970). An Introduction in Population Genetics Theory. Harper and Row, New York.

Demange, P. A., Hottenga, J. J., Abdellaoui, A., Eilertsen, E. M., Malanchini, M., Domingue, B. W., Armstrong-Carter, E., De Zeeuw, E. L., Rimfeld, K., Boomsma, D. I., et al. (2022). Estimating effects of parents’ cognitive and non-cognitive skills on offspring education using polygenic scores. Nature Communications, 13(1):4801.

Eaves, L. J., Pourcain, B. S., Smith, G. D., York, T. P., and Evans, D. M. (2014). Resolving the effects of maternal and offspring genotype on dyadic outcomes in genome wide complex trait analysis (“M-GCTA”). Behavior Genetics, 44:445–455.

Edelaar, P. and Bolnick, D. I. (2012). Non-random gene flow: an underappreciated force in evolution and ecology. Trends in Ecology & Evolution, 27(12):659–665.

Ewens, W. J. and Spielman, R. S. (1995). The transmission/disequilibrium test: history, subdivision, and admixture. American Journal of Human Genetics, 57(2):455–464.

Felsenstein, J. (1981). Continuous-genotype models and assortative mating. Theoretical Population Biology, 19(3):341–357.

Fisher, R. A. (1952). Statistical methods in genetics. Heredity, 6(1):1–12.

Fletcher, J., Wu, Y., Li, T., and Lu, Q. (2021). Interpreting polygenic score effects in sibling analysis. BioRxiv, doi: https://doi.org/10.1101/2021.07.16.452740.

Freeman, G. (1973). Statistical methods for the analysis of genotype-environment interactions. Heredity, 31(3):339–354.

Fry, A., Littlejohns, T. J., Sudlow, C., Doherty, N., Adamska, L., Sprosen, T., Collins, R., and Allen, N. E. (2017). Comparison of sociodemographic and health-related characteristics of UK Biobank participants with those of the general population. American Journal of Epidemiology, 186(9):1026–1034.

Gauderman, W. J., Mukherjee, B., Aschard, H., Hsu, L., Lewinger, J. P., Patel, C. J., Witte, J. S., Amos, C., Tai, C. G., Conti, D., et al. (2017). Update on the state of the science for analytical methods for gene-environment interactions. American Journal of Epidemiology, 186(7):762–770.

Greene, W. H. (2018). Econometric Analysis. Pearson, New York, 8th edition.

Haller, B. C. and Messer, P. W. (2019). SLiM 3: forward genetic simulations beyond the Wright–Fisher model. Molecular Biology and Evolution, 36(3):632–637.

Harpak, A. and Przeworski, M. (2021). The evolution of group differences in changing environments. PLoS Biology, 19(1):e3001072.

Haworth, S., Mitchell, R., Corbin, L., Wade, K. H., Dudding, T., Budu-Aggrey, A., Carslake, D., Hemani, G., Paternoster, L., Smith, G. D., et al. (2019). Apparent latent structure within the UK Biobank sample has implications for epidemiological analysis. Nature Communications, 10(1):1–9.

Hayes, B. and Goddard, M. (2010). Genome-wide association and genomic selection in animal breeding. Genome, 53(11):876–883.

Hayward, L. K. and Sella, G. (2022). Polygenic adaptation after a sudden change in environment. eLife, 11:e66697.

Horwitz, T. B. and Keller, M. C. (2022). A comprehensive meta-analysis of human assortative mating in 22 complex traits. bioRxiv, doi: https://doi.org/10.1101/2022.03.19.484997.

Howe, L. J., Nivard, M. G., Morris, T. T., Hansen, A. F., Rasheed, H., Cho, Y., Chittoor, G., Ahlskog, R., Lind, P. A., Palviainen, T., et al. (2022). Within-sibship genome-wide association analyses decrease bias in estimates of direct genetic effects. Nature Genetics, 54(5):581–592.

Josephs, E. B., Stinchcombe, J. R., and Wright, S. I. (2017). What can genome-wide association studies tell us about the evolutionary forces maintaining genetic variation for quantitative traits? New Phytologist, 214(1):21–33.

Kong, A., Thorleifsson, G., Frigge, M. L., Vilhjalmsson, B. J., Young, A. I., Thorgeirsson, T. E., Benonisdottir, S., Oddsson, A., Halldorsson, B. V., Masson, G., et al. (2018). The nature of nurture: Effects of parental genotypes. Science, 359(6374):424–428.

Kong, A., Thorleifsson, G., Gudbjartsson, D. F., Masson, G., Sigurdsson, A., Jonasdottir, A., Walters, G. B., Jonasdottir, A., Gylfason, A., Kristinsson, K. T., et al. (2010). Fine-scale recombination rate differences between sexes, populations and individuals. Nature, 467(7319):1099–1103.

Lander, E. S. and Schork, N. J. (1994). Genetic dissection of complex traits. Science, 265(5181):2037–2048.

Lee, H. and Lee, M. H. (2023a). Disentangling linkage and population structure in association mapping. https://github.com/hanbin973/hanbin973.github.io/raw/master/_data/LeeAndLee2023a.pdf.

Lee, H. and Lee, M. H. (2023b). Theoretical interpretation of genetic studies in admixed populations. https://github.com/hanbin973/hanbin973.github.io/raw/master/_data/LeeAndLee2023b.pdf.

Lee, J. J., Wedow, R., Okbay, A., Kong, E., Maghzian, O., Zacher, M., Nguyen-Viet, T. A., Bowers, P., Sidorenko, J., Karlsson Linnér, R., et al. (2018). Gene discovery and polygenic prediction from a genome-wide association study of educational attainment in 1.1 million individuals. Nature Genetics, 50(8):1112–1121.

Li, A., Liu, S., Bakshi, A., Jiang, L., Chen, W., Zheng, Z., Sullivan, P. F., Visscher, P. M., Wray, N. R., Yang, J., et al. (2023). mBAT-combo: a more powerful test to detect gene-trait associations from GWAS data. American Journal of Human Genetics, 110(1):30–43.

Marchini, J., Donnelly, P., and Cardon, L. R. (2005). Genome-wide strategies for detecting multiple loci that influence complex diseases. Nature Genetics, 37(4):413–417.

Martin, A. R., Gignoux, C. R., Walters, R. K., Wojcik, G. L., Neale, B. M., Gravel, S., Daly, M. J., Bustamante, C. D., and Kenny, E. E. (2017). Human demographic history impacts genetic risk prediction across diverse populations. American Journal of Human Genetics, 100(4):635–649.

Morris, T. T., Davies, N. M., Hemani, G., and Smith, G. D. (2020). Population phenomena inflate genetic associations of complex social traits. Science Advances, 6(16):eaay0328.

Mostafavi, H., Harpak, A., Agarwal, I., Conley, D., Pritchard, J. K., and Przeworski, M. (2020). Variable prediction accuracy of polygenic scores within an ancestry group. eLife, 9:e48376.

Nei, M. and Li, W.-H. (1973). Linkage disequilibrium in subdivided populations. Genetics, 75(1):213–219.

Nivard, M., Belsky, D., Harden, K. P., Baier, T., Ystrom, E., and Lyngstad, T. H. (2022). Neither nature nor nurture: Using extended pedigree data to elucidate the origins of indirect genetic effects on offspring educational outcomes. PsyArXiv, doi: https://doi.org/10.31234/osf.io/bhpm5.

Okbay, A., Wu, Y., Wang, N., Jayashankar, H., Bennett, M., Nehzati, S. M., Sidorenko, J., Kweon, H., Goldman, G., Gjorgjieva, T., et al. (2022). Polygenic prediction of educational attainment within and between families from genome-wide association analyses in 3 million individuals. Nature Genetics, 54(4):437–449.

Patel, R. A., Musharoff, S. A., Spence, J. P., Pimentel, H., Tcheandjieu, C., Mostafavi, H., Sinnott-Armstrong, N., Clarke, S. L., Smith, C. J., Durda, P. P., et al. (2022). Genetic interactions drive heterogeneity in causal variant effect sizes for gene expression and complex traits. American Journal of Human Genetics, 109(7):1286–1297.

Peiffer, J. A., Romay, M. C., Gore, M. A., Flint-Garcia, S. A., Zhang, Z., Millard, M. J., Gardner, C. A., McMullen, M. D., Holland, J. B., Bradbury, P. J., et al. (2014). The genetic architecture of maize height. Genetics, 196(4):1337–1356.

Pfaff, C. L., Parra, E. J., Bonilla, C., Hiester, K., McKeigue, P. M., Kamboh, M. I., Hutchinson, R. G., Ferrell, R. E., Boerwinkle, E., and Shriver, M. D. (2001). Population structure in admixed populations: effect of admixture dynamics on the pattern of linkage disequilibrium. American Journal of Human Genetics, 68(1):198–207.

Pirastu, N., Cordioli, M., Nandakumar, P., Mignogna, G., Abdellaoui, A., Hollis, B., Kanai, M., Rajagopal, V. M., Parolo, P. D. B., Baya, N., et al. (2021). Genetic analyses identify widespread sex-differential participation bias. Nature Genetics, 53(5):663–671.

Platt, A., Vilhjálmsson, B. J., and Nordborg, M. (2010). Conditions under which genome-wide association studies will be positively misleading. Genetics, 186(3):1045–1052.

Price, A. L., Patterson, N. J., Plenge, R. M., Weinblatt, M. E., Shadick, N. A., and Reich, D. (2006). Principal components analysis corrects for stratification in genome-wide association studies. Nature Genetics, 38(8):904–909.

Price, A. L., Zaitlen, N. A., Reich, D., and Patterson, N. (2010). New approaches to population stratification in genome-wide association studies. Nature Reviews Genetics, 11(7):459–463.

Pritchard, J. K. and Przeworski, M. (2001). Linkage disequilibrium in humans: models and data. American Journal of Human Genetics, 69(1):1–14.

Pritchard, J. K. and Rosenberg, N. A. (1999). Use of unlinked genetic markers to detect population stratification in association studies. American Journal of Human Genetics, 65(1):220–228.

Pritchard, J. K., Stephens, M., Rosenberg, N. A., and Donnelly, P. (2000). Association mapping in structured populations. American Journal of Human Genetics, 67(1):170–181.

Rosenberg, N. A. and Nordborg, M. (2006). A general population-genetic model for the production by population structure of spurious genotype–phenotype associations in discrete, admixed or spatially distributed populations. Genetics, 173(3):1665–1678.

Sanjak, J. S., Sidorenko, J., Robinson, M. R., Thornton, K. R., and Visscher, P. M. (2018). Evidence of directional and stabilizing selection in contemporary humans. Proceedings of the National Academy of Sciences, 115(1):151–156.

Sella, G. and Barton, N. H. (2019). Thinking about the evolution of complex traits in the era of genome-wide association studies. Annual Review of Genomics and Human Genetics, 20:461–493.

Selzam, S., Ritchie, S. J., Pingault, J.-B., Reynolds, C. A., O’Reilly, P. F., and Plomin, R. (2019). Comparing within-and between-family polygenic score prediction. American Journal of Human Genetics, 105(2):351–363.

Shen, H. and Feldman, M. W. (2020). Genetic nurturing, missing heritability, and causal analysis in genetic statistics. Proceedings of the National Academy of Sciences, 117(41):25646–25654.

Simons, Y. B., Mostafavi, H., Smith, C. J., Pritchard, J. K., and Sella, G. (2022). Simple scaling laws control the genetic architectures of human complex traits. bioRxiv, doi: https://doi.org/10.1101/2022.10.04.509926.

Słoczyński, T. (2022). Interpreting OLS estimands when treatment effects are heterogeneous: Smaller groups get larger weights. Review of Economics and Statistics, 104(3):501–509.

Sohail, M., Maier, R. M., Ganna, A., Bloemendal, A., Martin, A. R., Turchin, M. C., Chiang, C. W., Hirschhorn, J., Daly, M. J., Patterson, N., et al. (2019). Polygenic adaptation on height is overestimated due to uncorrected stratification in genome-wide association studies. eLife, 8:e39702.

Spielman, R. S., McGinnis, R. E., and Ewens, W. J. (1993). Transmission test for linkage disequilibrium: the insulin gene region and insulin-dependent diabetes mellitus (IDDM). American Journal of Human Genetics, 52(3):506–516.

Stulp, G., Simons, M. J. P., Grasman, S., and Pollet, T. V. (2017). Assortative mating for human height: A meta-analysis. American Journal of Human Biology, 29(1):e22917.

Trejo, S. and Domingue, B. W. (2018). Genetic nature or genetic nurture? Introducing social genetic parameters to quantify bias in polygenic score analyses. Biodemography and Social Biology, 64(3-4):187–215.

Tropf, F. C., Lee, S. H., Verweij, R. M., Stulp, G., Van Der Most, P. J., De Vlaming, R., Bakshi, A., Briley, D. A., Rahal, C., Hellpap, R., et al. (2017). Hidden heritability due to heterogeneity across seven populations. Nature Human Behaviour, 1(10):757–765.

Tyrrell, J., Zheng, J., Beaumont, R., Hinton, K., Richardson, T. G., Wood, A. R., Davey Smith, G., Frayling, T. M., and Tilling, K. (2021). Genetic predictors of participation in optional components of UK Biobank. Nature Communications, 12(1):886.

Ulizzi, L. and Terrenato, L. (1987). Natural selection associated with birth weight v. the secular relaxation of the stabilizing component. Annals of Human Genetics, 51(3):205–210.

Veller, C., Muralidhar, P., and Haig, D. (2020). On the logic of Fisherian sexual selection. Evolution, 74(7):1234–1245.

Vilhjálmsson, B. J. and Nordborg, M. (2013). The nature of confounding in genome-wide association studies. Nature Reviews Genetics, 14(1):1–2.

Visscher, P. M., Medland, S. E., Ferreira, M. A. R., Morley, K. I., Zhu, G., Cornes, B. K., Montgomery, G. W., and Martin, N. G. (2006). Assumption-free estimation of heritability from genome-wide identity-by-descent sharing between full siblings. PLoS Genetics, 2(3):e41.

Weiner, D. J., Wigdor, E. M., Ripke, S., Walters, R. K., Kosmicki, J. A., Grove, J., Samocha, K. E., Goldstein, J. I., Okbay, A., Bybjerg-Grauholm, J., et al. (2017). Polygenic transmission disequilibrium confirms that common and rare variation act additively to create risk for autism spectrum disorders. Nature Genetics, 49(7):978–985.

Weir, B. S. (2008). Linkage disequilibrium and association mapping. Annual Review of Genomics and Human Genetics, 9(1):129–142.

Wolf, J. B., Brodie III, E. D., Cheverud, J. M., Moore, A. J., and Wade, M. J. (1998). Evolutionary consequences of indirect genetic effects. Trends in Ecology & Evolution, 13(2):64–69.

Wright, S. (1921). Systems of mating. III. Assortative mating based on somatic resemblance. Genetics, 6(2):144–161.

Yair, S. and Coop, G. (2022). Population differentiation of polygenic score predictions under stabilizing selection. Philosophical Transactions of the Royal Society B, 377(1852):20200416.

Yamamoto, K., Sonehara, K., Namba, S., Konuma, T., Masuko, H., Miyawaki, S., Kamatani, Y., Hizawa, N., Ozono, K., Yengo, L., et al. (2023). Genetic footprints of assortative mating in the Japanese population. Nature Human Behaviour, 7(1):65–73.

Yang, J., Zaitlen, N. A., Goddard, M. E., Visscher, P. M., and Price, A. L. (2014). Advantages and pitfalls in the application of mixed-model association methods. Nature Genetics, 46(2):100–106.

Yengo, L., Robinson, M. R., Keller, M. C., Kemper, K. E., Yang, Y., Trzaskowski, M., Gratten, J., Turley, P., Cesarini, D., Benjamin, D. J., et al. (2018). Imprint of assortative mating on the human genome. Nature Human Behaviour, 2(12):948–954.

Young, A. I., Benonisdottir, S., Przeworski, M., and Kong, A. (2019). Deconstructing the sources of genotype-phenotype associations in humans. Science, 365(6460):1396–1400.

Young, A. I., Frigge, M. L., Gudbjartsson, D. F., Thorleifsson, G., Bjornsdottir, G., Sulem, P., Masson, G., Thorsteinsdottir, U., Stefansson, K., and Kong, A. (2018a). Relatedness disequilibrium regression estimates heritability without environmental bias. Nature Genetics, 50(9):1304–1310.

Young, A. I., Nehzati, S. M., Benonisdottir, S., Okbay, A., Jayashankar, H., Chanwook, L., Cesarini, D., Benjamin, D. J., Turley, P., and Kong, A. (2022). Mendelian imputation of parental genotypes improves estimates of direct genetic effects. Nature Genetics, 54:897–905.

Young, A. I., Wauthier, F. L., and Donnelly, P. (2018b). Identifying loci affecting trait variability and detecting interactions in genome-wide association studies. Nature Genetics, 50(11):1608–1614.

Zaidi, A. A. and Mathieson, I. (2020). Demographic history mediates the effect of stratification on polygenic scores. eLife, 9:e61548.

Zaitlen, N., Huntsman, S., Hu, D., Spear, M., Eng, C., Oh, S. S., White, M. J., Mak, A., Davis, A., Meade, K., et al. (2017). The effects of migration and assortative mating on admixture linkage disequilibrium. Genetics, 205(1):375–383.

Zaitlen, N., Pasaniuc, B., Sankararaman, S., Bhatia, G., Zhang, J., Gusev, A., Young, T., Tandon, A., Pollack, S., Vilhjálmsson, B. J., et al. (2014). Leveraging population admixture to characterize the heritability of complex traits. Nature Genetics, 46(12):1356–1362.

Zhu, C., Ming, M. J., Cole, J. M., Kirkpatrick, M., and Harpak, A. (2022). Amplification is the primary mode of gene-by-sex interaction in complex human traits. bioRxiv, doi: https://doi.org/10.1101/2022.05.06.490973.

